# Evolution of Larval Segment Position across 12 *Drosophila* Species

**DOI:** 10.1101/653121

**Authors:** Gizem Kalay, Joel Atallah, Noemie C. Sierra, Austin M. Tang, Amanda E. Crofton, Mohan K. Murugesan, Sherri Wykoff-Clary, Susan E. Lott

**Affiliations:** The Department of Evolution and Ecology, University of California, Davis, One Shields Avenue, Davis, CA 95616; Earth and Planetary Sciences Department, University of California, Davis, One Shields Avenue, Davis, CA 95616

**Keywords:** segment patterning, *Drosophila*, evolution, robustness, larval stage

## Abstract

Many developmental traits that are critical to the survival of the organism are also robust. These robust traits are resistant to phenotypic change in the face of variation. This presents a challenge to evolution. In this paper, we asked whether and how a well-established robust trait, *Drosophila* segment patterning, changed over the evolutionary history of the genus. We compared segment position scaled to body length at the first-instar larval stage among 12 *Drosophila* species. We found that relative segment position has changed many times across the phylogeny. Changes were frequent, but primarily small in magnitude. Phylogenetic analysis demonstrated that rates of change in segment position are variable along the *Drosophila* phylogenetic tree, and that these changes can occur in short evolutionary timescales. Correlation between position shifts of segments decreased as the distance between two segments increased, suggesting local control of segment position. The posterior-most abdominal segment showed the highest magnitude of change on average, had the highest rate of evolution between species, and appeared to be evolving more independently as compared to the rest of the segments. This segment was exceptionally elongated in the cactophilic species in our dataset, raising questions as to whether this change may be adaptive.

## Introduction

Many developmental phenotypes are critical to the survival and fitness of the organism. These phenotypes are often observed to be robust, in that they produce a stereotyped outcome despite variation encountered during development (Wagner 2005; Félix and Wagner 2006; Masel and Siegal 2009; Siegal and Leu 2014). The variation experienced in ontogeny can come in a variety of forms, including stochastic, genetic, and environmental variation (Wagner 2005; Félix and Wagner 2006; Masel and Siegal 2009; Siegal and Leu 2014). Some robust phenotypes have specific mechanisms in place to ensure they are produced faithfully (Félix and Wagner 2006; Masel and Siegal 2009; Siegal and Leu 2014; Nijhout et al. 2017). However, these same robust phenotypes do evolve over periods of evolutionary time (Félix et al. 2000; Arthur and Chipman 2005; Lott et al. 2007; 2010; Fowlkes et al. 2011; Félix 2012; Combs and Fraser 2018). This poses a fundamental question: how do robust traits evolve if the phenotypic variation necessary to selection is suppressed?

One such developmental phenotype is segmentation along the head to tail (anterior-posterior) axis. Segmentation is the periodic repetition of anatomical structures. It is a shared feature of three big animal phyla, annelids, arthropods, and chordates (Davis and Patel 1999; Tautz 2004). In *Drosophila*, the foundation of segmentation is laid during the beginning of embryogenesis (St Johnston and Nüsslein-Volhard 1992), when development is under control of the maternal gene products, before the zygotic genome is activated. Maternal gene products that are located in the anterior and posterior parts of the fertilized egg set up a concentration gradient along the length of the embryo (St Johnston and Nüsslein-Volhard 1992; Surkova et al. 2018). These maternal gene products regulate one another, and also regulate genes expressed later in development by the zygote (St Johnston and Nüsslein-Volhard 1992; Surkova et al. 2018). This genetic network of regulators, which has been well-established through decades of critical study (Nüsslein-Volhard and Wieschaus 1980; Kornberg and Tabata 1993; Nasiadka et al. 2002; Nüsslein-Volhard et al. 2008; Clark 2017) precisely divide the embryo into progressively smaller subsections over embryonic development, until the correct number of body segments is reached. At the end of the embryonic stage, a first-instar larva with a highly organized segmented pattern of differentiated structures is produced.

As long as segmentation has been investigated, its fundamental role in *Drosophila* development has been clear, as defects in this process can be detrimental (or lethal) to the organism (Wieschaus and Nuesslein-Volhard 2016). Indeed, the lethality of homozygous mutations in segmentation genes was critical to their discovery in mutant screens (Wieschaus and Nuesslein-Volhard 2016). Subsequent generations of experiments using increasingly advanced methods have demonstrated that in addition to being critical, segmentation is a very precise process, with measurements of expression domains of segmentation genes being highly reproducible between embryos (Houchmandzadeh et al. 2002; Gregor et al. 2007; Surkova et al. 2008; Jaeger 2010; Petkova et al. 2014; Bentovim et al. 2017). Additionally, research has shown that segmentation can proceed correctly, while keeping its precision, in the face of substantial perturbations. For instance, in a seminal study launching the interface between the gradient of the maternal gene *bicoid (bcd)* and its downstream target *hunchback (hb)* as a model for understanding precision and scaling in developmental signaling, Houchmandzadeh et al. (2002), also demonstrated that the relative position of the Hb expression boundary (relative to embryo size) was robust to substantial genetic and environmental perturbations. This study tested genetic perturbations in the form of mutations in important maternal genes or absence of whole chromosomes, and found very little or no variation in the relative position of Hb expression (Houchmandzadeh et al. 2002). Moreover, the standard deviation of the Hb expression boundary did not show a significant increase with these drastic perturbations, indicating that these genetic changes did not decrease precision in the Hb boundary (Houchmandzadeh et al. 2002). There is also substantial evidence that segmentation proceeds precisely in the face of dramatic environmental perturbation. For example, when whole (Houchmandzadeh et al. 2002) or two halves (Lucchetta et al. 2005) of *D. melanogaster* embryos were raised at different temperature extremes after fertilization, inducing different developmental rates, expression domains were formed in the same relative position and with the same level of precision observed in control embryos, by the gap or pair rule stage of embryonic development, respectively (Houchmandzadeh et al. 2002; Lucchetta et al. 2005).

The above studies, along with many others, show that *Drosophila* segmentation is a complex trait that is robust to many forms of variation (Driever and Nüsslein-Volhard 1988; Houchmandzadeh et al. 2002; Lucchetta et al. 2005; Manu et al. 2009b). It is perhaps unsurprising then that with the exception of some Hawaiian *Drosophila* species (Spieth 1981), it is difficult to observe any gross differences in segment position, size and shape, scaled to full body size, between adults of different *Drosophila* species. However, over longer evolutionary time scales, segment patterning can vary substantially between arthropod species (Regier et al. 2010). At the early embryonic stage in *Drosophila*, previous studies have shown small quantitative differences in relative position of segmentation gene expression domains between both closely related (Lott et al. 2007; 2010; Fowlkes et al. 2011; Combs and Fraser 2018) and more distantly related species (Fowlkes et al. 2011; Wunderlich et al. 2019). Within a species, there is little evidence for variation in the relative position of segmentation gene boundaries, with a few exceptions. For example, between *D. melanogaster* lines with variable egg size, Lott et al. (2007) found no significant differences in relative position of *even-skipped* (*eve*) stripe boundaries. Even when lines with extremes of egg size were crossed to generate the full range of embryo lengths between the parental lines, relative segment position was invariable between different genotypes. However, between *D. melanogaster* lines that were subjected to strong artificial selection for egg size, Miles et al. (2010) found few small quantitative differences in *eve* patterning using three-dimensional imaging techniques. And, Jiang et al. (2015) was able to identify a line of *D. melanogaster* from the *Drosophila* Genetic Reference Panel (Mackay et al. 2012) with altered *even-skipped* pattern formation. It is not known, however, whether these changes persist beyond the embryonic stage, or if these trends appear beyond the limited number of species examined. So, while segment patterning is well documented to be robust, it has also been demonstrated to have some small quantitative level of variation between species, and in some circumstances, within species. Evolution of this robust trait does occur, and the presence of some variation within species suggests that evolution of small changes may be possible without causing catastrophic failure of patterning. Thus, we hypothesized that the evolution of segmentation may occur by relatively small shifts, rather than rare large leaps. This would require large sample sizes to detect.

In order to investigate the evolution of segment patterning systematically across a range of divergence times, we characterized segment position across 12 species of *Drosophila*. These 12 species spanned the evolutionary history of the genus (40-60 million years; (Russo et al. 1995; Obbard et al. 2012; Russo et al. 2013) and included three pairs of sister species. We measured position of each abdominal segment relative to body length, referred to throughout the manuscript as relative segment position, in the first-instar larvae (Fig. 1). Larval stage is the latest developmental stage where all segments are visible and easy to measure, and the use of this stage facilitated the measuring of more than a hundred larvae from each species. We found that relative segment positions at the larval stage have changed many times in the evolutionary history of *Drosophila*, and that these changes were mostly small in magnitude, with some larger changes. Most species-pair comparisons showed differences in the relative position of most abdominal segments. Most sister species were significantly different at every segment, however some of the most diverged species in our dataset showed no differences in their patterning. The magnitude of differences in relative position increased towards the posterior segments in most species, most strikingly in *D. persimilis* and *D. mojavensis*. Phylogenetic modelling showed that rate of segment position change was highly variable among the branches along the *Drosophila* phylogeny, and evolutionary rate changed even between closely related species. Results of a correlation analysis for position change between segments within species suggested local control of segment position, as correlations decrease with physical distance. Correlations of changes in the rate of evolution of segment position between branches of the phylogenetic tree also decreased with physical distance between segments. Overall, these results demonstrate that this complex and robust developmental trait does evolve, even over short timescales, and it does so primarily by small frequent steps with occasional large leaps. This may permit the evolution of novel patterns over long periods of evolutionary time, without compromising the integrity of this critical developmental phenotype.

**Figure 1.**
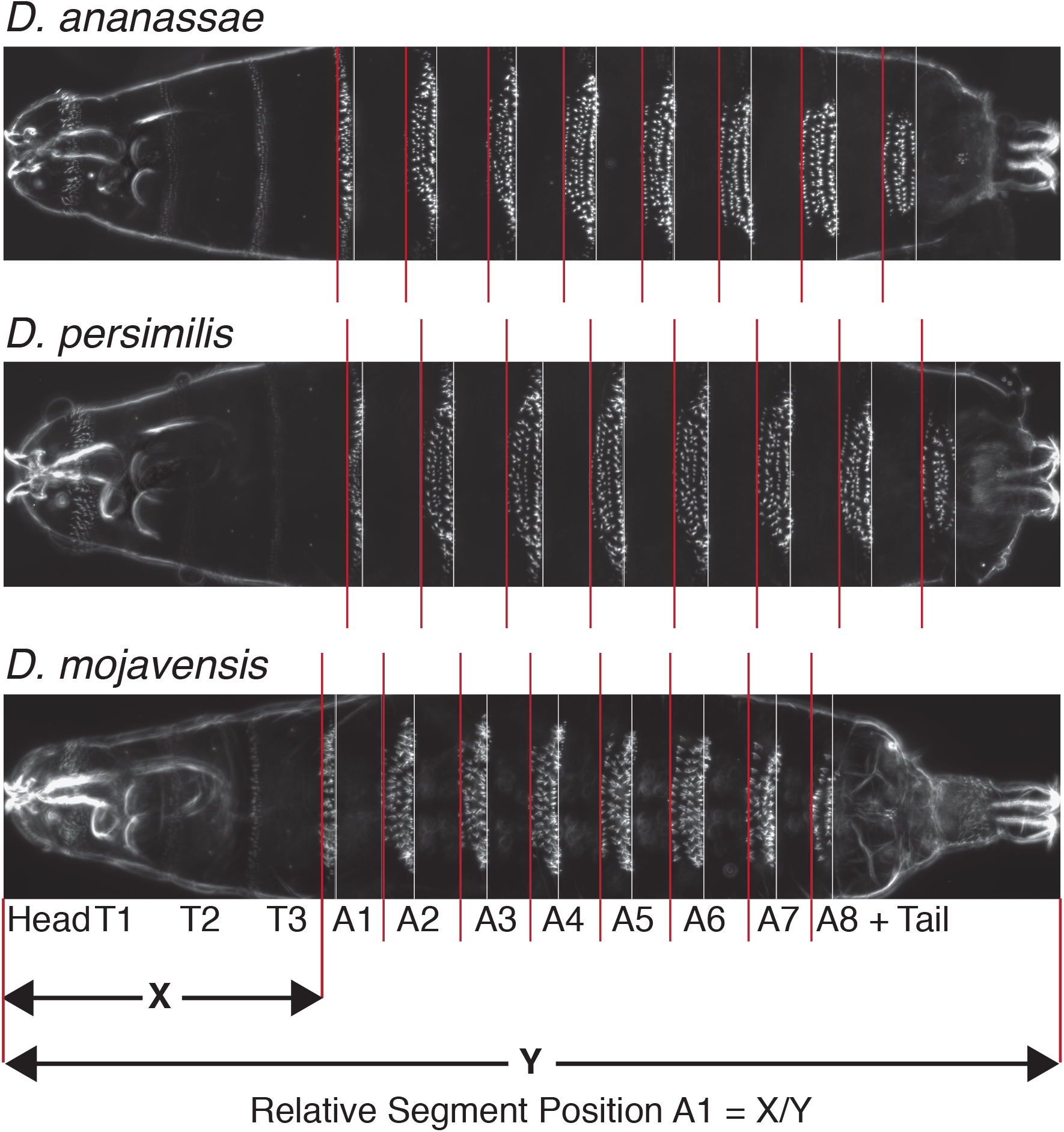
Example larval images and description of measurements. Example dark field images of first instar larvae are shown for three species. *D. ananassae* is presented as it is closest to the mean of all species for segment positions, while *D. persimilis* and *D. mojavensis* are exceptional, especially in the length of their most posterior segments. Red lines mark the anterior border and white lines mark the posterior border of denticle belts, which are rows of bristles on the ventral surface of the animal and are recognized by our image processing program. The anterior border of the denticle belt was used as a proxy for segment border. To measure the position of an abdominal segment (e.g. A1) relative to the body length (Y), the distance from the anterior border of the larvae to the anterior border of that segment (X) is divided by the total body size (Y). Region encompassing Head, T1, T2, T3 is referred to as “h+t” throughout the main text. T = Thoracic Segment, A = Abdominal Segment.

## Materials and Methods

### Species used

We took relative position measurements of abdominal segments from 12 *Drosophila* species spanning the evolutionary history of the genus. These species were *D. melanogaster*, *D. simulans*, *D. sechellia*, *D. yakuba*, *D. santomea*, *D. erecta*, *D. ananassae*, *D. pseudoobscura*, *D. persimilis*, *D. willistoni*, *D. mojavensis wriglei* and *D. virilis* (Fig. 2A). We used the sequenced lines from 11 of these species (Clark et al. 2007). For *D. santomea* we used stock #14021-0271.01 (The National *Drosophila* Species Stock Center).

**Figure 2.**
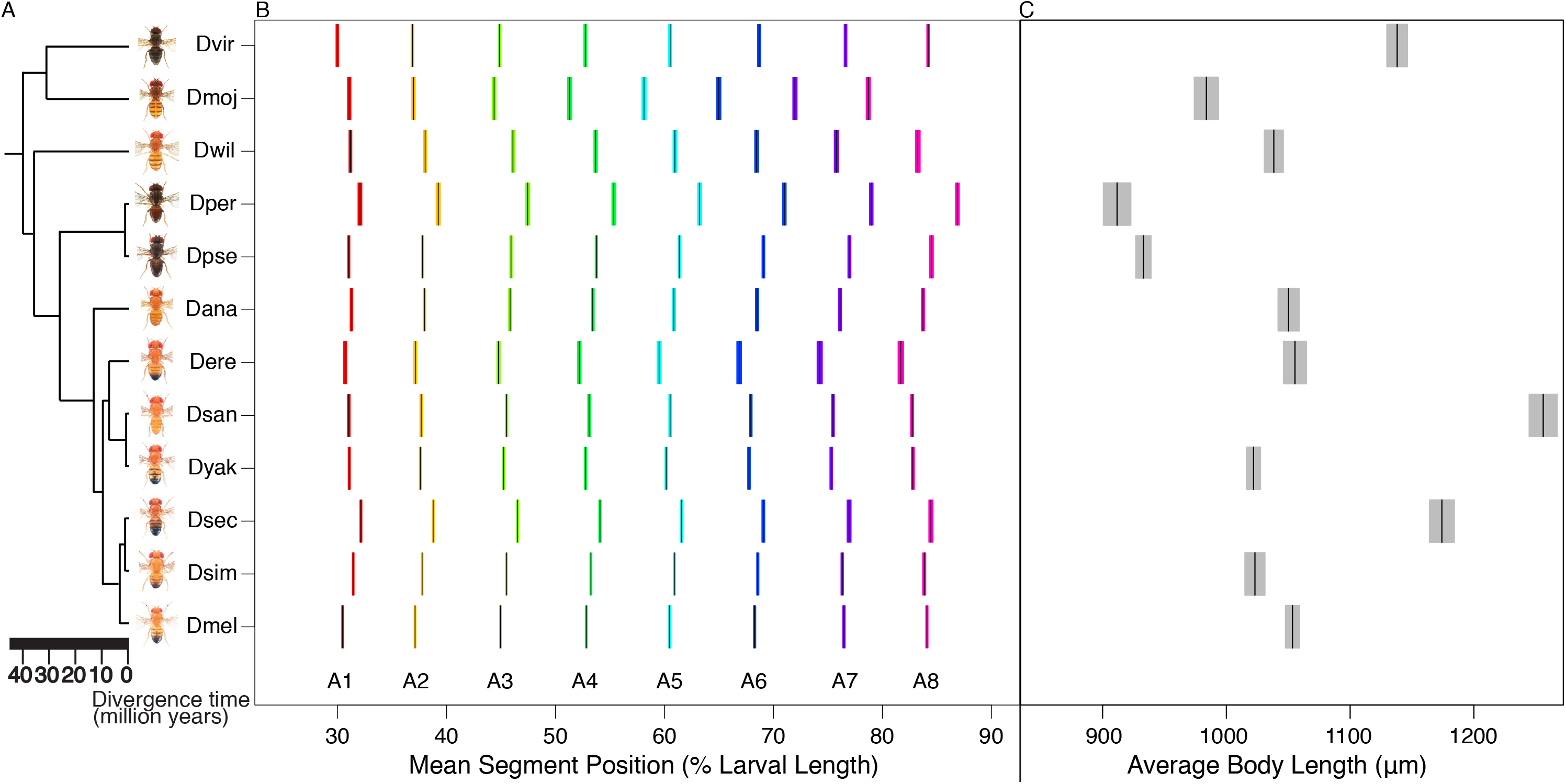
Relative segment position and body size is highly variable among 12 species of *Drosophila*. (A) 12 *Drosophila* species on an evolutionary tree. Adult male species photos from Nicolas Gompel were downloaded from FlyBase. (B) Each black bar represents the mean position in percent larval length for each segment in each species. The color spread on the two sides of each black bar is 95% confidence interval. Each segment is represented by a different color. x-axis is relative segment position in percent larval length. A = Abdominal Segment. (C) Each black bar represents the average first instar larval body length for each species. The gray shaded area around the black bars is 95% confidence interval. x-axis is average body length in micrometers.

### Fly husbandry and population size control

All flies were kept in plastic bottles in 20°C incubator with 60% humidity. Fly stocks were raised on standard cornmeal food. The amount of time to sexual maturation varied between species (Markow et al. 2008). Population control was conducted in a species-specific manner through controlling the number of sexually mature adults added in a given bottle, and how often the bottles were changed (Table S1). As a result, each bottle ended up with 40-50 pupae. Adult flies were discarded 14 days after they became fertile, to control for parental age.

### Control experiment on the effect of the larval mounting procedure

In order to address whether the larval mounting procedure has species-specific effects on measurements of relative segment position in first-instar *Drosophila* larvae, we examined relative segment position in *D. melanogaster*, *D. sechellia* and *D. virilis* at three different stages of the larval prep procedure: before heating, after heating, and after mounting (Fig. S1). As these three species are variable in their egg (Markow et al. 2008) and larval size (Fig. 2C) and have varying phylogenetic distances (Fig. 2A), we hypothesized that they might have a higher chance of being differentially affected by the larval preparation protocol. First, first-instar larvae that hatched in distilled water were temporarily immobilized using a 3-5 minute incubation on ice, put on a microscope slide with water and then imaged using a Zeiss AxioImager microscope in brightfield. These larvae were then put back in distilled water incubated at 60°C for 50 minutes and imaged in water a second time using a brightfield microscope. The larvae were then mounted in PVA Mounting Medium (BioQuip) (see next section). The larvae were then imaged a third time using dark field microscopy (see “Imaging” section). In the images of the iced (untreated control, the icing procedure slows larvae enough for imaging) and then the heated larva, each of the 2nd and 3rd thoracic, plus eight abdominal segments were marked by a node between bulges on the larval cuticle (Fig. S1). In the dark field images, segment borders were determined by the anterior border of each denticle belt. Denticle belts are rows of bristles on the ventral side of the larvae that are used for traction while crawling (Bejsovec 2013). The anterior border of each denticle belt is a proxy for the anterior border of each abdominal segment (Lohs-Schardin et al. 1979). Using the Image J (version 1.47t) “Line tool”, measurements were taken from the anterior end of the larva to the border of each segment. Segment position was then determined by dividing this value by the full length of the larvae (Fig. 1). Segment position data was analyzed using the following linear models (using the “lm” function, all terms were fixed), implemented in R (R Core Team 2019).

Relative segment position = µ + prep + species + prep x species + ε.

Relative segment position = µ + prep + species + segment + prep x species x segment + ε.

As a result, whether we included segment as a factor (p-value = 0.07 to 0.82) or not (p-value = 0.42 to 0.83) in our linear model formula, we did not find any significant species-specific effects of the preparation protocol on segment position measurements.

### Preparing Drosophila first-instar larvae for imaging

For each species, 20-50 newly emerged adults were obtained from a population-controlled bottle and put in an egg collection bottle with a cap containing glucose-agar food. For *D. sechellia*, cornmeal fly food was used in an egg lay cap with yeast sprinkled on it, as this prevents this species from withholding egg laying, a particular problem with this species (Markow et al. 2008). The bottle was then placed in 20°C incubator upside down. The next day eggs were collected from egg lay caps, put onto a mesh and thoroughly washed using distilled water to remove residual yeast and egg lay cap food. The eggs were then placed in a petri dish filled with distilled water. This petri dish was placed in 20°C until the eggs developed into first-instar larvae. In addition to the variability in the number of days necessary to reach sexual maturity, different species also varied in the number of days necessary for a fertilized egg to develop into first-instar larvae (Markow and O’Grady 2005). The petri dish with water and larvae was then placed in 60°C oven for ∼50 minutes (Table S1), which killed and straightened the larvae. These larvae were then mounted on standard glass slides using PVA Mounting Medium (BioQuip), standard coverslips and a dissection microscope. Larvae were oriented such that their ventral side was facing up, their posterior spiracles were protruding from their body and their left-right symmetry was protected. Once the slides were ready, each coverslip was sealed with clear nail polish. The slides were then incubated at 60°C overnight.

### Imaging

The slides were imaged at 40X objective using a Zeiss AxioImager microscope and a dark field filter. Using automated tiling, 64 high resolution images were taken for each larva, which were later stitched using the ZEN 2012 (blue edition) software. Each image was then exported to “tagged image format”. These images will be made available prior to publication.

### Image processing

Measurements for the position of each abdominal denticle belt were made using a custom Python script (the program will be made available prior to publication). This program rotated and positioned each larva horizontally, anterior to the left and posterior to the right, and cropped the image at the anterior, posterior and lateral borders. The program then marked the anterior and posterior borders of the abdominal denticle belts. It measured the distance from the anterior-most point of the larva to the anterior as well as posterior borders of each denticle belt, and from the anterior-most point of the larva to the posterior-most point of the larva (Fig. 1). Relative segment positions were calculated as the distance of the anterior denticle boundary from the anterior of the larva, divided by larval length (Fig. 1). The number of larval samples, from which segment position measurements were taken and used for data analysis, varied from 105 to 145 for each species (Table S1).

### Image editing

Some of the images were edited using ImageJ “Brush” tool to paint over bubbles around the larvae and to adjust brightness and contrast when necessary. We found both of these edits increased the number of successful runs by the custom image processing program and did not change the measurements taken from these images.

### Data analysis

The segment measurement data is available at https://doi.org/10.6084/m9.figshare.8170787. A linear model ANOVA was fitted to the data using R (R Core Team 2019) with the effects of species and segment and an interaction term. “lm” and “aov” functions were used to implement the following formulas. All terms were fixed.

Relative segment position. = µ + species + segment + species x segment + ε

This was followed by Tukey’s HSD (Honestly Significant Differences) (Steel et al. 1997) test to conduct pairwise comparisons of the relative position of each segment between species. Specifically, we used “HSD.test” function from the R package “agricolae” (De Mendiburu 2019). We replicated these results using t-tests between each species pair and then applying Bonferroni multiple test correction (“t.test” and “p.adjust” functions in R, respectively) for both multiple species and multiple segments. These analyses were done also for the posterior border and anterior-posterior width of denticle belts. For the details of “posterior border” and “width” analyses of denticle belts, see File S1.

Principal component analysis was performed in R, using the “prcomp” function, and mean centered positions of each segment as the data. To examine what PC1 represented in our data, we used Pearson correlation (“cor.test” function in R) to correlate PC1 with the relative positions of each segment. Using the same method, we also tested whether PC1 was correlated with larval length.

For correlation analysis between changes in relative segment position, we first calculated deviation from the between-species mean of the relative position of each segment in each species. We then used “cor” and “cor.test” functions in R to obtain Pearson correlation coefficients and the associated p-values.

### Phylogenetic Ancestral State Estimation

We performed phylogenetic analyses to infer the evolution of relative segment position across the 12 *Drosophila* species here. For this, we first inferred an ultrametric phylogeny published in Turelli et al. (2018) under three candidate relaxed molecular-clock models. For each of the resulting phylogenies, we assessed the fit of four candidate models that variously describe how rates of segment evolution vary across branches of the phylogeny. Finally, we jointly inferred the phylogeny, model of segment evolution, and the ancestral states for each segment under the preferred model of segment evolution. We summarized various aspects of the evolution of relative segment position from the resulting joint posterior probability distribution of ancestral states. Complete details of these analyses are in File S2.

## Results

### Larval segment allometry is highly variable across *Drosophila* species

To elucidate whether relative segment position has changed over *Drosophila* evolutionary history, we compared relative abdominal segment position (Fig. 1) at the first-instar larval stage between 12 different species (Fig. 2A). We found that there are many significant differences in relative segment position among species (Fig. 2B). Overall, in 21 out of 66 species-pair comparisons, all eight segments had a significant difference in their position. We note that species pairs do not represent independent observations due to the underlying phylogeny (we perform a full phylogenetic analysis below). For the majority of species-pair comparisons, the relative positions of five or more segments differed (Fig. S2). In fact, there were only two pairs of species compared (*D. simulans* - *D. ananassae* and *D. ananassae* - *D. willistoni*) where none of the eight abdominal segments had a significant difference in their relative positions (Table S2). Intriguingly, these species pairs are not closely related, with 15 and 32 million years of divergence, respectively. In all sister species comparisons, except for *D. yakuba* – *D. santomea*, the relative position of the majority of abdominal segments were different. On the other hand, between divergent species pairs, such as *D. melanogaster* and *D. virilis*, relative positions of only two segments were different (Table S2).

Interestingly, when considering all species-pair comparisons together, some species were responsible for a larger proportion of differences than others (Table S3). Out of the 410 segment position differences observed over all species-pair comparisons, differences with *D. persimilis* constituted 87 of these, the highest proportion of any species (∼21%). Differences with *D. sechellia*, *D. erecta* and *D. mojavensis* followed with 78 each (∼19%), whereas the number of differences with *D. ananassae* was the lowest with 56 (∼13.5%). This is consistent with the observation that relative segment positions of *D. mojavensis* and *D. persimilis*, followed by *D. erecta* and *D. sechellia*, have the highest total deviation from the species mean, whereas those of *D. ananassae* are closest to the species mean making it the “average” species (Table S3).

Next, we examined whether differences in segment position between species were toward the head (anterior) or tail (posterior) of the larva. To do this, we calculated mean positions of each segment across the 12 species and characterized whether a particular segment in a particular species was more anterior or more posterior relative to this mean (Fig. 3). For the majority of the species, the relative position of segments physically closest to each other differed in the same direction. When the direction of the difference changed, it was between segments that had the smallest (except for *D. willistoni*) magnitude of difference in a given species (Fig. 3).

**Figure 3.**
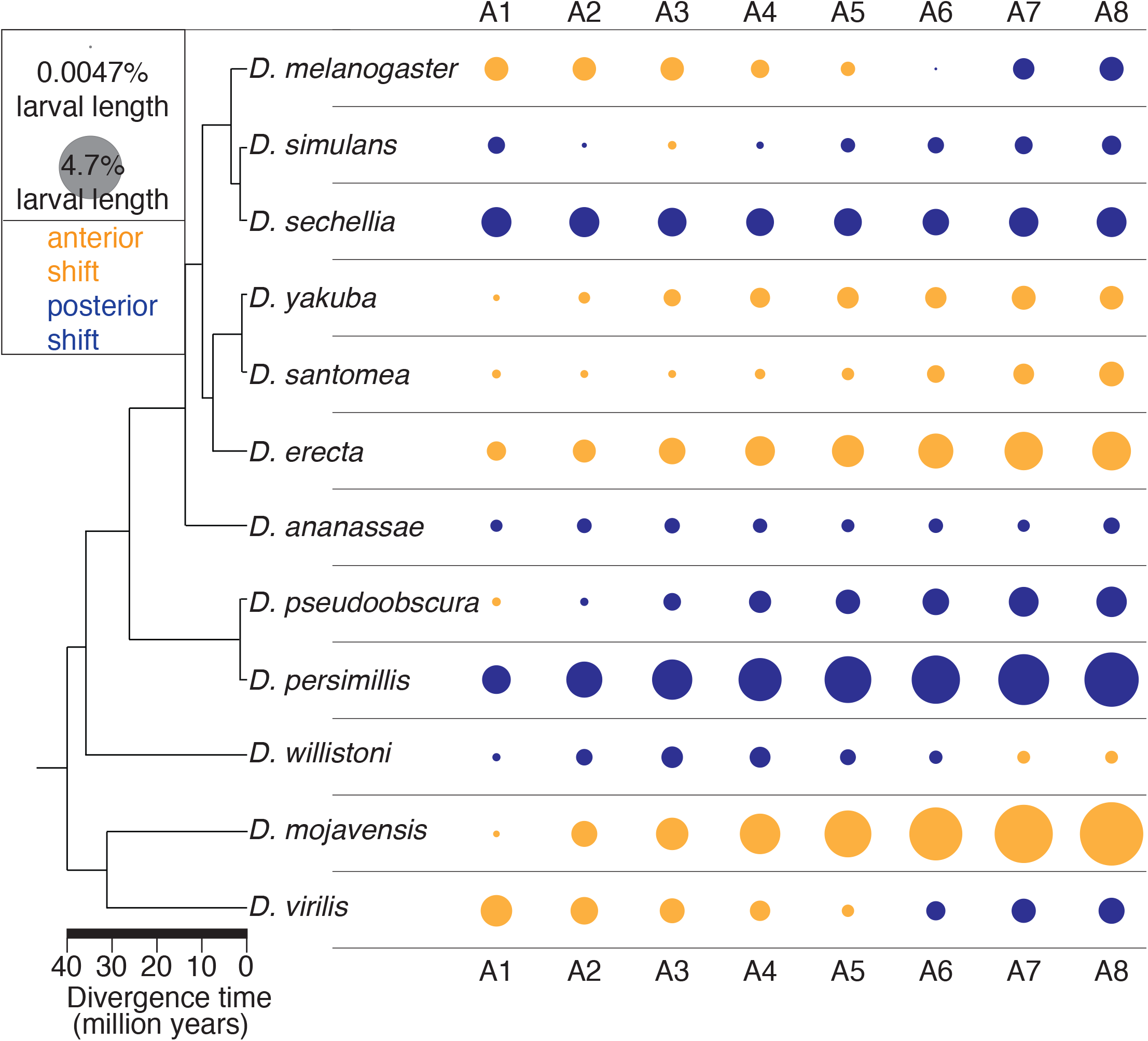
Coordinated direction of relative segment position changes in 12 *Drosophila* species. Neighboring segments tend to shift together in a particular direction, toward the anterior or posterior of the larva rather than shifting in opposite directions. Yellow indicates anterior, blue indicates posterior shift in relative position as compared to the mean position across all species for a given segment. The area of the circle is proportional to the size of the shift in relative segment position.

Notably, averaged over all species, deviation from the species mean in relative segment position increased from anterior to posterior of the larva, with the posterior-most segments having largest differences from the mean over all species. (Figs. 3, S3A). Principal component analysis (Fig. S3B) also demonstrates this pattern, with PC1 correlating highly with the more posterior segment positions, explaining 88% of the variance in the dataset. The trend of deviations in segment position increasing from anterior to the posterior of the larva was strongest for *D. mojavensis* and *D. persimilis*, (Figs. 3, S3C), but was nonetheless true in a number of other species as well (Fig. S3C, Table S3). Indeed, when segment position data for *D. mojavensis* and *D. persimilis* were excluded from the analysis, the trend still held (Fig. S3D).

In most species, magnitude of relative position change was highest for segment A8 (Table S3), which extends from the anterior boundary of A8 to the tail of the larvae (Fig. 1). We will refer to this segment as A8+tail. *D. mojavensis* and *D. persimilis* have the largest differences for segment A8: *D. mojavensis* has the largest magnitude of deviation from the species mean, with an anteriorly shifted segment A8 border and thus a much longer A8+tail segment; while *D. persimilis* has a posteriorly shifted segment A8 border and thus an exceptionally short A8+tail segment. (Figs. 1, 2B and S5C). While the magnitude of differences was highest for A8, the total number of differences are not increased for this segment. Significant differences in relative position observed in species-pair comparisons are highly similar in number for segments A3 through A8, with segments A1 and A2 showing a slightly lower number of significant differences in relative position (Fig. S4B, black line). Additionally, to determine if segment positioning in the posterior of the embryo is less precise (noisier) as compared to the anterior, we measured coefficient of variation for the position of each segment along the larva. The coefficient of variation, averaged across all species, does not vary considerably and is low across the length of the larva (∼0.03) (Fig. S6A). This suggests that while segment A8+tail has the largest magnitude differences in our dataset, its position is not more variable between species, nor is it positioned any less precisely than the other segments.

### Changes in the size of A8+tail segment are responsible for a significant portion of the shifts in relative segment position between species

Given that the relative position of segment A8+tail, as compared to the other segments, had the largest deviations from the species mean for most of the 12 species (Fig. S3, Table S3), we asked how much of the total variation in relative segment position is driven by A8+tail. In order to address this, we recalculated relative segment position in the absence of A8+tail (see File S3). To determine whether any changes in segment position we see was due to the removal of A8+tail region specifically, and not simply due to the removal of a terminal segment, we made a separate recalculation of relative segment position after removing the head and thoracic region (h+t) (Fig. 1). In both cases we determined the total number of significant differences in relative segment position between pairs of species (t-test with Bonferroni correction), and corrected this number relative to the total number of comparisons. We found that the corrected total number of significant position differences was reduced from 51.25 to 38.42 (∼25% decrease) when A8+tail was removed, but increased from 51.25 to 53.71 when h+t was removed (Fig. S4A, compare also Fig. 2B to Fig. S7A and B). This trend held even when segment position data for species with highest magnitude of posterior segment position differences, *D. mojavensis* and *D. persimilis*, were removed from calculations (from 31.63 down to 24, ∼24% decrease, when A8+tail was removed, but up to 32.71 when h+t was removed). This suggests that changes in the size of A8+tail drove a substantial portion of the differences in relative segment position between species. On the other hand, changes in the size of h+t appear to have masked some of the inter-species differences in relative position observed in the rest of the segments. These results are consistent with the finding that the average difference in relative position of A1, and hence the difference in the size of h+t, are the smallest in magnitude among all segments (Figs. S3A, S5B), while relative position of A8, hence the size of A8+tail, has the largest magnitude changes (Figs. S3A, S5C). They are also consistent with the finding that total number of significant position changes are lower for segment A1 than they are for A8 (Fig. S4B). For a more detailed analysis of “end removal” and its effects as well as differences in the size of h+t and A8+tail, see File S3.

### Changes in the length of larval body

As we had collected larval length measurements in order to calculate relative segment positions, we also compared whole body length at the first-instar larval stage among 12 *Drosophila* species (Fig. 2C), and tested whether length had an effect on relative segment positioning. *D. santomea* was the largest at this stage, followed by *D. sechellia* and *D. virilis*. *D. persimilis* and *D. pseudoobscura* were the smallest, followed by *D. mojavensis*. Body length for the rest of the species showed few and smaller differences (Fig. 2C). We had two comparisons between sister species with large size differences, as *D. santomea* and *D. sechellia* were among the largest larvae, while their sisters, *D. yakuba* and *D. simulans*, respectively, were of roughly average size. Sister species *D. santomea* and *D. yakuba* had few differences in segment position, whereas sister species *D. sechellia* and *D. simulans* were different for every segment, suggesting that larval size is not predictive of segment position differences. Overall, larval length is not correlated with number of segment position differences (Fig. S5A, R2 = 0.0016, p-value = 0.75). In other words, having a bigger difference in body length was not correlated with having more differences in relative segment position between species. Additionally, larval length had no effect on the direction (anterior vs. posterior) of segment position differences, as we detected no relationship between body length and the number of segments that are shifted to the anterior or posterior compared to the species mean (Wilcoxon test, p-value = 1). Finally, we returned to our PCA (above), and found that PC1, which was highly correlated with the more posterior segment positions, was uncorrelated with larval length (Pearson correlation, p-value = 0.06).

### Phylogenetic analysis shows variable rates of segment position evolution

To investigate how relative segment position has evolved during *Drosophila* evolutionary history, we implemented phylogenetic methods to estimate the rates of morphological evolution over the phylogeny. We employed a multivariate Brownian motion model of evolution (Huelsenbeck and Rannala 2003; Lartillot and Poujol 2010), and tested various morphological branch rate prior models that describe how rates of morphological evolution vary across the branches of the tree. These analyses were performed in RevBayes (v. 1.0.7; Höhna et al. 2016), and are outlined in detail in File S2. Of the candidate models of segment evolution that we explored, the data significantly preferred the uncorrelated lognormal relaxed molecular clock model, which allowed rates to vary episodically along ancestor-descendant branches (*i.e.*, rates of segment evolution are not correlated between ancestor-descendant branches). Results presented here are for this model, but other morphological evolutionary models gave similar results (File S2).

Our phylogenetic analyses indicate that evolutionary rates of relative segment position are highly variable across branches of the *Drosophila* phylogeny (Figs. 4A, S8A and B). Many of the highest rates of evolution are on branches for species with sister species included in the analysis (i.e. *D. sechellia* and *D. simulans*; *D. santomea* and *D. yakuba*; *D. persimilis* and *D. pseudoobscura*). This points to segment position evolving quickly between these sibling species. The patterns vary among segments (Fig. S9). Two lineages that are also evolving rapidly, particularly in the posterior half of the larva (Figs. 4A, S9), those leading to *D. mojavensis* and *D. persimilis*, have the longest and shortest posterior-most segment (A8+tail), respectively, in our dataset (Figs. 2B, S5C). It is intriguing that the branch leading to *D. mojavensis* showed one of the fastest rates of evolution, as it is also on a long branch. This high rate sustained over a long branch is consistent with the A8+tail segment of *D. mojavensis* having the largest magnitude change in our dataset (Figs. 2B, S3C).

Given that several species had large changes in segment position toward the posterior of the larvae (Figs. 2B, 3, and S3C), we asked whether the overall rate of evolution is also higher for some segments than it is for others. We examined the rate of position evolution for each segment over all branches of the tree and found that rate of evolution varies among segments, with posterior segments exhibiting elevated rates (Fig. 4B). Segment A8 had the highest rate of change in segment position, with segment A7 also having a slightly elevated rate than the rest of the segments (Fig. 4B). The high rate of evolution for A8 is consistent with some of the largest magnitude changes in our raw data (Figs. 2B, 3) being found for this segment.

**Figure 4.**
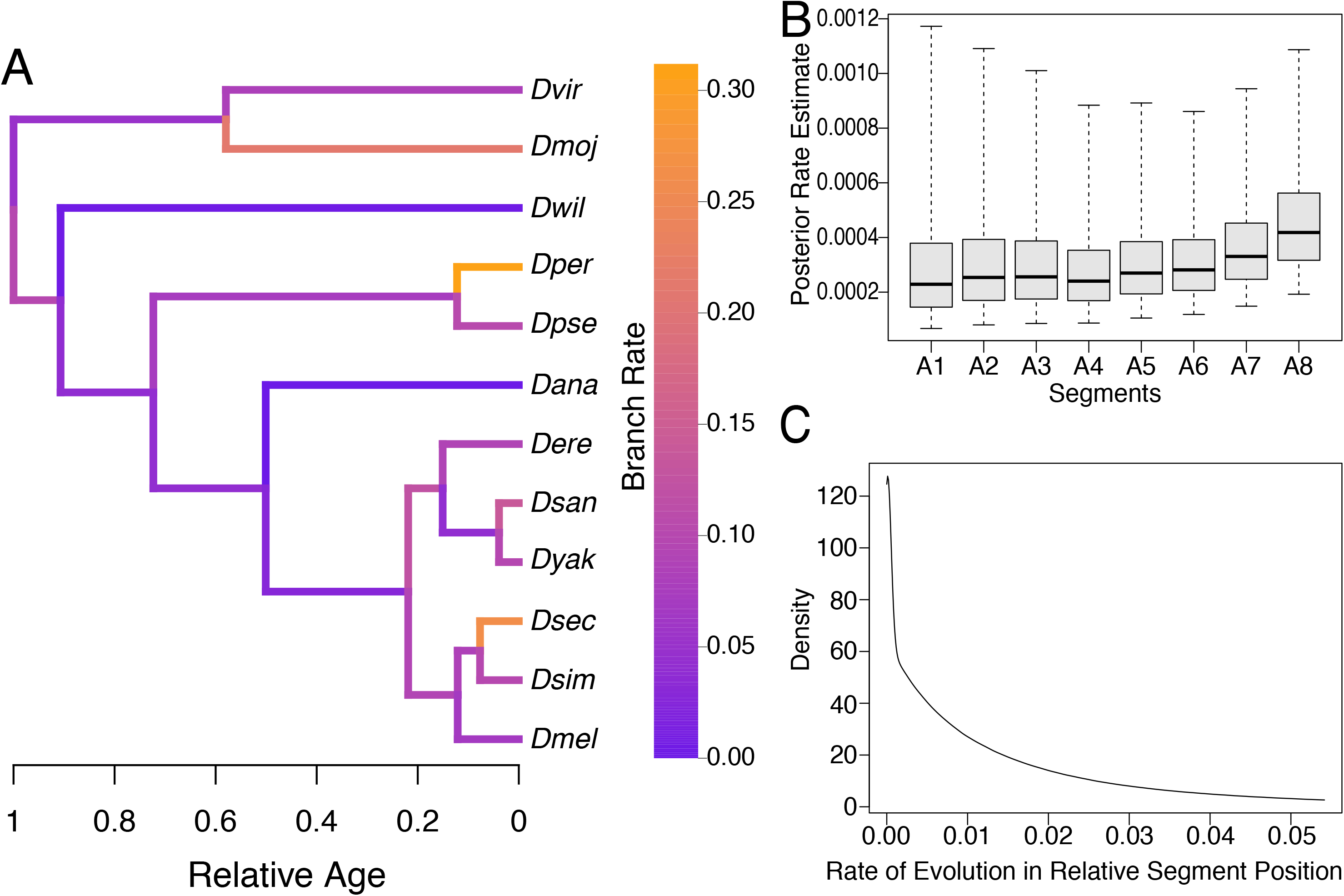
Phylogenetic analysis of segment evolution in the 12 *Drosophila* species. (A) Phylogeny inferred from nuclear loci with relative divergence times. Branches of the tree are colored to indicate the overall rate of relative segment position evolution. Rates are variable across the phylogeny, with some big differences in rate observed even between closely related species. (B) Boxplots indicate the posterior distribution of the evolutionary rate of relative segment position. The bar indicates the posterior mean rate; the boxes and whiskers indicate the 50% and 95% posterior credible intervals, respectively. The rate is higher for the most posterior segments, especially A7 and A8 (C) Marginal posterior probability distribution of the evolutionary rates of relative segment position across branches. Most changes are small in magnitude, with occasional larger changes.

To determine whether relative segment position has evolved through frequent small changes or rare large changes, we examined the distribution of the amount of change in relative segment position normalized across all branches of the *Drosophila* evolutionary tree (i.e. amount of change per unit time; see File S2, S.3.4.5 and S.3.4.6 for details). This distribution of normalized magnitude of change in segment position across all the branches of the phylogeny (Fig. 4C) showed that most segment positions on most branches have experienced changes that are small in magnitude, whereas some segment positions on some branches have experienced substantially larger changes within a given amount of evolutionary time. This analysis shows that throughout *Drosophila* evolutionary history, relative segment position has changed predominantly through small quantitative steps, with occasional large leaps.

### Is segment position along the anterior posterior axis controlled locally or globally?

Within each species, we asked how the shift in the relative position of one segment might be correlated to shifts in the relative position of other segments. This would indicate whether there is a focal point along the anterior-posterior axis controlling segment position or whether there is a more complex underlying regulatory mechanism. We found that within all 12 species, the correlation between shifts in the relative positions of any two segments is inversely proportional to the distance between the two segments (Figs. 5A, S10, and S11). We did not observe a focal point governing the shift in segments, where it might be expected that correlations would drop off as the distance from that particular point increased. It appears that small adjustments in segment position are made locally. These correlations are looking at the relationship between segment positions across individuals within a species. As the variation within species reflects developmental variation in a genetically identical line, these correlations show that developmental variation in the positioning of one segment have stronger effects on nearby segments than more distant segments. This is consistent with evidence from the embryonic stage that showed correlation between deviation from mean in expression boundary for various segmentation genes decreased with increasing distance between the expression boundary (Lott et al. 2007). Interestingly, averaged over all species, shift in the relative position of A8 was less well-correlated to shifts in the rest of the segments (Figs. 5B, S12). This suggests that shifts in the relative position of A8 are more independent than the shifts in the relative position of other segments within species (See also File S4).

Given that the relative position of A8+tail changed more independently within a species as compared to the other segments (Fig. 5B), we asked whether relative position would be more tightly regulated across the other segments in the absence of A8+tail or whether A8+tail was essential for proper segment positioning. To address this question, we calculated correlation coefficients using relative segment positions determined in the absence of A8+tail. These recalculated correlations resulted in lower average correlation coefficients between position shifts of adjacent (as well as more distant) segments as compared to when all segments were included in position calculations. (Figs. S13, S14, red line, *D. sechellia* was exceptional, see Fig. S15). This was also true when we recalculated correlations using relative segment positions determined in the absence of h+t (Figs. S13, S14, blue line). These results suggest that both ends of the larvae are needed for proper segment positioning, and removal of either one decreases the level of coordination between segments, as demonstrated by the reduction in correlation coefficients.

**Figure 5.**
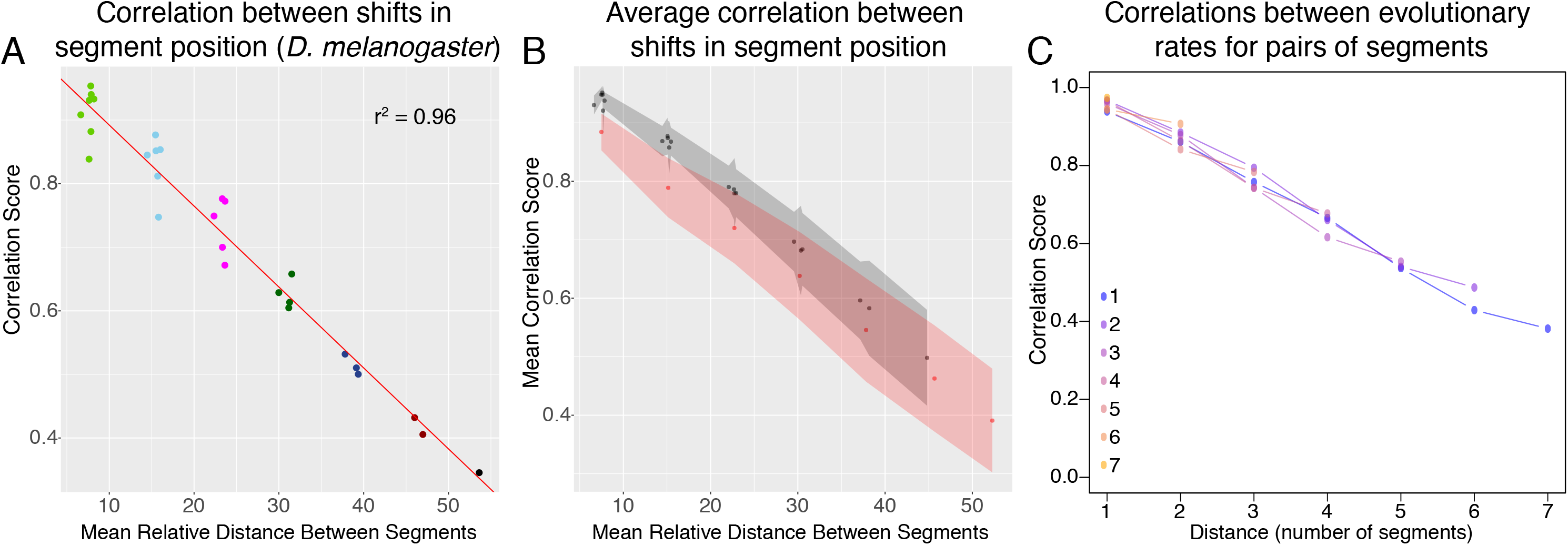
Correlations between segment positions, within and between species. (A) Within a species (here, *D. melanogaster* is shown as an example), correlations between relative segment positions are highest for neighboring segments, and fall off as the physical distance between two segments increases. Colors are used to distinguish between segment pairs with different number of segments separating them. This suggests local control of segment positioning. For plots for the rest of the species see Fig. S10. (B) This plot is similar to that in part A, but instead shows averages of correlation coefficients between segment positions over all species. Similarly, correlations between relative segment positions are highest between neighboring segments, but fall as distance between segments increase. In red are all comparisons with segment A8. At any distance, correlations with this segment are lower (in black are all other comparisons not involving A8). The gray and red shaded areas around the black and red points, respectively, indicate their 95% confidence intervals. This demonstrates that changes in the position of A8 are less correlated, and thus are more independent, than changes in the position of the rest of the segments, given its distance from any other segment. The unit of the x-axes in panels A and B, for mean relative distance between segments, is percent larval length. (C) Between species, correlations between evolutionary rates for a pair of segments are lower with increasing distance between segment pairs. This plot represents a similar analysis to part B, but in the phylogenetic framework and correlates rates of evolution for each segment. Color indicates the first segment in the comparison, with the most-blue indicating comparisons with segment 1, and the most yellow being comparisons with segment 7. Rates of evolution are highly correlated between neighboring segments, and correlations decrease with increasing distance between segments.

### Coordination of segment position evolution across the larva

Within a species, segment position seems to be controlled locally in response to developmental variation, as correlations between segments decrease with physical distance in the larva. How then are evolutionary segment position changes correlated across species? To address this question, we used the phylogenetic model to estimate correlations in the rate of position change between segments across species. These phylogenetic correlations may reflect underlying genetic correlations (in a quantitative genetic sense) or be the result of constraints placed on the system by selection pressures. Evolutionary rates of relative segment position were highly correlated across the branches of the phylogeny, and correlation decreased as the anatomical distance between segments increased (Fig. 5C). All correlations were positive, indicating that, for example, when rapid evolution occurs in one segment, changes occur across all segments. However, across species, the rate of evolution of segment A8 is not less correlated than expected, given its distance from other segments. This suggests that while this segment may evolve at a faster rate (Fig. 4B), on average it does not seem to be evolving in a less coordinated fashion than the other segments, when the whole phylogeny is considered.

## Discussion

Many critical developmental processes and traits are known to be robust, in that they are produced faithfully despite variation encountered in ontogeny. This tolerance of variation does have limits, however, and exceeding these limits may disrupt the development of these traits in such a way that development does not proceed. Hence, robustness is desirable during development because it assures the precise production of critical traits and processes within a range of developmental conditions.

Given the suppression of phenotypic variation in robust traits, their evolution has been a subject of considerable interest to researchers over the years, and has produced both theoretical and empirical work (Wagner 2005; Félix and Wagner 2006; Masel and Siegal 2009; Siegal and Leu 2014; Payne and Wagner 2019). One proposal for the way for a robust trait to evolve is for conditions to exceed the tolerance for variation in that trait, and expose genetic variation that had previously been masked, i.e. cryptic genetic variation, by the very robustness of the trait (Rutherford and Lindquist 1998; Gibson and Dworkin 2004; Paaby and Rockman 2014). Exposing any amount of genetic variation of unknown consequence to a critical trait at a time of great stress seems like a dangerous proposition in a multicellular animal. Our data supports an alternative model, where there is always some small amount of variation available, even in the most robust traits, and that quantitative changes may evolve in robust traits without disrupting robustness.

Here, we focus on a well-studied robust process and trait, segmentation in *Drosophila*. Previous studies have highlighted the ability of the segmentation network to produce precisely localized segment markers in embryogenesis despite experimentally produced perturbations in development (Félix and Wagner 2006; Masel and Siegal 2009). Our study extends this to measure the trait of segmentation in larvae, across 12 *Drosophila* species. Here, we demonstrate that segmentation varies considerably across species, and that the rate of evolution of segment position varies considerably across the phylogeny as well. The variation in segment position consists mostly of small magnitude differences, suggesting that this trait evolves by many small changes, with occasional larger changes. While it can be difficult to quantitatively compare earlier stages of segmentation in the embryo to segments in the larva, this result is consistent with patterning differences in the embryo observed between species (Lott et al. 2007; 2010). Our data shows that segment position differences between species exist also in later developmental stages. Moreover, each line of each species measured here precisely produces its characteristic segmentation pattern in the larva. This points to the ability of segmentation to evolve without losing its ability to be precisely localized (Lott et al. 2010). While we did not test the robustness of each species to genetic and environmental variation specifically, this also suggests that segmentation can evolve and remain robust to stochastic variation. If segmentation can evolve small quantitative changes over evolutionary time without losing robustness, then this suggests that not all genetic variation in this trait is cryptic (Miles et al. 2010; Jiang et al. 2015), and small-scale differences (and the occasional larger-scale difference) in segmentation patterns may be available to selection (Weber 1992). Or, perhaps there is a neutral space where a range of segment positions are tolerated, and stabilizing selection keeps them in that range.

The evolutionary patterns observed here may also be simply what might be expected of complex traits generally, with changes in a number of genes producing small quantitative changes in phenotype over evolutionary time. Here, we find that segment position seems locally determined along the length of an individual, as position shifts are highly correlated between adjacent segments, and correlations drop off over distance, consistent with a previous result within species in embryos (Lott et al. 2007). This is also consistent with the known regulatory interactions in this network, where patterns are refined over developmental time (DiNardo and O’Farrell 1987; Surkova et al. 2008) and regulation of segment patterning becomes more localized to a smaller portion of the developing animal at each stage as compared to the previous one (Nüsslein-Volhard and Wieschaus 1980; Kornberg and Tabata 1993; Nüsslein-Volhard et al. 2008). Our results suggests that changes in many genes produced the observed differences observed between species, and hence, it may be difficult to determine the identity of individual genetic changes that may cause the differences in segment position. However, the genetic network behind segment patterning has been fertile ground for modeling in the embryo (Jaeger et al. 2004; Manu et al. 2009a; Wunderlich and Depace 2011; Bentovim et al. 2017; Clark 2017; Verd et al. 2018; Petkova et al. 2019). Implementing a modeling framework developed for the embryonic stage, and extending it past the embryonic segment polarity stage to the positions of segments in the larva, is likely the best approach for identifying potential genetic causes of the differences observed here.

Across the larva, the most striking pattern we observed was in the posterior-most segment, from A8 to the tail of the animal. We found that this segment has the largest magnitude of differences between species, and evolves faster between species than the other segments. It is the most independently controlled segment within a species, with correlations between its position and all other segments being the lowest found. As the h+t region anterior of A1 does not share these properties, it is not simply a feature of terminal segments, but specific to the A8+tail region. The A8+tail region contains the posterior spiracles, important breathing structures in the larva (Hu and Castelli-Gair 1999), as well as the genital imaginal disc from which the genital structures in the adult are produced (Sánchez and Guerrero 2001). While genital structures are known to evolve rapidly between species (Eberhard 2013), it is unclear whether this would produce differences in the entire segment in which the genital imaginal disc is found. Alternatively, the differing conditions in which larvae find themselves may produce differences in behavior that would require differences in posterior spiracle length. We are currently exploring this possibility in relatives of *D. mojavensis*, as this species has an exceptionally long A8+tail segment. In our care, *D. mojavensis* larvae burrow deeply in food with only their long posterior spiracles visible above the surface of the food. As *D. mojavensis* and its relatives are desert-dwelling cactus specialists (Oliveira et al. 2012), perhaps the larvae burrow deeply into cactus in their natural environment to find a more hospitable microclimate (McKenzie and McKechnie 1979; Green et al. 1983).

Overall, our results show for the first time that segment position in *Drosophila* has changed frequently throughout the evolutionary history of the genus. The changes were mostly small in magnitude, presumably representing only small perturbations to the development of the organism. Rate of evolution of this trait varies across the phylogeny, with larger magnitude differences and a higher rate of evolution observed for the tail of the larva. Changes in segment position within species, as well as rates of evolution between species, are highly correlated between neighboring segments, indicating the highly coordinated nature of the genetic network underlying this trait. Future studies are needed to unravel the nature of the genetic changes underlying segment position differences between species and whether any of the observed phenotypic differences between species might be adaptive.

## Acknowledgements

We thank Lott Lab members Emily L. Cartwright, Anna A. Feitzinger, Charles S. Omura for comments on the manuscript; and Graham Coop for comments and statistical help. We also thank The National Drosophila Species Stock Center for the Drosophila species used in this project. This work was financially supported in part by National Institutes of Health grants, R01GM111362, R00GM098448, and start-up funds provided to S.E.L. by the University of California, Davis.

---

**Figure S1.**
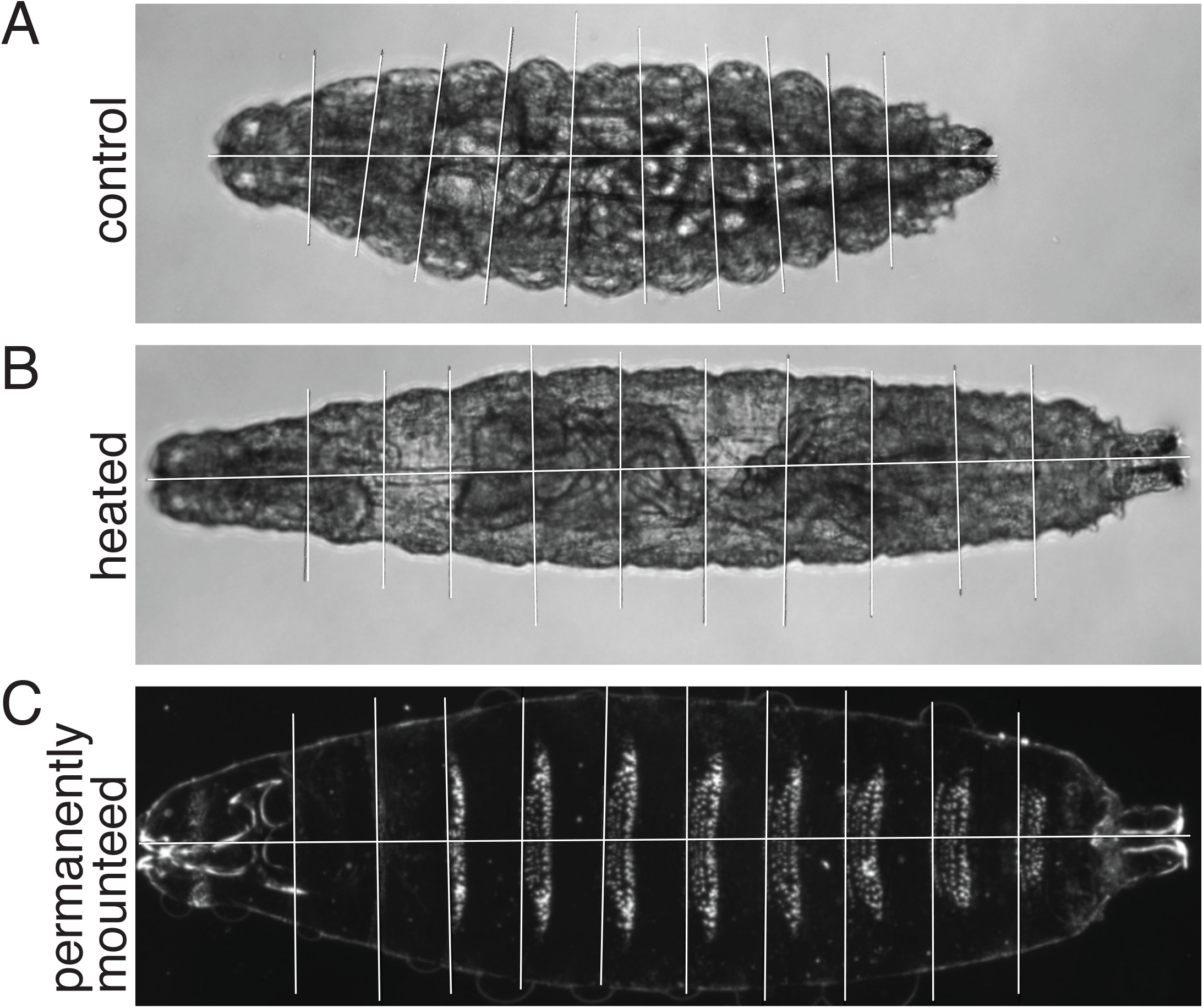
Testing the effect of larval mounting procedures on relative segment position. For this experiment, relative segment position measurements were taken before and after multiple stages of the mounting protocol, to test the effect of the mounting protocol on segment position across multiple species (see Methods) (A) Bright field image of a *D. melanogaster* first instar larva, hatched in water, iced for 2-3 minutes before imaging. (B) The same *D. melanogaster* larva heated at 60°C for an hour prior to bright field imaging. (C) Dark field image of the same *D. melanogaster* larva, mounted using PVA, and incubated in the 60°C oven overnight. This represents the end stage of our mounting protocol. White horizontal lines mark the full length of the larvae, vertical lines mark the segment borders.

**Figure S2.**
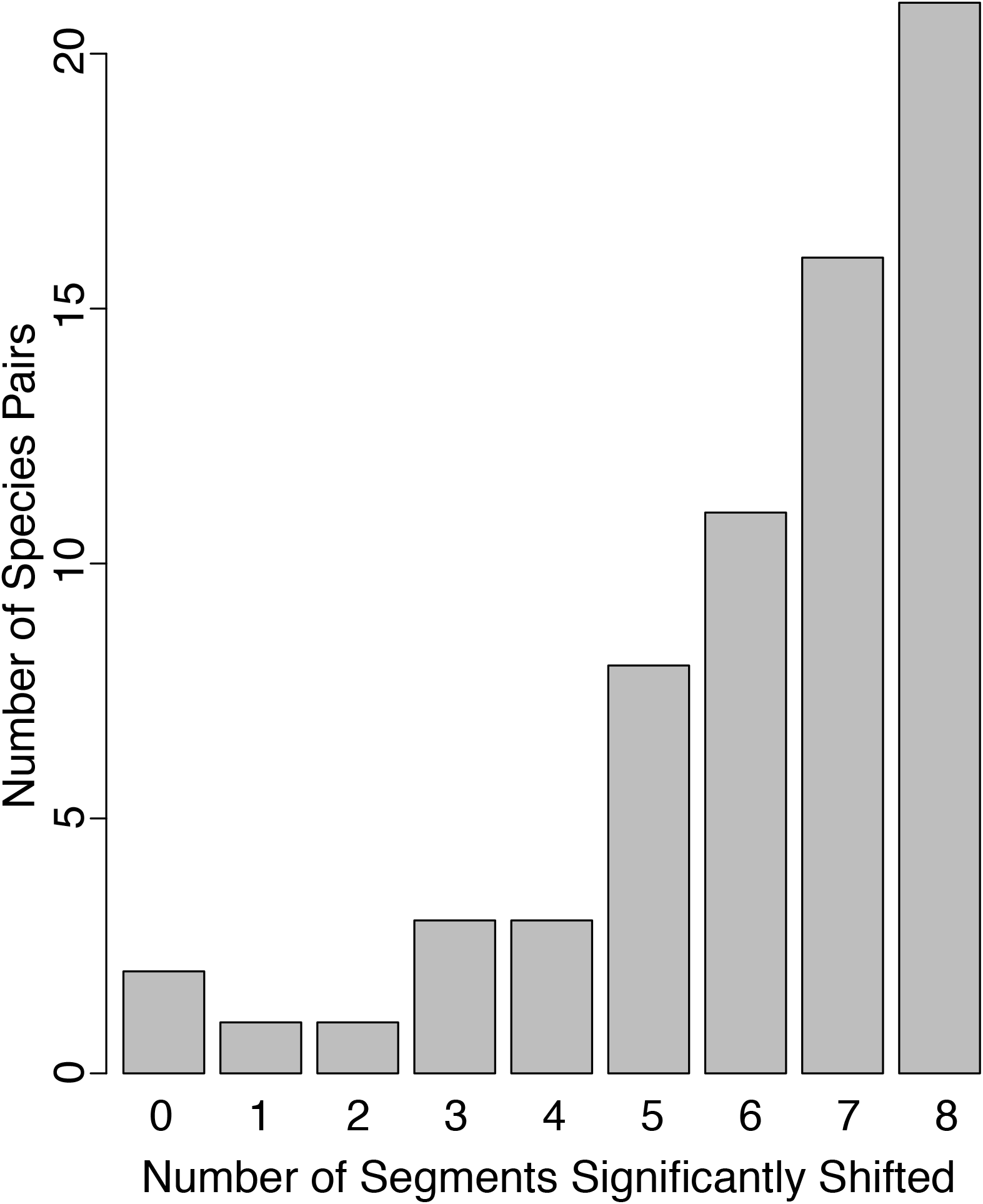
Comparing pairs of species, most segments were in different relative positions. For the majority of species pair comparisons, five or more segments had significantly shifted relative positions (t-tests with Bonferroni correction, p-value ≤ 0.05). The graph shows the number of species pairs on the y-axis and number of segments significantly shifted (see Methods) on the x-axis.

**Figure S3.**
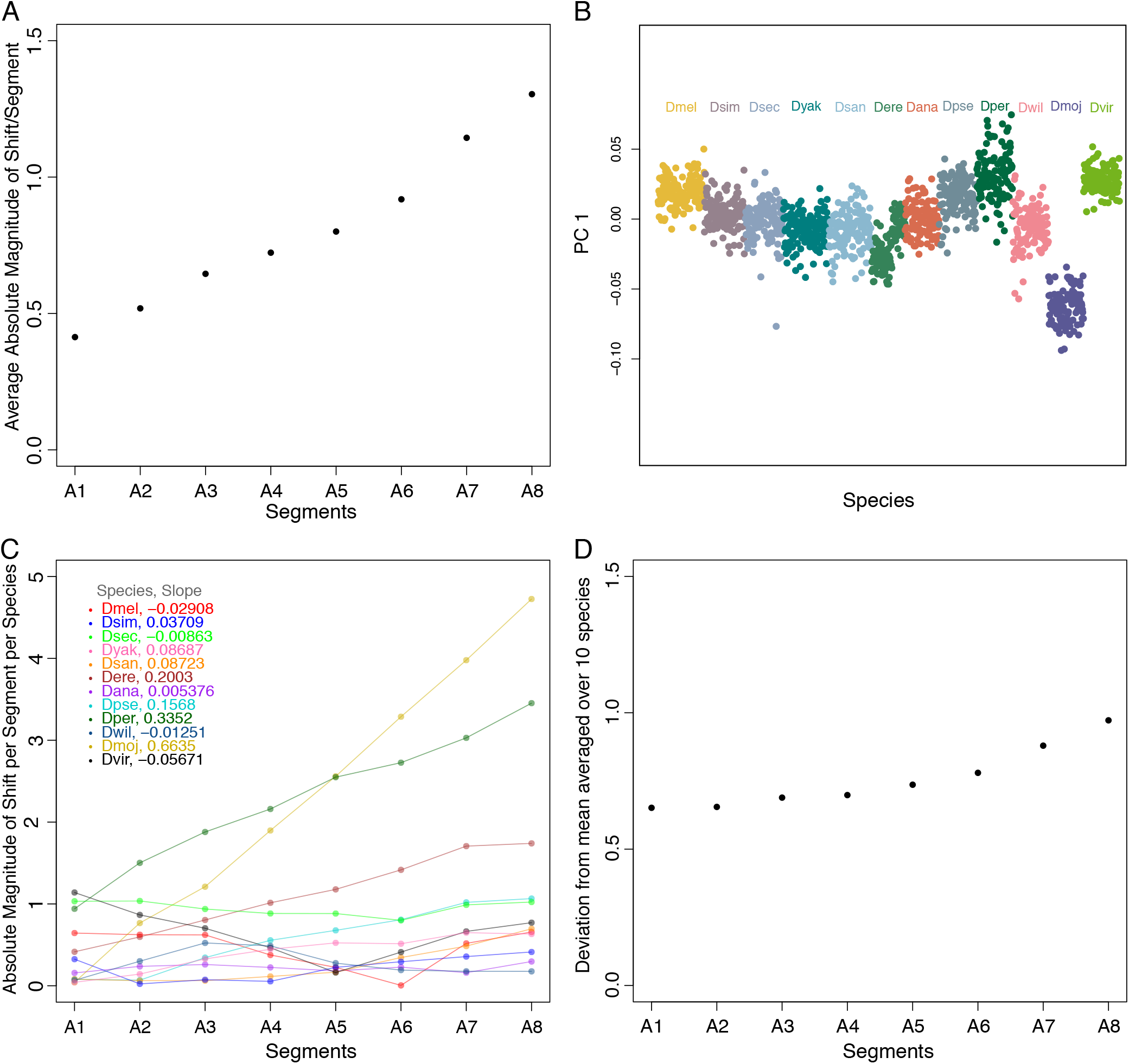
The posterior-most segments show the largest magnitude in differences from mean across species. (A) Average absolute magnitude of shift in the relative position of each segment, as compared to the across species mean, increases from anterior to the posterior of the larvae. On the y-axis of this graph is absolute magnitude of shift, in percent larval length, for each segment averaged over all species, on the x-axis are abdominal segments A1 through A8. (B) From principal component analysis, PC1 is plotted for all species. Positive values correspond to posterior shifts of segments, negative values indicate anterior shifts of segments, relative to mean centered positions for each segment. PC1 explains 88% of the variance and is highly positively correlated with the positions of the most posterior segments (Pearson correlation coefficients with each segment, A1-A8: A1= −0.15, A2=0.11, A3=0.28, A4=0.49, A5=0.64, A6=0.77, A7=0.85, A8=0.88). Note the similarity in pattern to Figure 3, with *D. mojavensis* having the largest deviation from the rest of the species, with a dramatic shift toward the anterior (negative PC1 values) and *D. persimilis* shifted toward the posterior (positive PC1 values). (C) Absolute magnitude of shift changes from anterior to posterior differently for each of the 12 *Drosophila* species. The graph shows absolute magnitude of shift in percent larval length on the y-axis and abdominal segments A1 through A8 on the x-axis. The legend in the graph shows the slopes calculated for each trend form anterior to posterior. Lines in the graph and the legend are color coordinated for each species. (D) Average absolute magnitude of shift in the relative position of each segment, as compared to the across species mean, increases from anterior to the posterior of the larvae even in the absence of segment position data for *D. persimilis* and *D. mojavensis* (compare to Figure S3A, this representation is the same, except with *D. persimilis* and *D. mojavensis* removed from the calculation of the mean). On the y-axis of this graph is absolute magnitude of shift, in percent larval length, for each segment averaged over all species, on the x-axis are abdominal segments A1 through A8.

**Figure S4.**
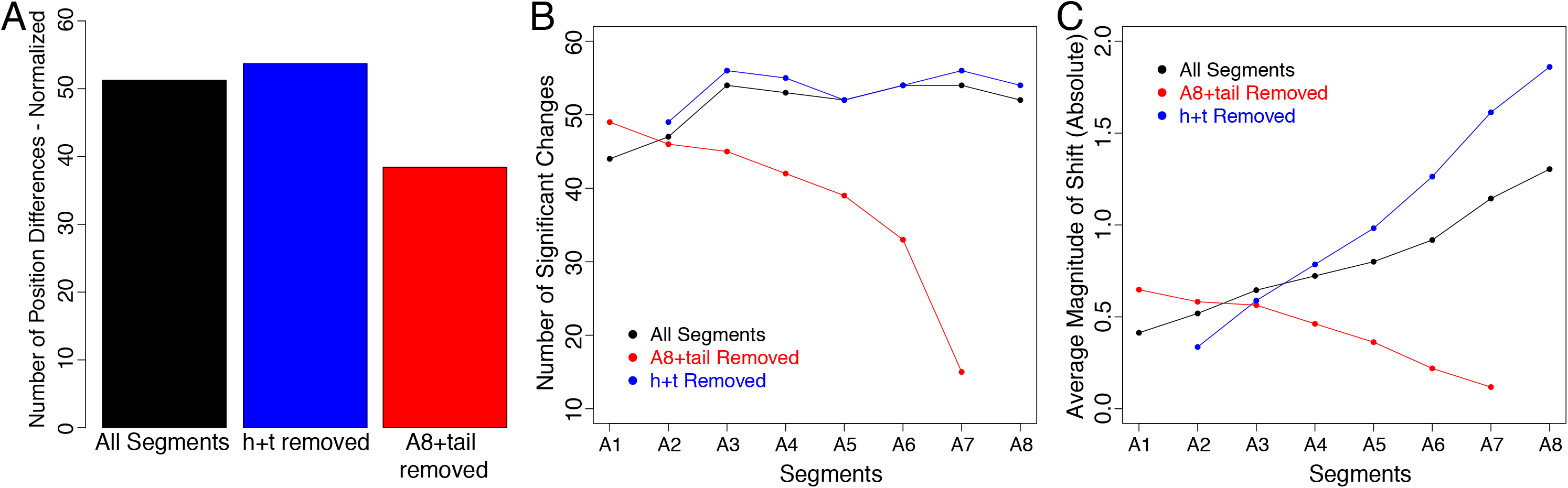
The A8+tail region is responsible for much of the number and magnitude of significant differences between species. For all these plots, compare the effect of when A8+tail is removed to when another terminal segment (h+t) is removed, to determine whether the effect is A8+tail specific or simply an effect of removing a terminal segment. (A) Comparison of numbers of significant position changes when segment positions were calculated with all segments included, with h+t removed or with A8+tail removed. Removal of A8+tail shows a decrease, whereas removal of h+t shows an increase, in the number of segments that have significantly changed their relative position as compared to when all eight abdominal segments were included in the position calculations. Y-axis is total number of significant changes, normalized to the number of segments included in position calculations. (B) This plot shows the same data as part A, but illustrates how the differences in number change across the length of the larva. When A8+tail was removed from position calculations, number of significant changes in relative segment position went down sharply towards the posterior end of the larvae. However, when h+t was removed in the same manner, number of significant changes in relative segment position followed the same pattern from anterior to posterior as when all abdominal segments were included in position calculations. y-axis is number of significant changes in relative position normalized to the number of segments included in position calculations. (C) Average absolute magnitude of shift (from the species mean) for each segment went down sharply towards the posterior of the larvae when A8+tail was removed from segment position calculations. On the other hand, removing h+t followed the same pattern as when all segments were included in calculations, but showed a bigger deviation from the species mean towards the posterior of the larvae. y-axis is the average magnitude of shift (from the species mean) in segment position, in percent larval length, ignoring the direction. X-axis is abdominal segments in both panels B and C.

**Figure S5.**
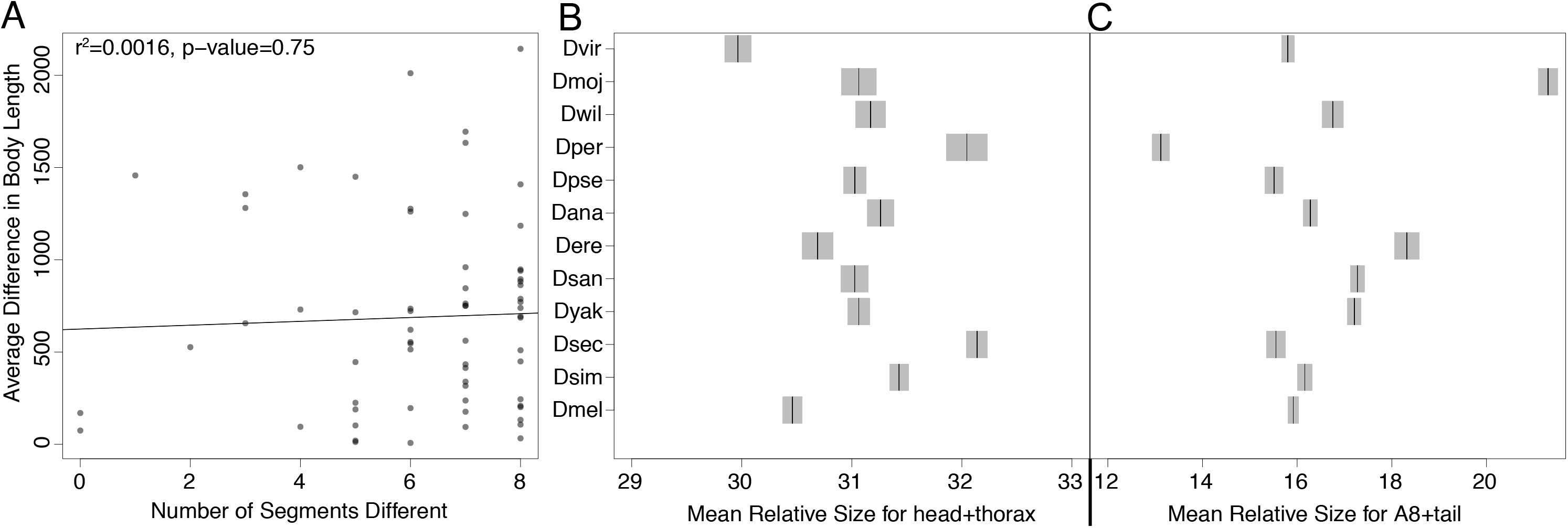
Features of body length. (A) Average body length difference between a given pair of species is not correlated with the number of segments with significant position shifts between the species. The y-axis in this graph shows average difference in body length (in pixels) between each pair of species and the x-axis shows the number of segments with significant position shifts in the same species pair. (B) This graph shows differences in mean relative size for h+t, in percent larval length, among 12 *Drosophila* species. The black bars indicate the mean and the gray shaded area is 95% confidence interval for each mean. (C) This graph shows differences in mean relative size for A8+tail, in percent larval length, among 12 *Drosophila* species. The black bars indicate the mean and the gray shaded area is 95% confidence interval for each mean. Parts B and C represent transformations of the position and length measurements, to highlight the length of the terminal portions of the larva.

**Figure S6.**
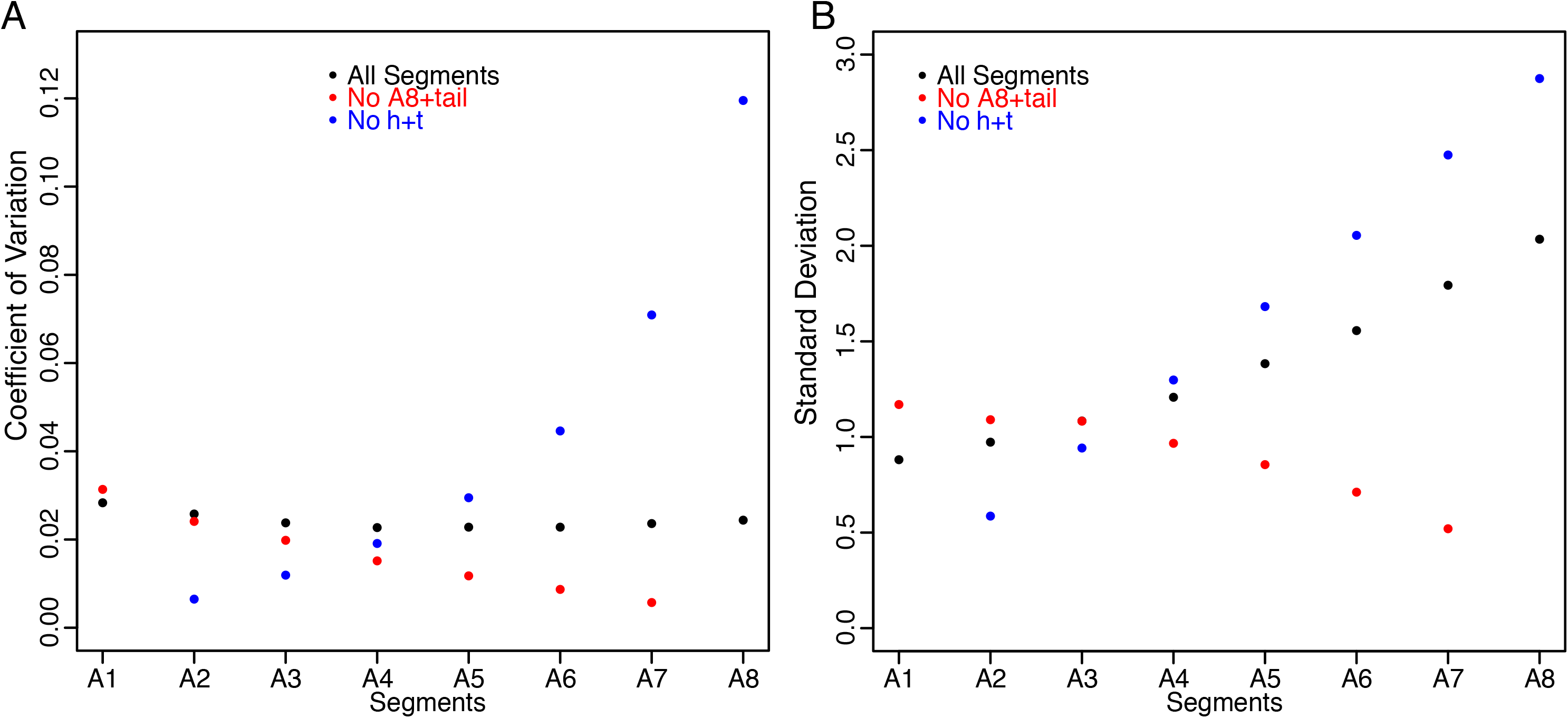
Coefficient of variation in segment positioning stays low throughout the larva, indicating that the relative positions of all segments are equally as precisely determined. However, standard deviation of segment positioning increases towards the posterior end of the larva, as the measurement values increase (relative position is measured in % of larval length from the anterior end). (A) Change in coefficient of variation, averaged over 12 *Drosophila* species, for all segments from anterior to posterior of the larva. The y-axis shows coefficient of variation, and the x-axis depicts each abdominal segment. When segment positions were calculated with all segments included (black dots), coefficient of variation (noise) did not change along the anterior-posterior axis of the larva, and remained low. Removal of A8+tail (red dots) overall decreased noise levels in segment positioning, whereas removal of h+t (blue dots) increased noise levels towards the posterior of the larva. (B) Change in standard deviation, averaged over 12 species, for all segments from anterior to posterior of the larva. The y-axis shows standard deviation and the x-axis depicts each abdominal segment. When segment positions were calculated with all segments included (black dots), standard deviation of segment positions increased towards the posterior of the larva. Removal of A8+tail (red dots) decreased standard deviations in the posterior, whereas removal of h+t (blue dots) increased standard deviations towards the posterior of the larva.

**Figure S7.**
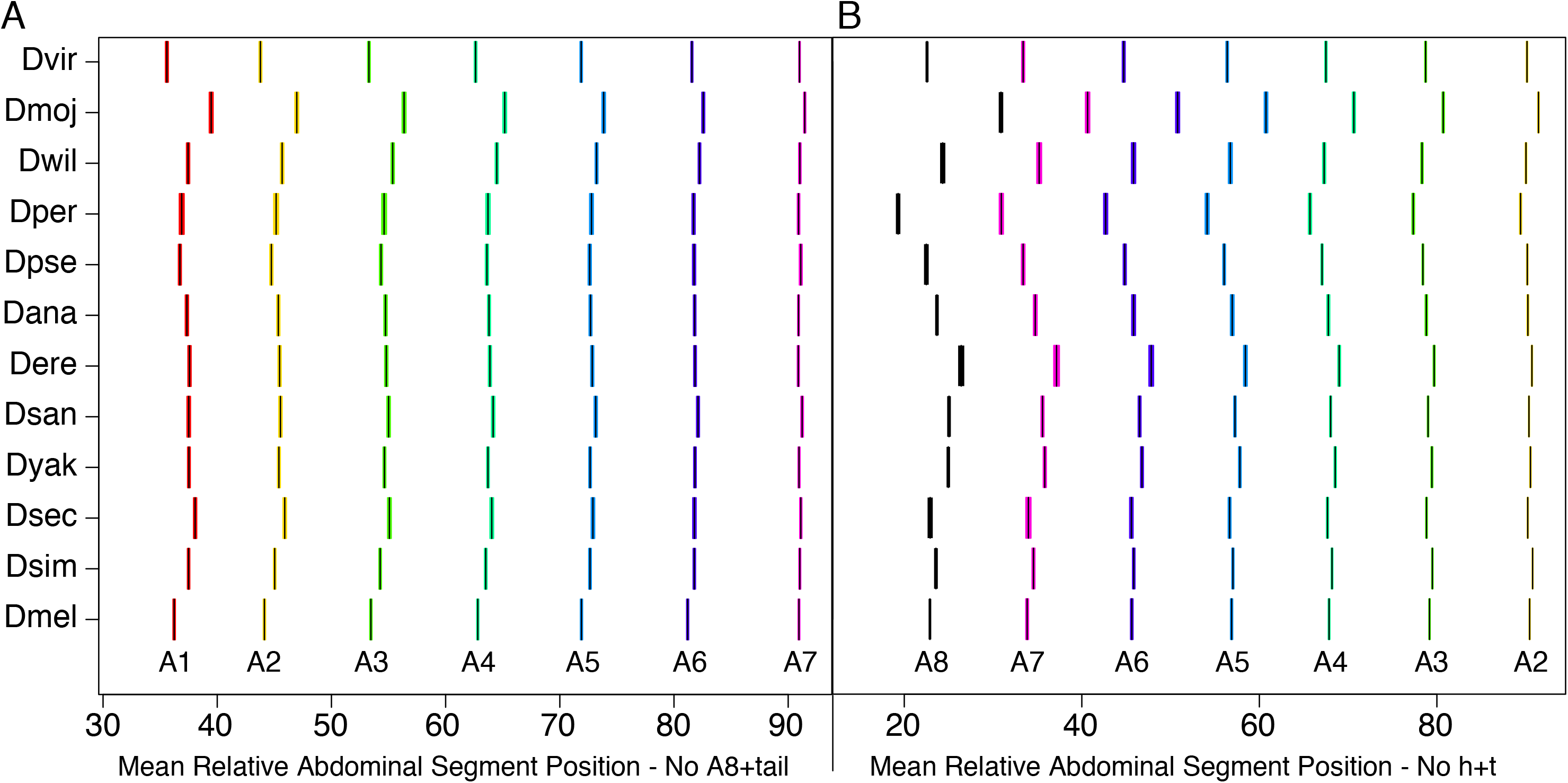
Relative segment position and body size is highly variable among 12 species of *Drosophila*, though less so when A8+ tail is removed. These graphs have the same format as Figure 2B and shows differences in relative segment position between species. (A) Relative segment positions are calculated in the absence of A8+tail. Notice a substantial number of nonsignificant differences as compared to Figure 2B. (B) Relative segment positions are calculated in the absence of h+t and also with measurements taken from the posterior of the larva instead of the anterior. As compared to Figure 2B the segment order is flipped, but the pattern of change in segment position is very similar. In both graphs, each black bar depicts the mean for each segment in each species. The colored areas are 95% confidence intervals. The y-axis depicts 12 *Drosophila* species and x-axis shows mean relative abdominal segment position in percent larval length.

**Figure S8.**
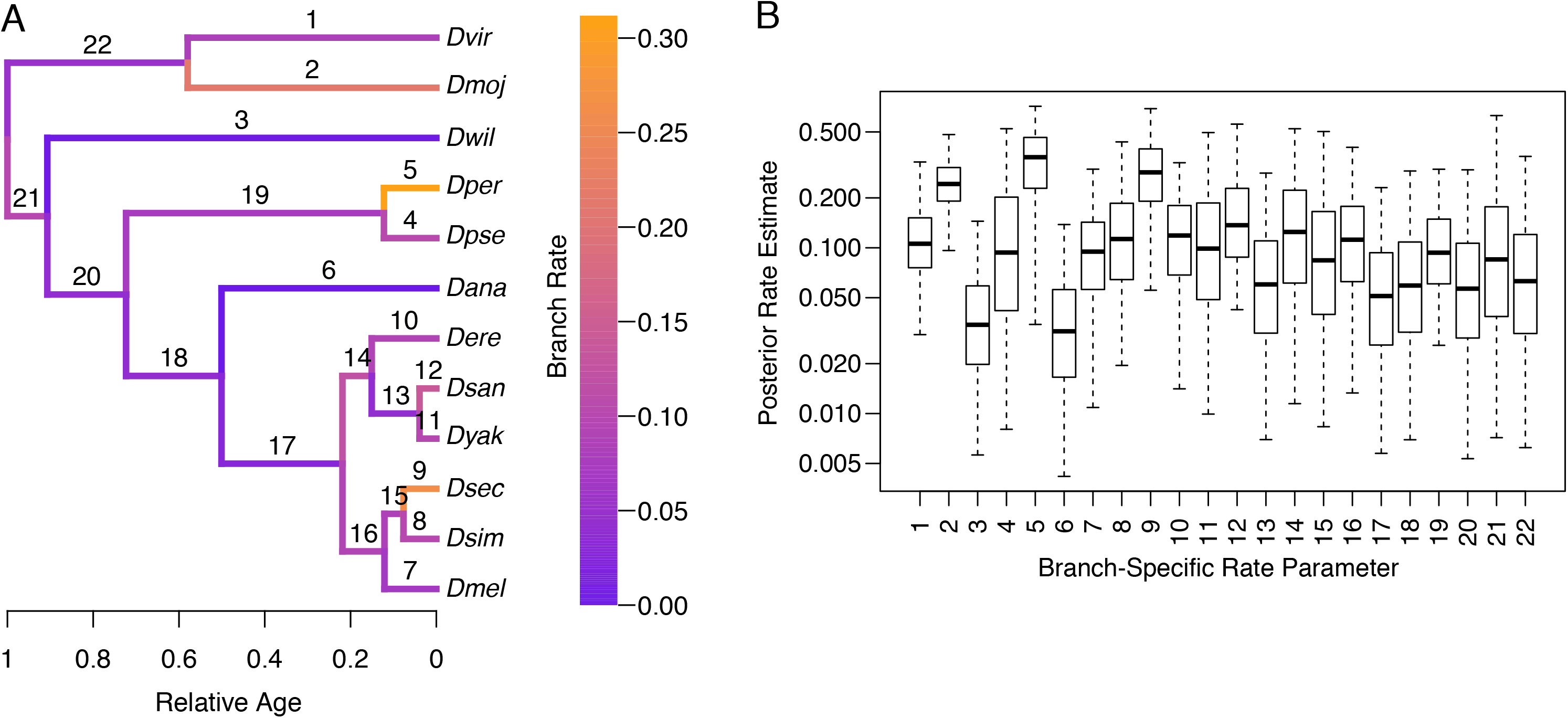
Phylogenetic analysis of relative segment evolution in the 12 *Drosophila* species, with posterior rate estimates for all branches. A) Like Figure 4A, the phylogeny was inferred from nuclear loci with relative divergence times, and branches of the tree are colored to indicate the overall rate of relative segment position evolution. Here, the branches are numbered to correspond with the values pictured in part B. B) Box plot depicting rate of evolution estimate for each branch of the phylogeny, x-axis corresponds to the branch numbers indicated on the tree in part A.

**Figure S9.**
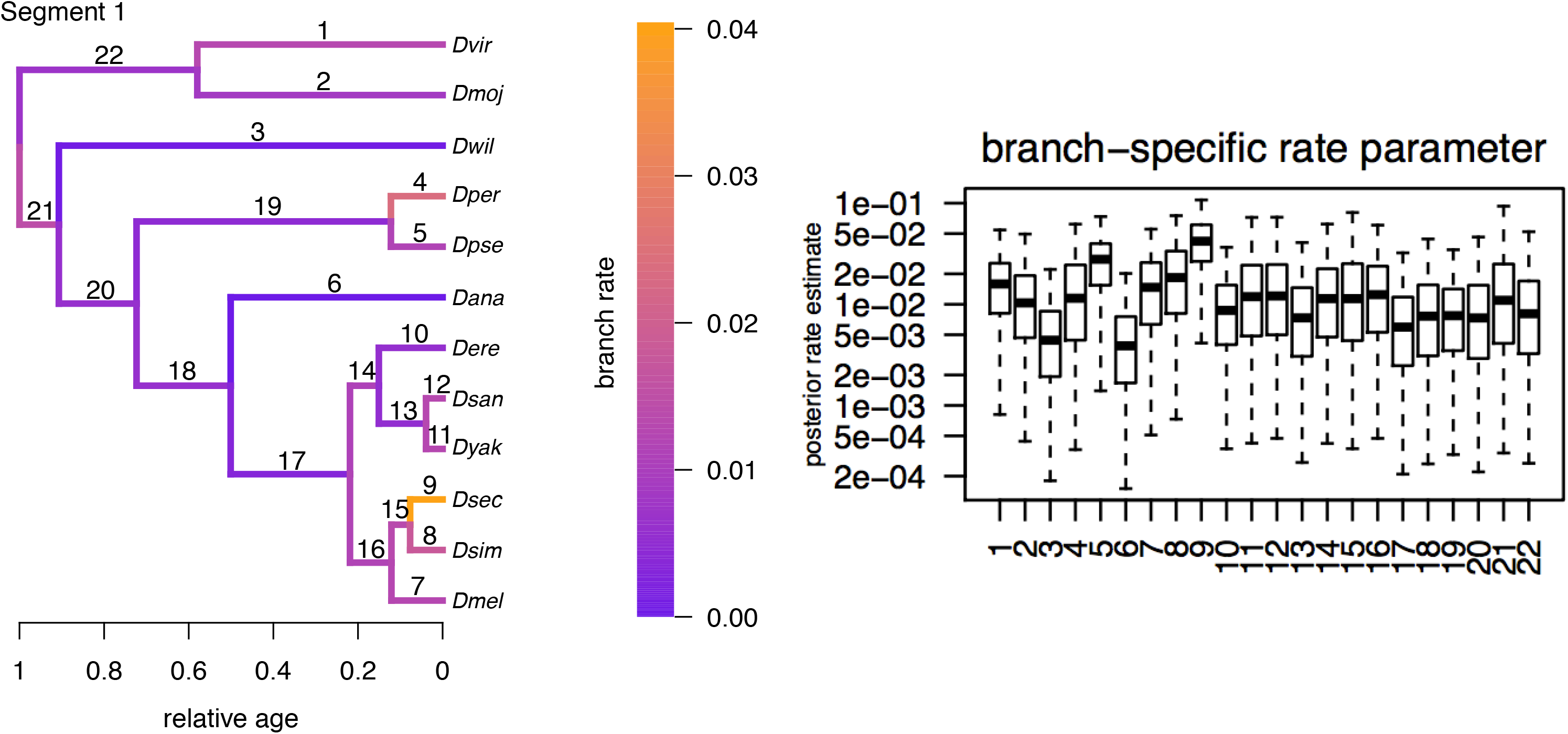
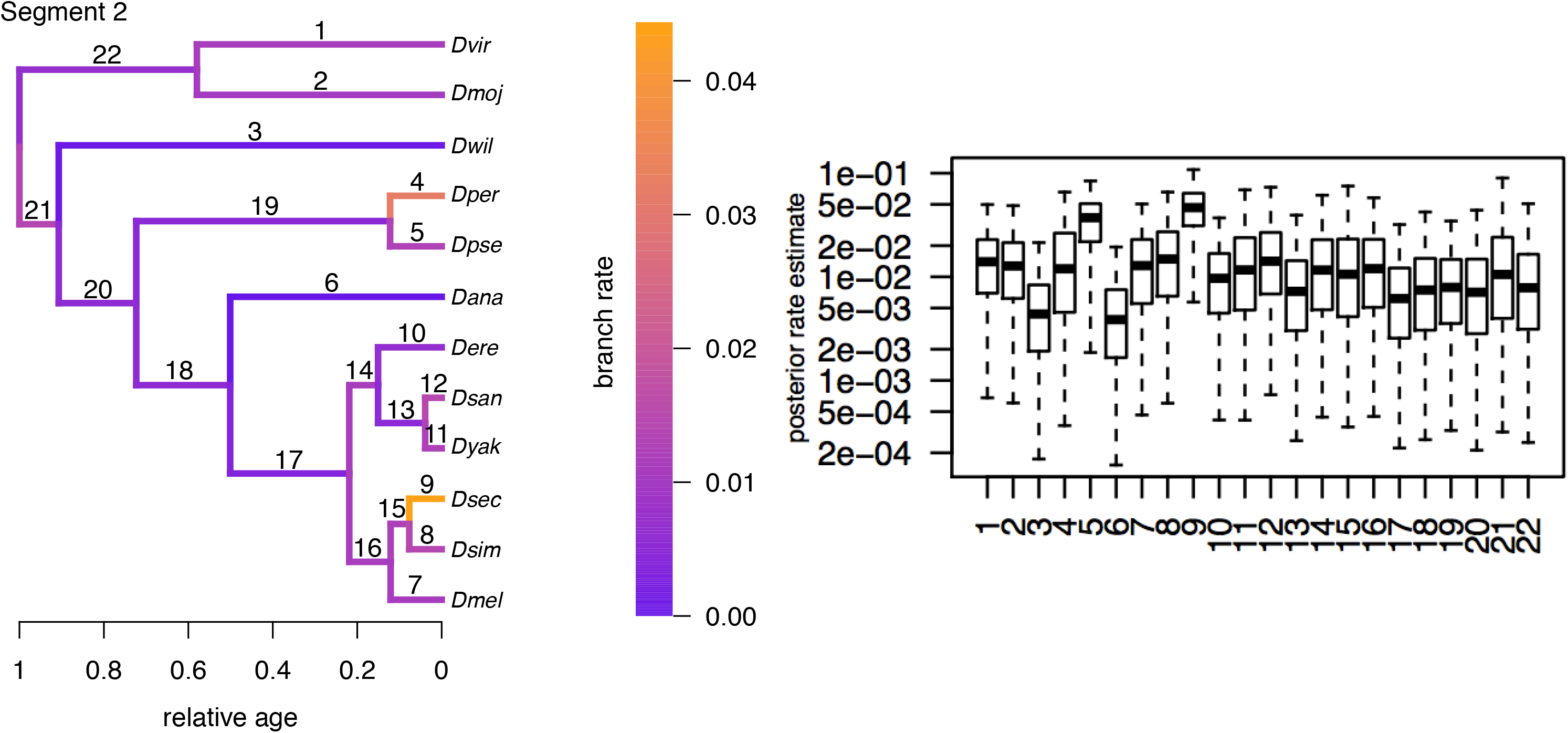
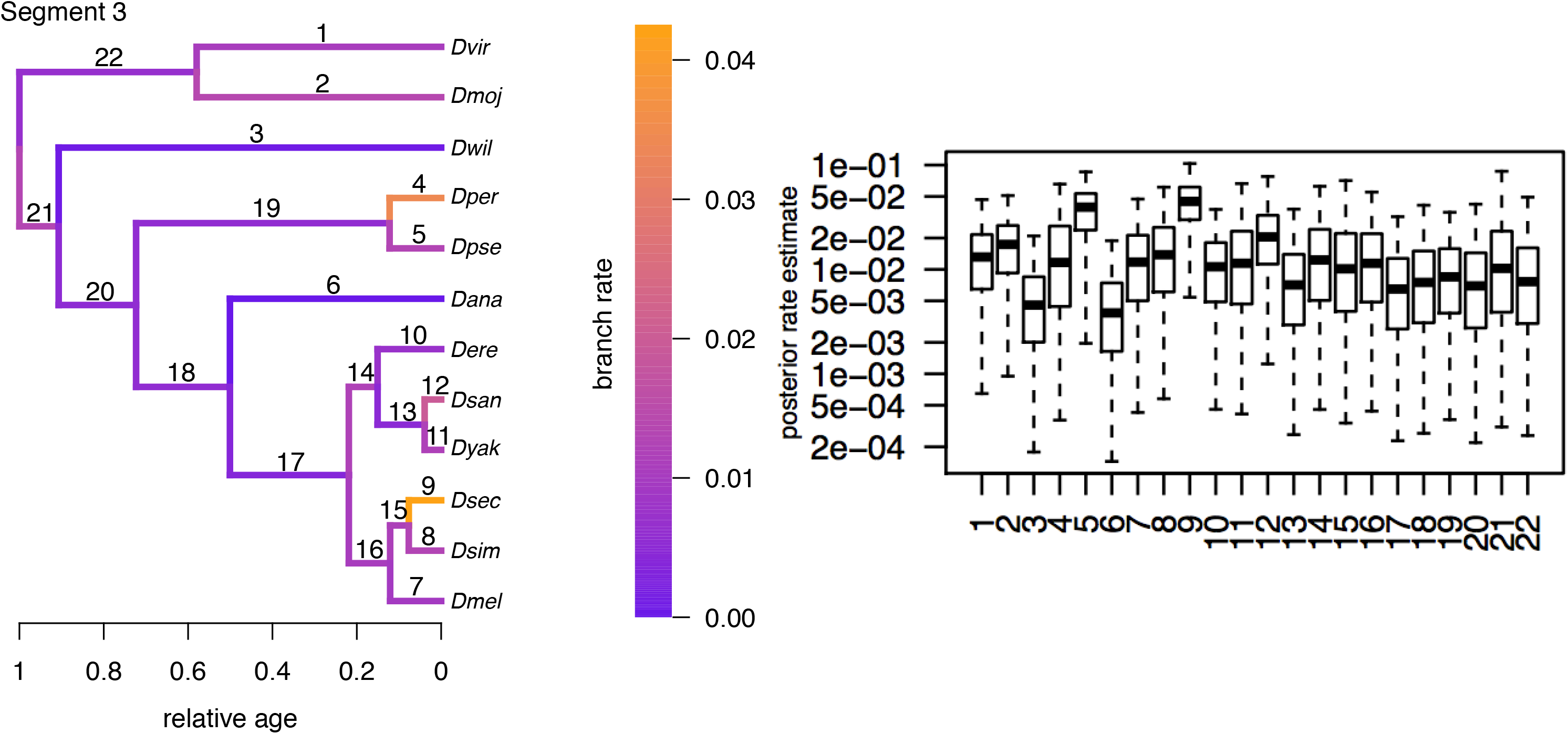
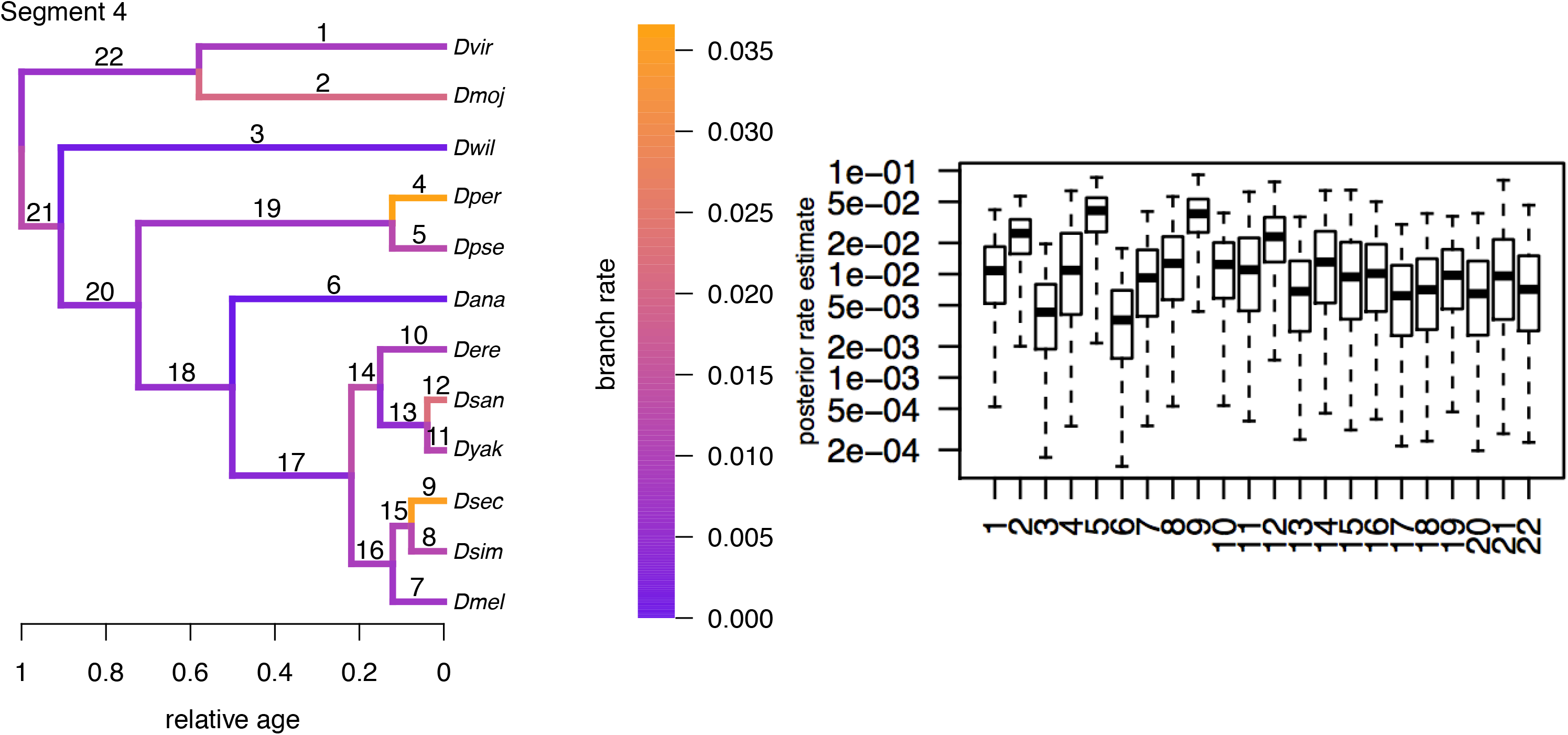
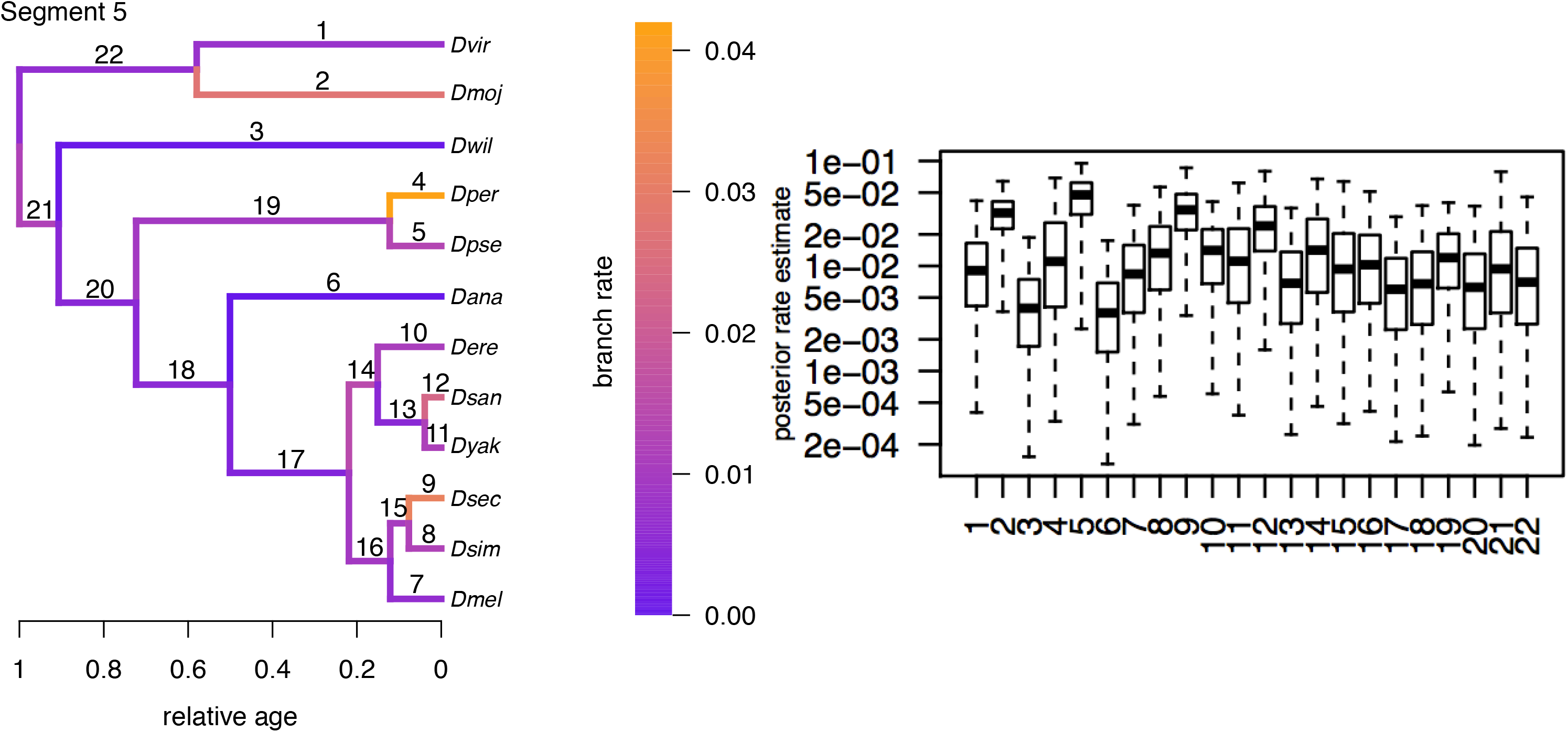
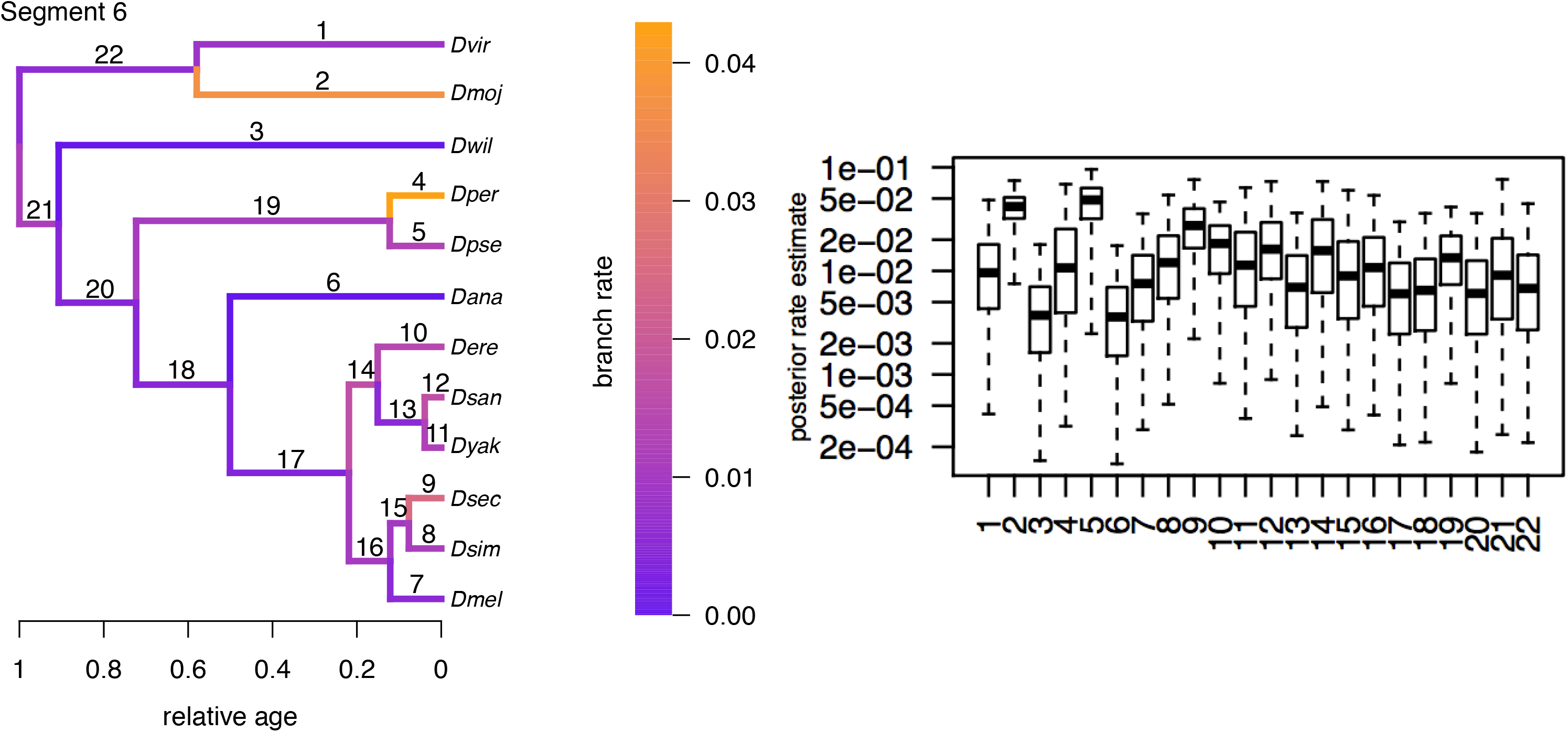
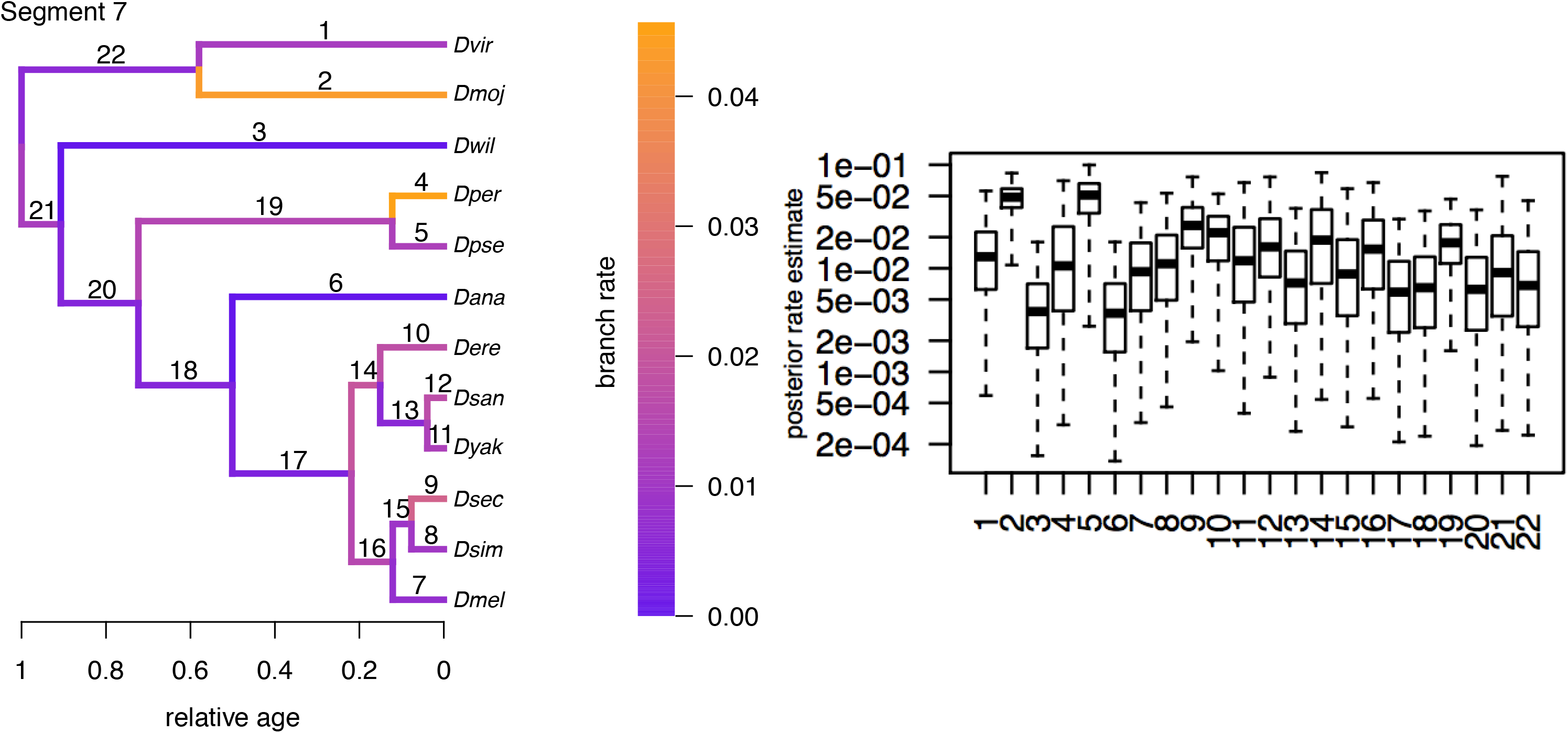
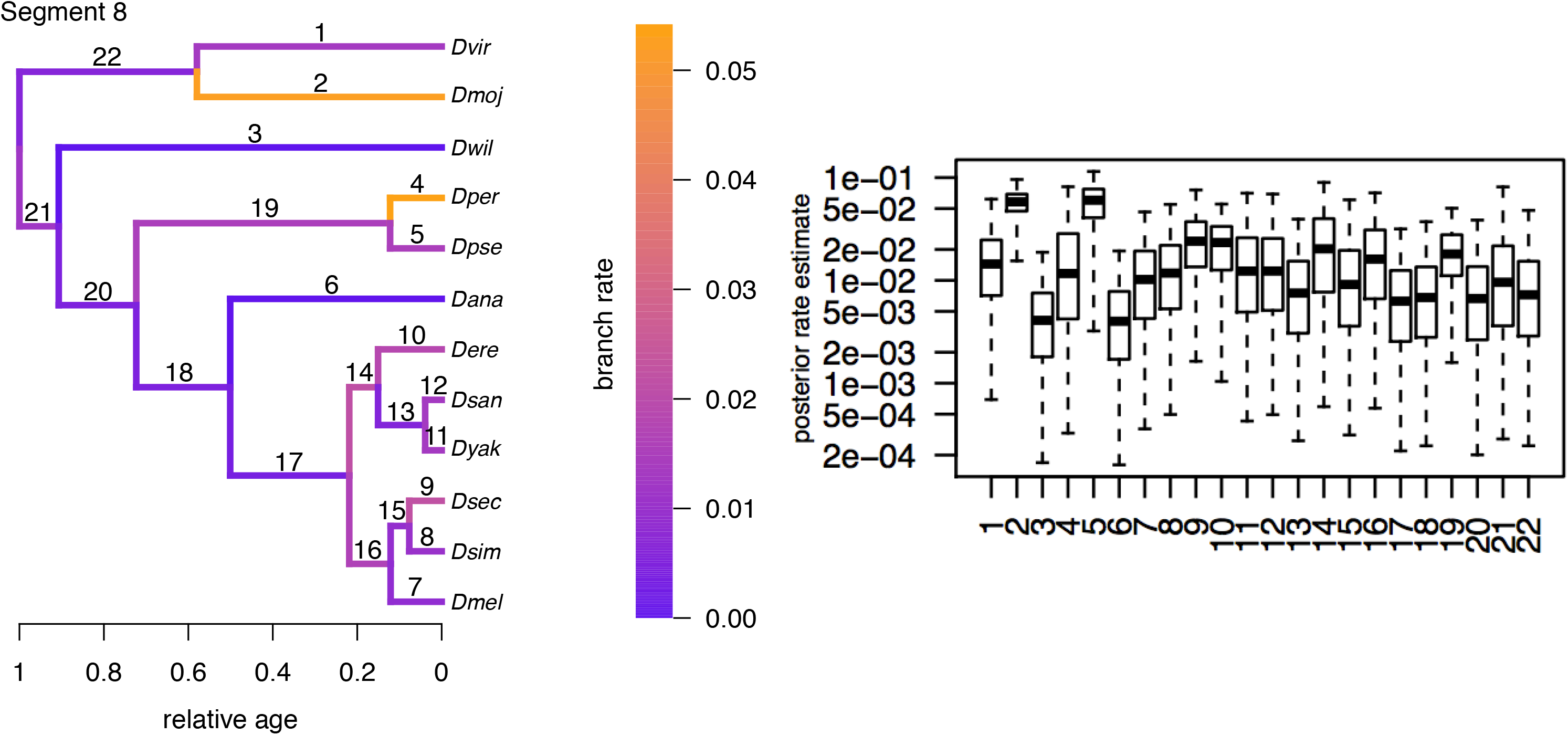
Phylogenetic analysis of relative segment position for each segment. Like Figure 4A, this figure shows a phylogeny inferred from nuclear loci with relative divergence times, and the branches of the tree are colored to indicate the overall rate of segment position evolution. Like Figure S8, the tree diagrams also include numbers for each linage, and are accompanied by a box plot showing the estimates for rates of segment evolution by each numbered lineage. Here, there are 8 plots, to show rate estimates for each abdominal segment separately.

**Figure S10.** Correlations between relative segment positions within each species. This series of graphs are constructed the same as Figure 5A, which was for *D. melanogaste*r, for each of the 12 *Drosophila* species. The y-axis shows correlation coefficients and the x-axis shows mean relative distance between pairs of segments in percent larval length.

**Figure S11.** Correlation coefficient heat maps for each of the 12 *Drosophila* species. This shows the correlation coefficient between shifts in the relative position (as compared to the across species mean) of each pair of segments within a species. Red indicates high positive correlation and white indicates low positive correlation. Pearson correlation coefficients are printed on each square box. Notice that for most species, the color scheme is lighter at the right edge of the triangle as compared to the bottom edge of the triangle. This is consistent with other figures showing shifts in the position of A8 has a lower correlation with shifts in the position of other segments, regardless of the distance in between any pair of segments.

**Figure S12.** Correlations between segment positions, comparing correlations including segment A8 with all other pairwise comparisons the same number of segments apart. This series of graphs represent the information from Figure 5B, but broken out by how many segments separate the two segments being compared. Each graph has mean correlation coefficient over all species on the y-axis, and mean relative distance in percent larval length over all species on the x-axis. Segments that are paired with A8 are in red. Error bars are 95% confidence intervals.

**Figure S13.** When all segments are included in position calculations, mean correlation coefficients between neighboring segments are highest, ever so slightly, in the middle of the larva. In comparison, when A8+tail is removed from position calculations, correlation between segment position shifts at the posterior of the larvae goes down significantly. This is true for the anterior part of the larva, when h+t are removed from position calculations.

**Figure S14.** This series of graphs are a continuation of Figure S13, as the distance between pairs of segments increase, from two to six segments apart. Notice that removal of h+t overall decreases correlation coefficients more than removal of A8+tail.

**Figure S15.** This series of graphs show the data represented in Figure S13 separately for each of the 12 *Drosophila* species. They show, for each species, how correlation coefficients change from the anterior to the posterior of the larvae, when relative segment position was calculated with all segments included (black) versus no A8+tail (red) or no h+t (blue).

**Table S1.** Experimental methods used for number of flies and days needed in a bottle to control population density of each species. Also includes, for each species, approximate embryonic developmental duration at 20°C and number of minutes needed in the 60°C oven for larvae to become straight.

**Table S2.** Lists the number of segments that are differentially positioned between pairs of species, and the divergence time between each pair in millions of years (Russo et al. 1995; Obbard et al. 2012; Russo et al. 2013).

**Table S3.** Lists the deviation of the position of each segment in each species from the “across-species” mean as well as total number of significant segment position changes for each species over all species-pair comparisons.

**File S1.** Describes the observed changes in the position of posterior border of denticle belts and denticle width between species.

**File S2.** Methods and results of the phylogenetic analysis.

**File S3.** Description of the method for removal of A8+tail or h+t from segment position calculations, as well as some additional results from that analysis. In particular, this file highlights the effects of the new calculations on differences in segment position along the larva as well as the variation in segment position within a species. It also gives more detail on the differences in the size of A8+tail and h+t between species when all segments are included in segment position calculations.

**File S4.** Describes correlation between changes in the position of adjacent segments along the anterior-posterior axis for each species, highlighting species-specific patterns.

## Notes

#### Summary of Updates

Introduction and Discussion revised to clarify points on robustness of segment patterning and within and between species differences in segment patterning. Results section revised to clarify points on changes in the position of the tail segments. Some figures and supplemental files have also been revised in accordance with the changes in the text.

https://doi.org/10.6084/m9.figshare.8085206

https://doi.org/10.6084/m9.figshare.8170787

https://github.com/joelatallah/larval_imaging

