## Supplementary material for "Evolution of Larval Segment Position across 12 *Drosophila* Species": FigS: Figure_S10_combined.pdf

Supplementary Figure 10

D.melanogaster Correlation score vs Pairwise distance

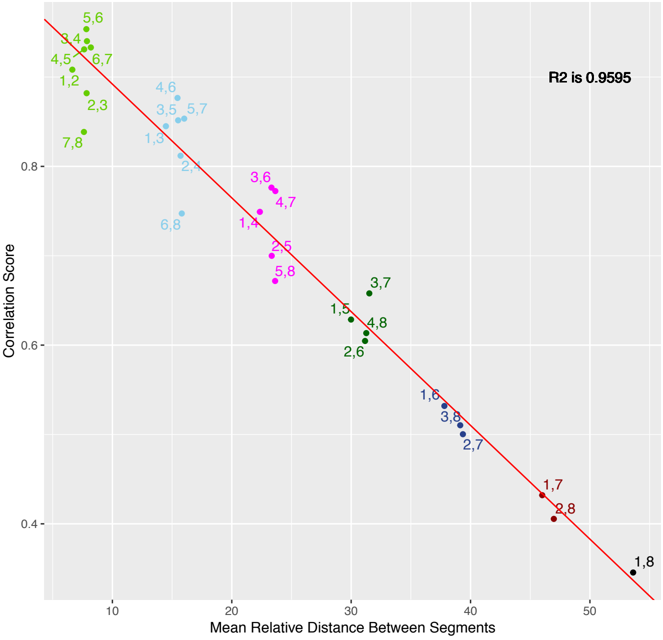


Supplementary Figure 10

D.simulans Correlation score vs Pairwise distance

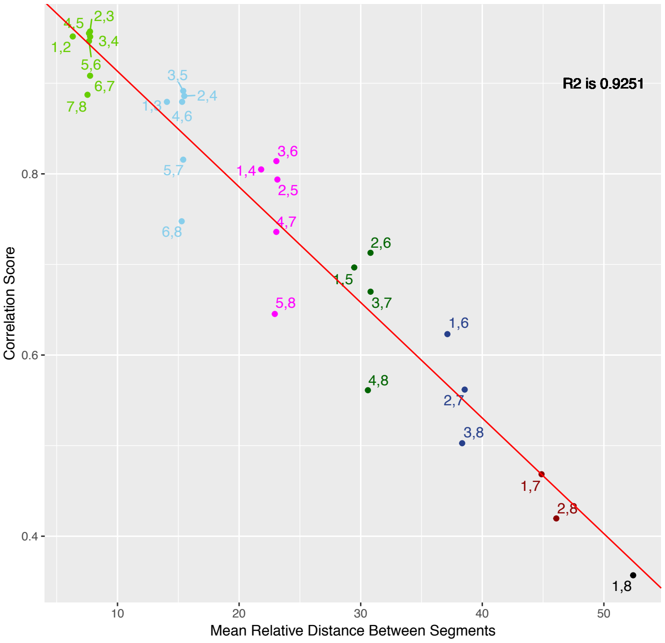


Supplementary Figure 10

D.sechellia Correlation score vs Pairwise distance

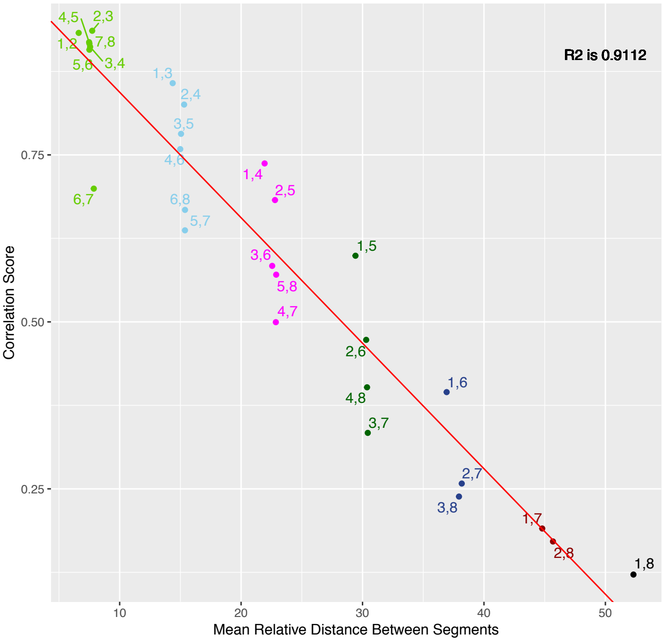


Supplementary Figure 10

D.yakuba Correlation score vs Pairwise distance

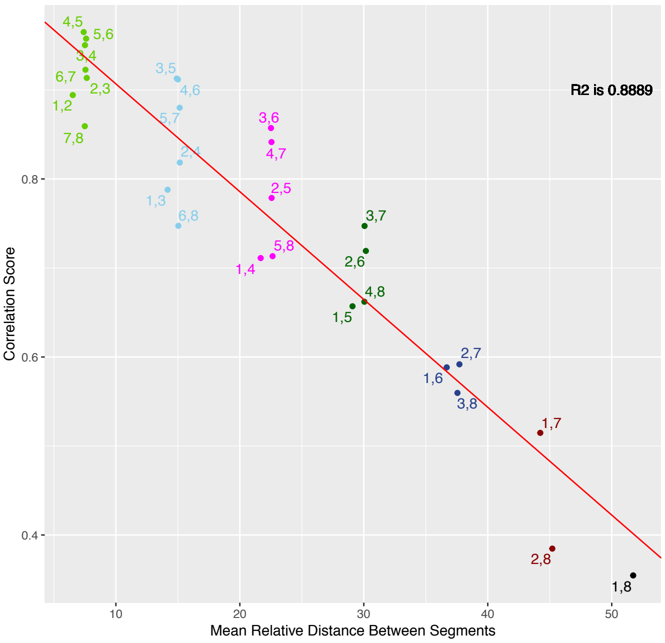


Supplementary Figure 10

D.santomea Correlation score vs Pairwise distance

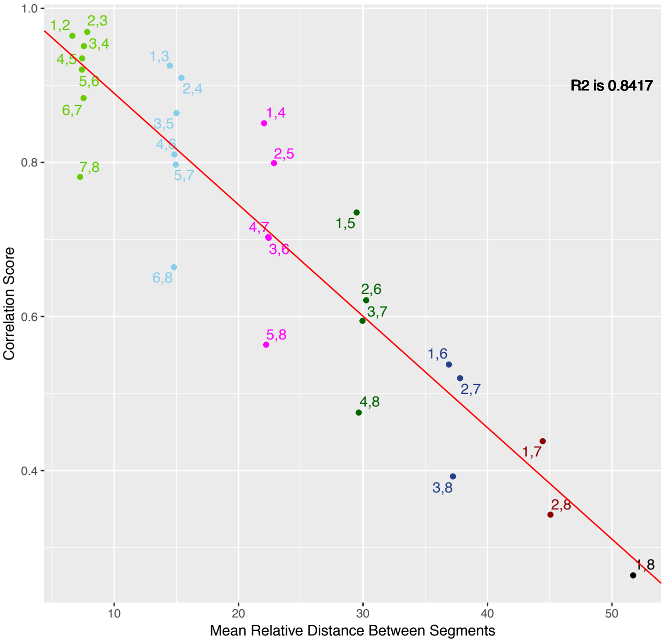


Supplementary Figure 10

D.erecta Correlation score vs Pairwise distance

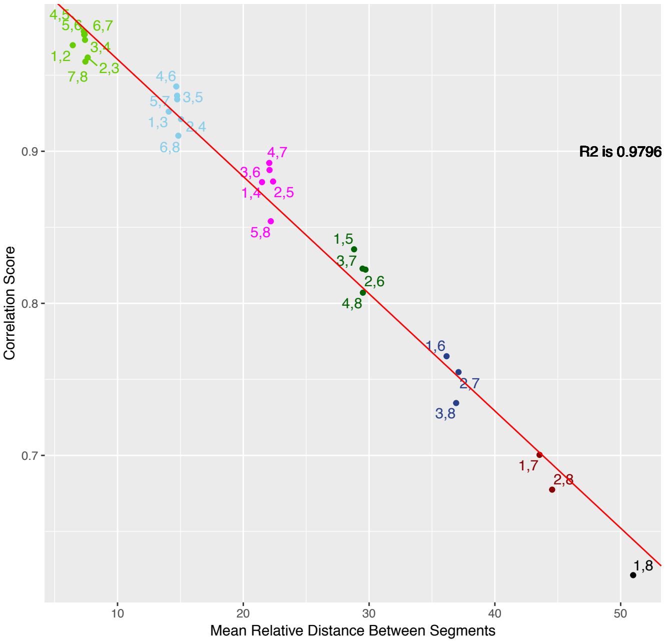

Supplementary Figure 10

D.ananassae Correlation score vs Pairwise distance

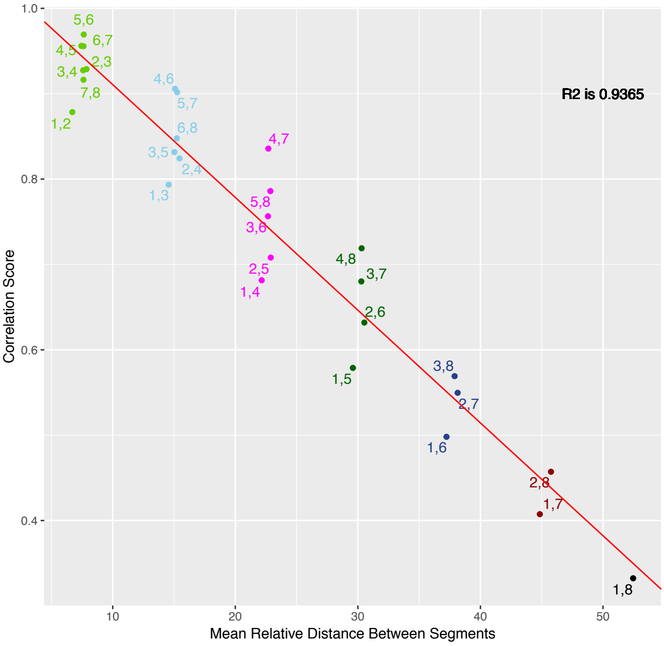


Supplementary Figure 10

D.pseudoobscura Correlation score vs Pairwise distance

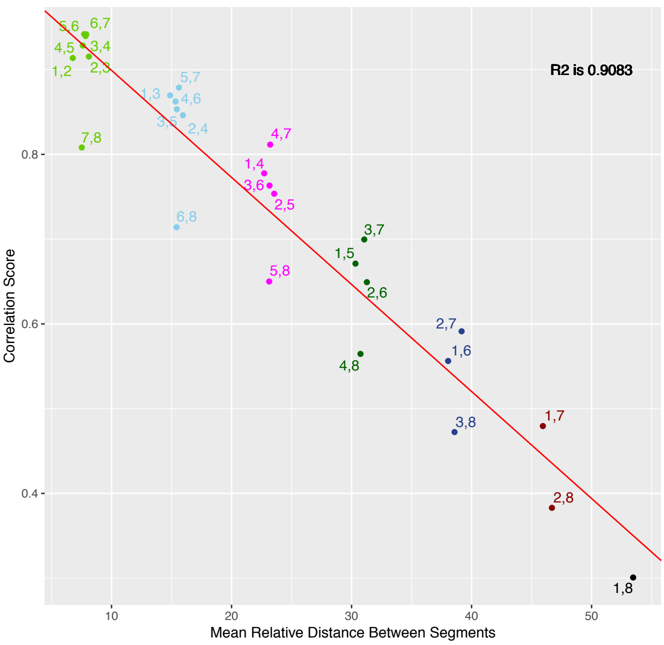


Supplementary Figure 10

D.persimilis Correlation score vs Pairwise distance

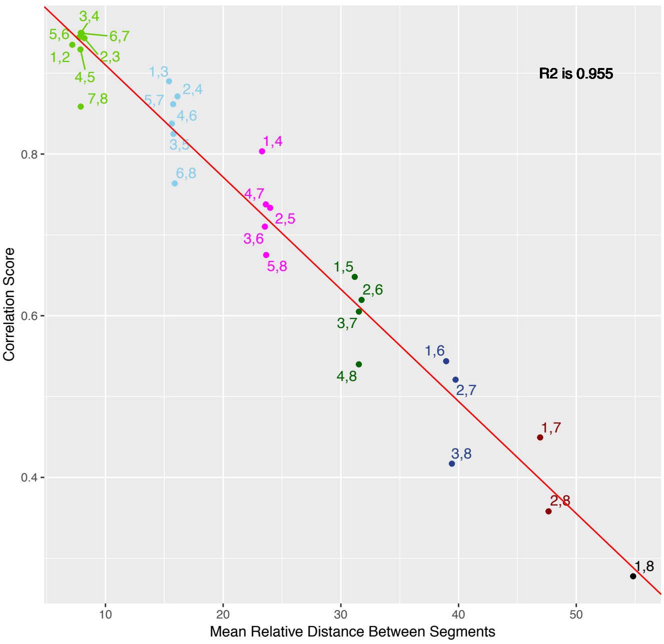


Supplementary Figure 10

D.willistoni Correlation score vs Pairwise distance

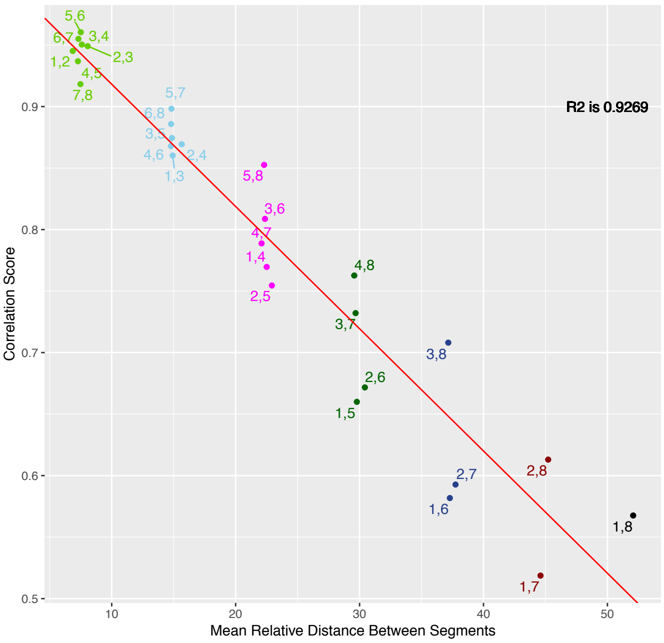


Supplementary Figure 10

D.mojavensis Correlation score vs Pairwise distance

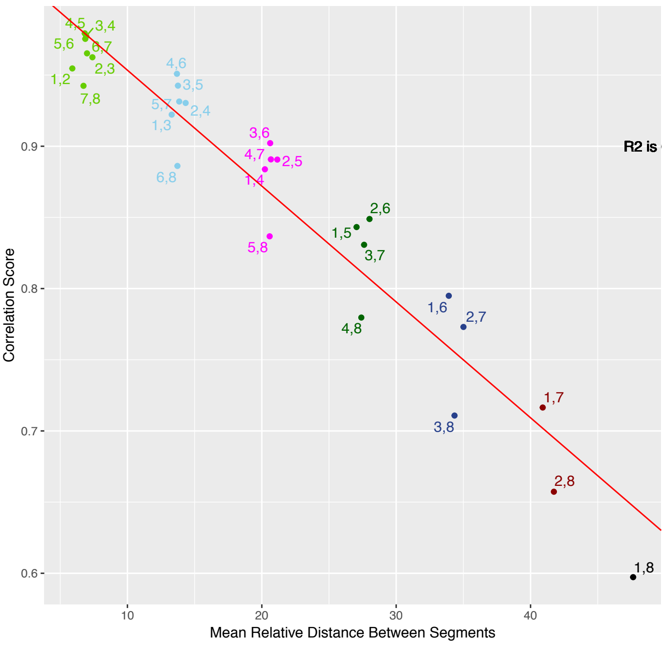


Supplementary Figure 10

D.virilis Correlation score vs Pairwise distance

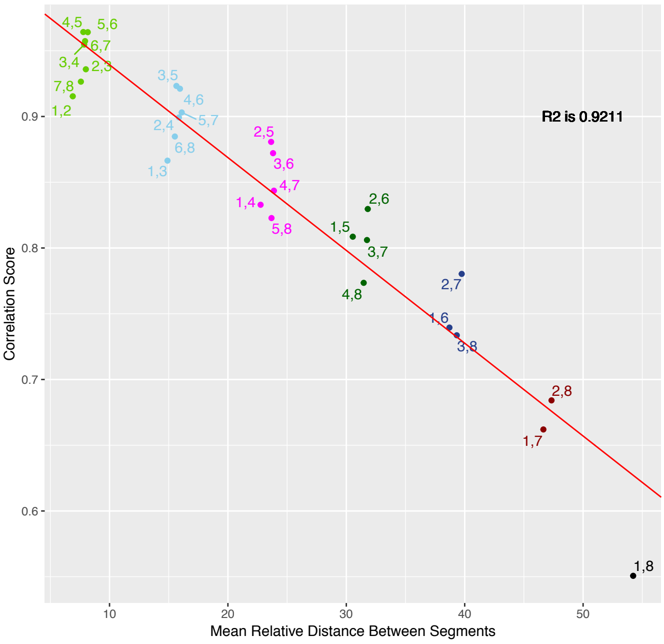
