## Supplementary material for "Evolution of Larval Segment Position across 12 *Drosophila* Species": FigS: Figure_S11_combined.pdf

Supplementary Figure 11

D.melanogaster Correlation Score Heatmap

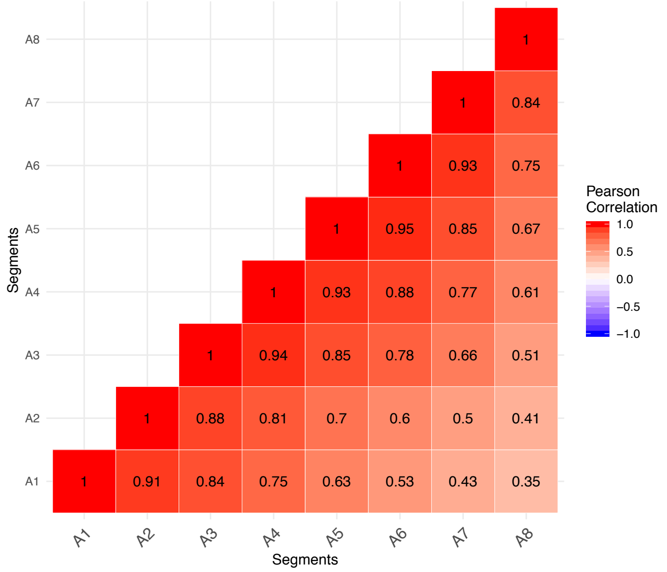


Supplementary Figure 11

D.simulans Correlation Score Heatmap

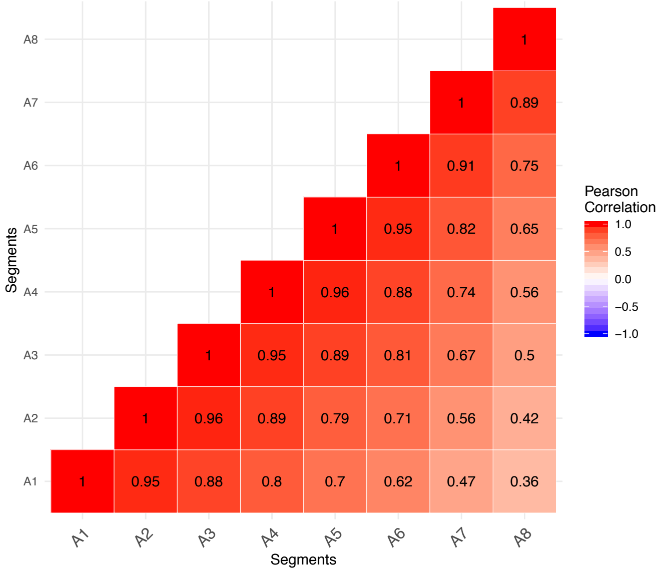


Supplementary Figure 11

D.sechellia Correlation Score Heatmap

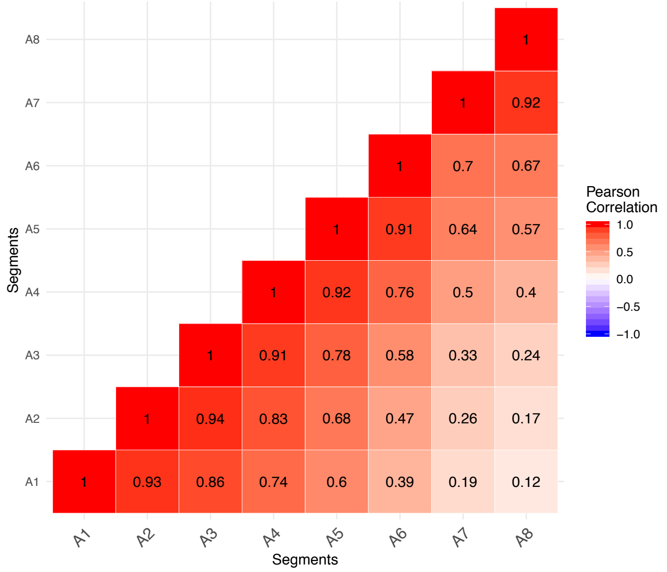


Supplementary Figure 11

D.yakuba Correlation Score Heatmap

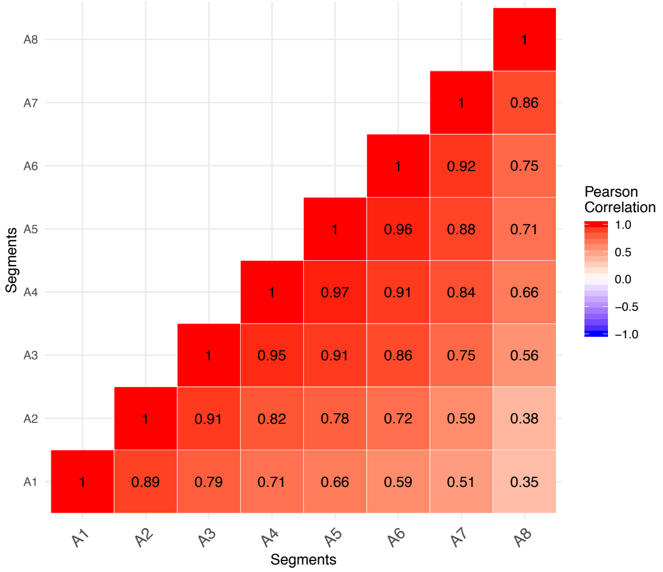


Supplementary Figure 11

D.santomea Correlation Score Heatmap

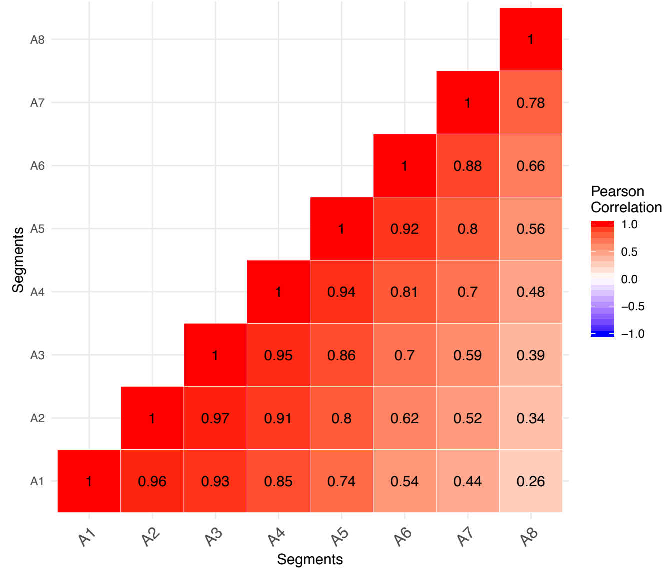


Supplementary Figure 11

D.erecta Correlation Score Heatmap

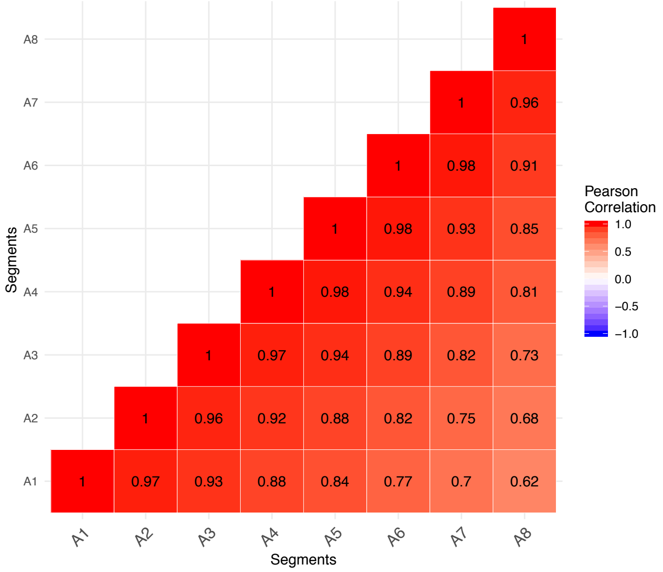

Supplementary Figure 11

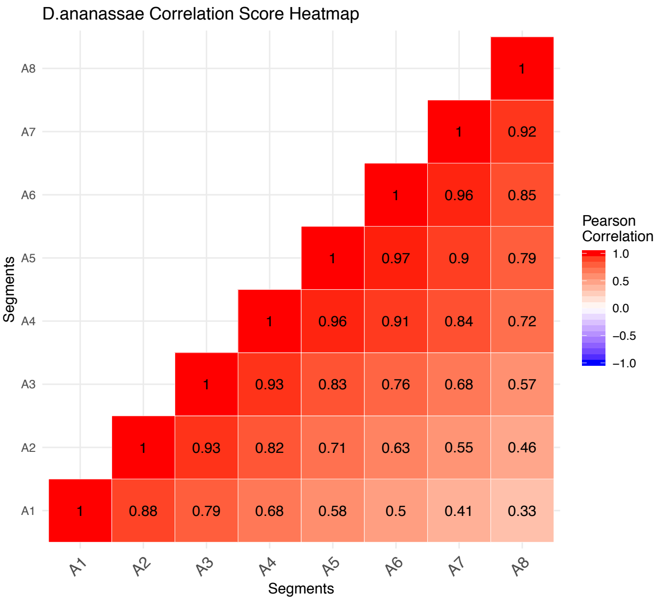


Supplementary Figure 11

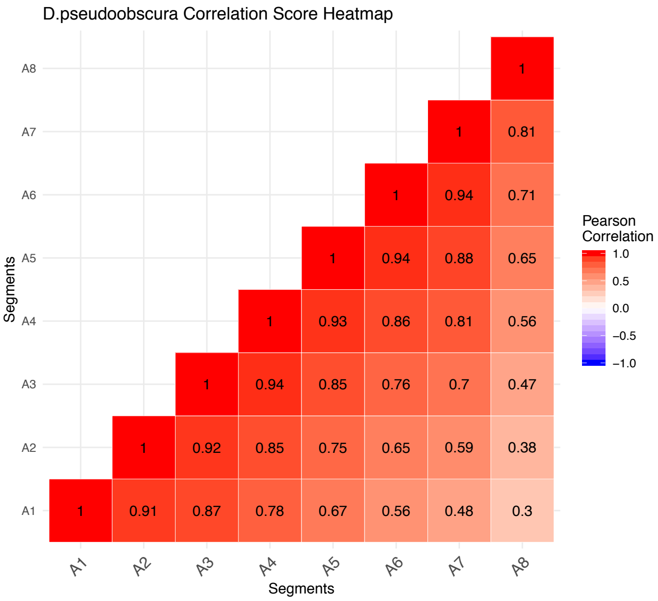


Supplementary Figure 11

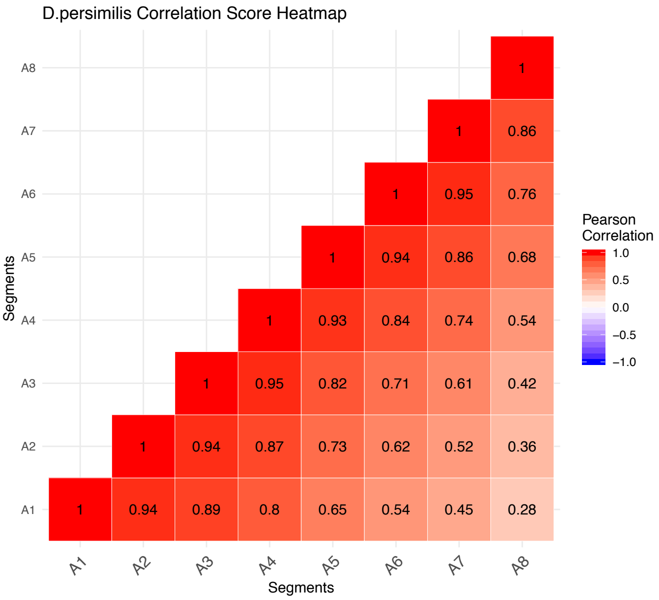


Supplementary Figure 11

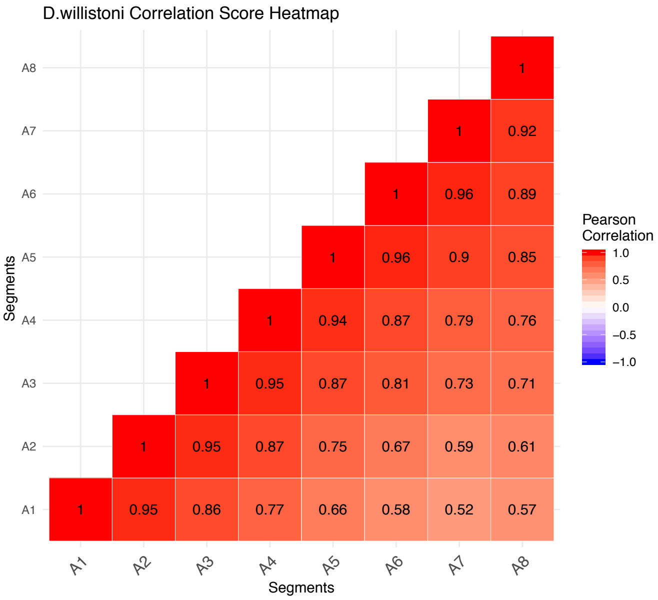


Supplementary Figure 11

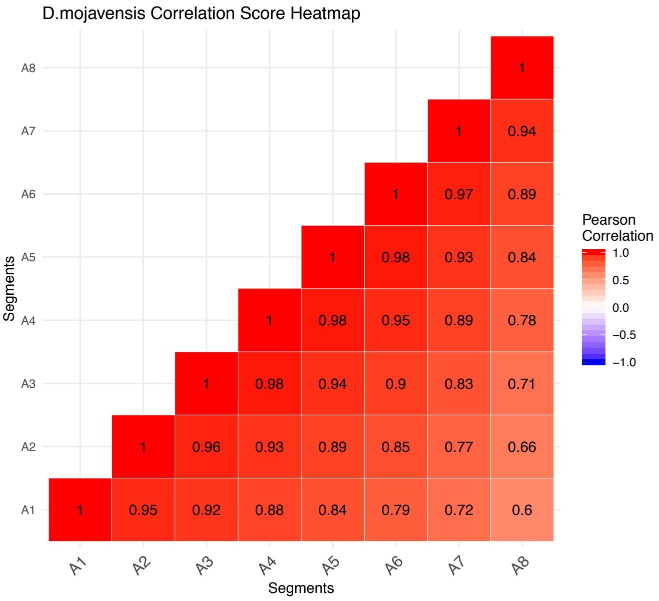


Supplementary Figure 11

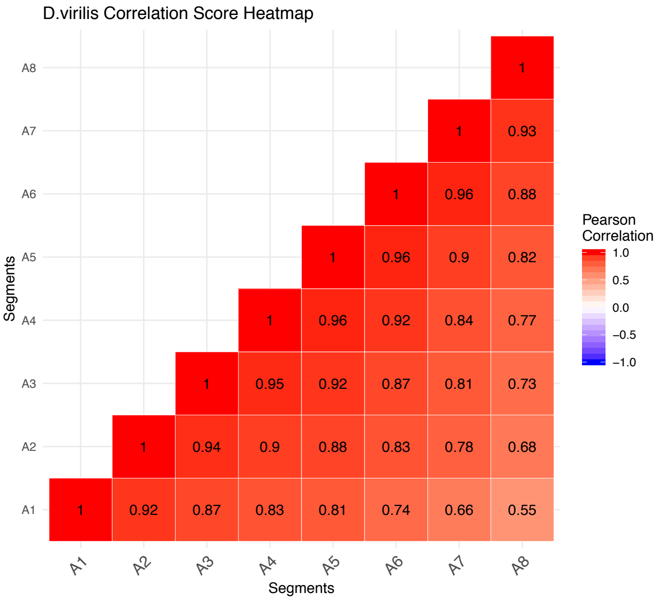
