## Supplementary material for "Evolution of Larval Segment Position across 12 *Drosophila* Species": FigS: Figure_S12_combinedinapage.pdf

Supplementary Figure 12

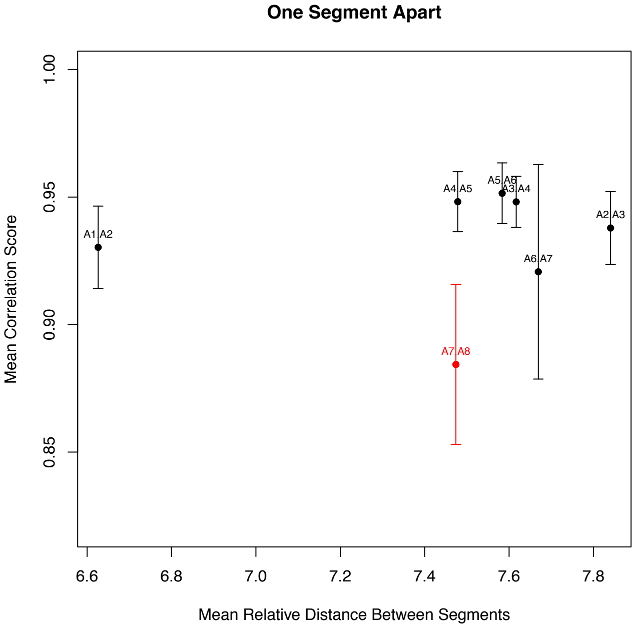


Supplementary Figure 12

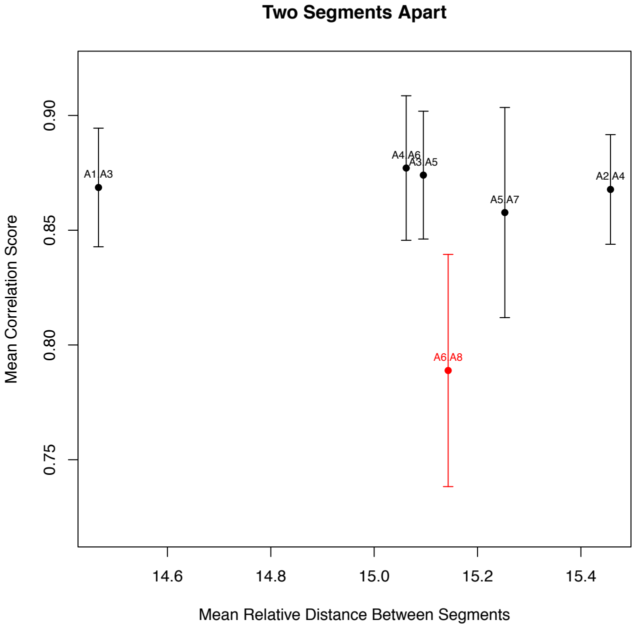


Supplementary Figure 12

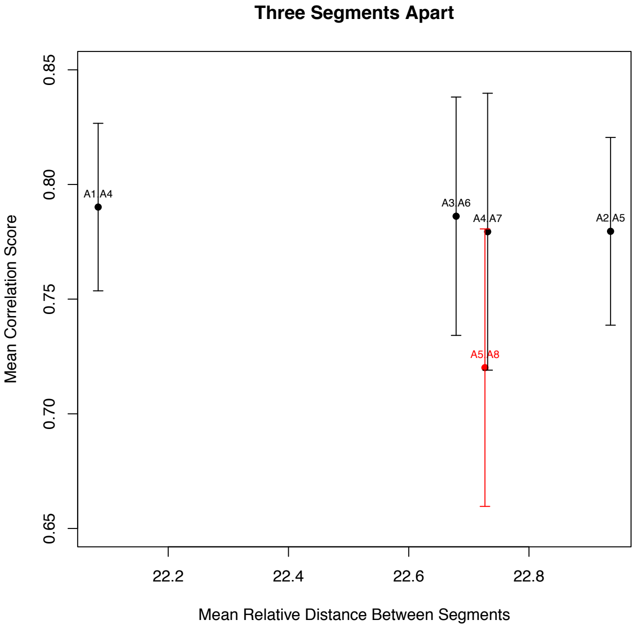


Supplementary Figure 12

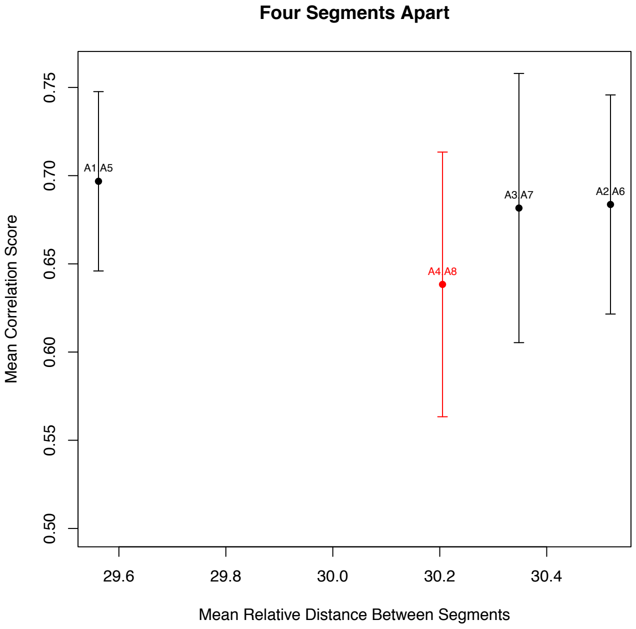


Supplementary Figure 12

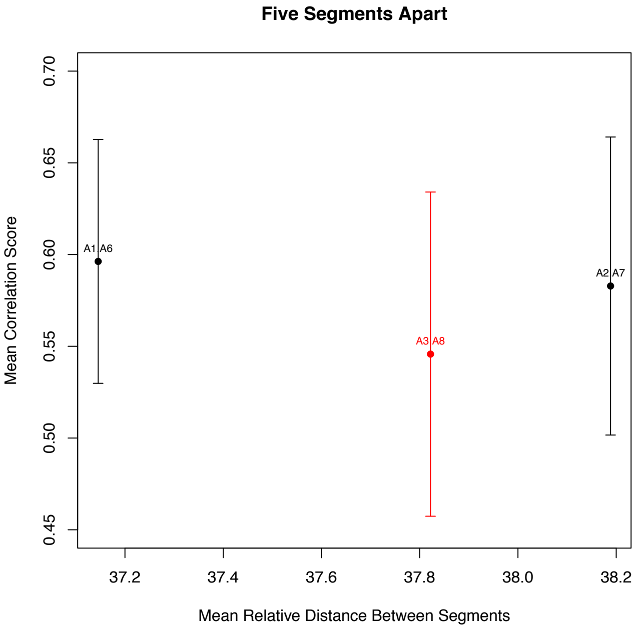


Supplementary Figure 12

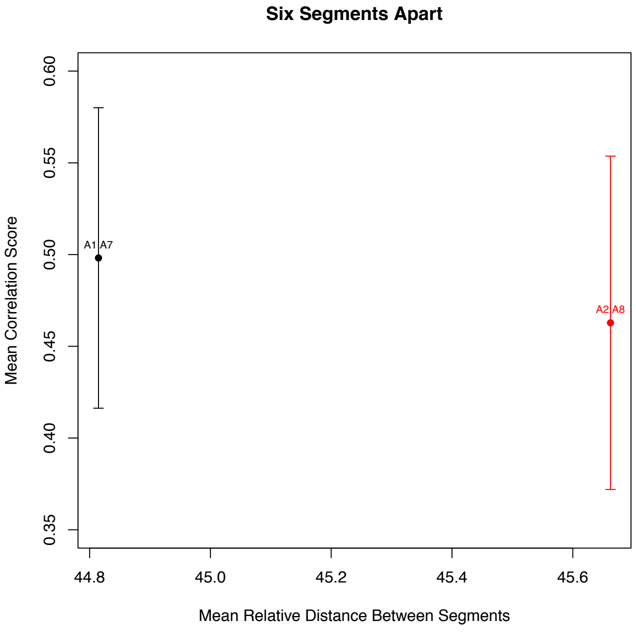
