## Supplementary material for "Evolution of Larval Segment Position across 12 *Drosophila* Species": FigS: Figure_S14_CombinedOnePage.pdf

Supplementary Figure 14

### Two Segments Apart

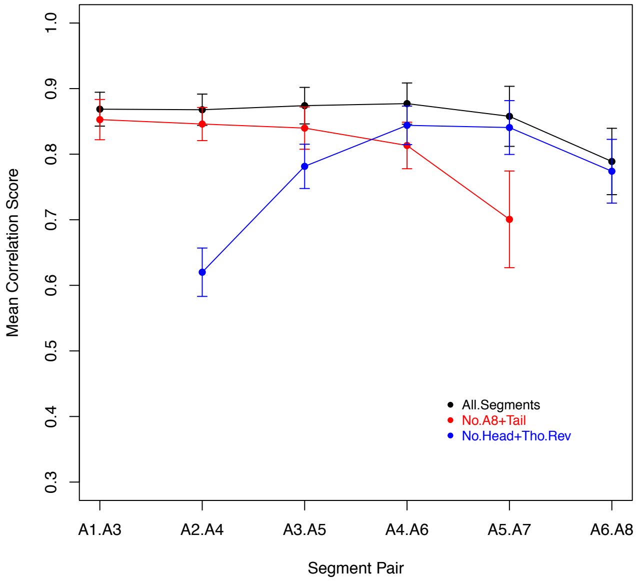


Supplementary Figure 14

### Three Segments Apart

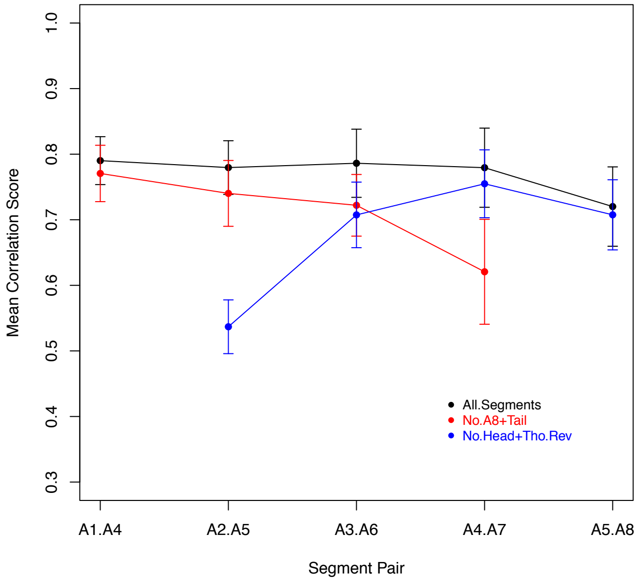


Supplementary Figure 14

### Four Segments Apart

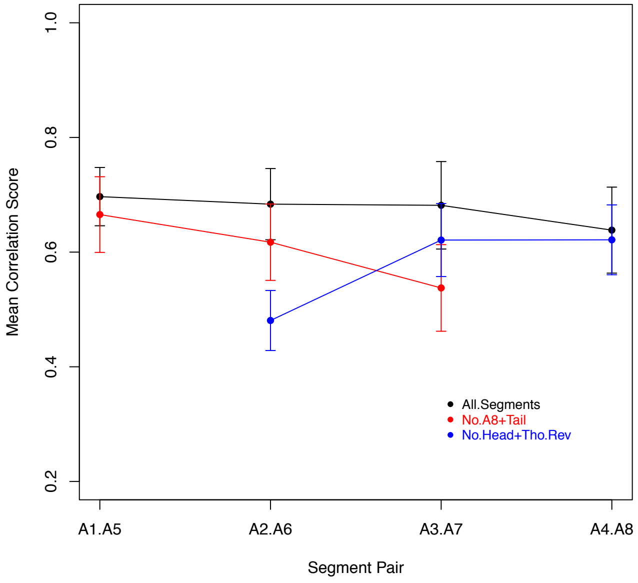


Supplementary Figure 14

### Five Segments Apart

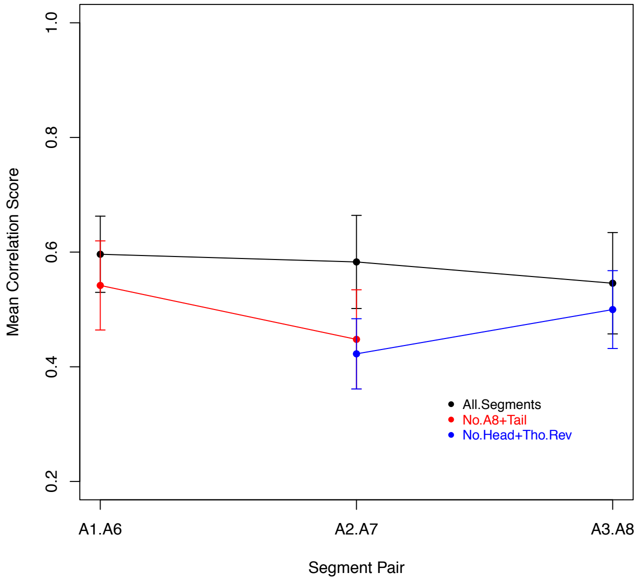


Supplementary Figure 14

### Six Segments Apart

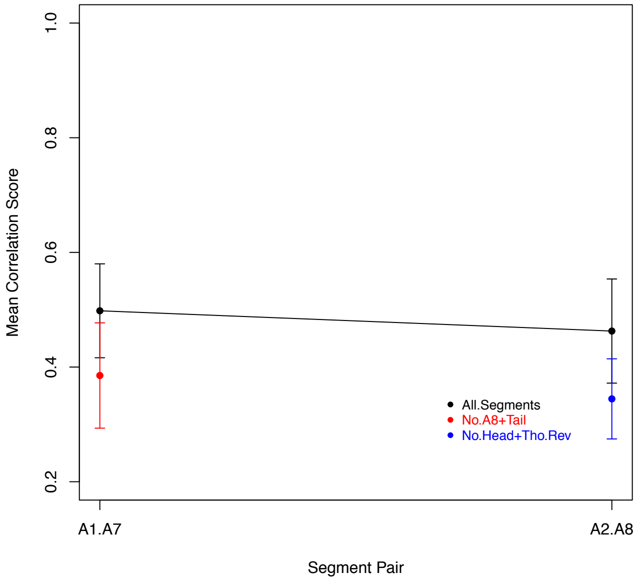
