## Supplementary material for "Evolution of Larval Segment Position across 12 *Drosophila* Species": FigS: Figure_S15_combined.pdf

Supplementary Figure 15

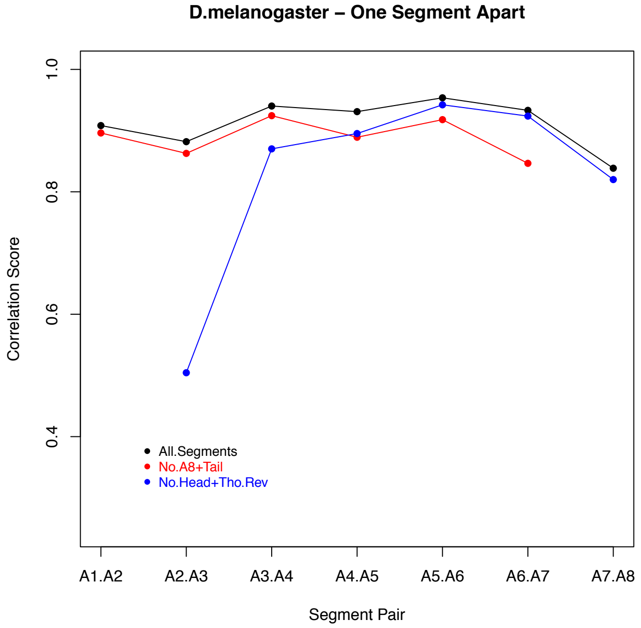


Supplementary Figure 15

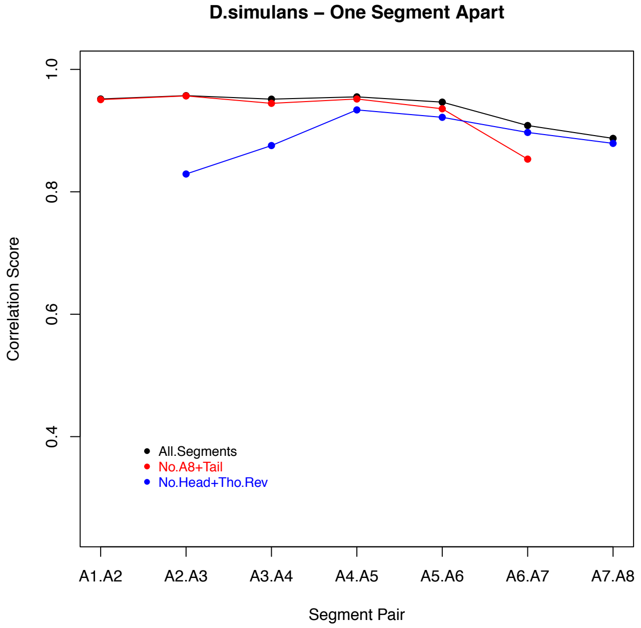


Supplementary Figure 15

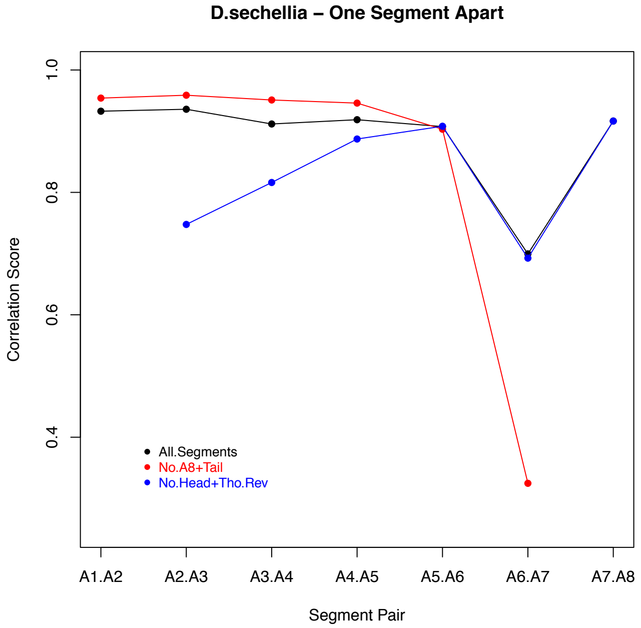


Supplementary Figure 15

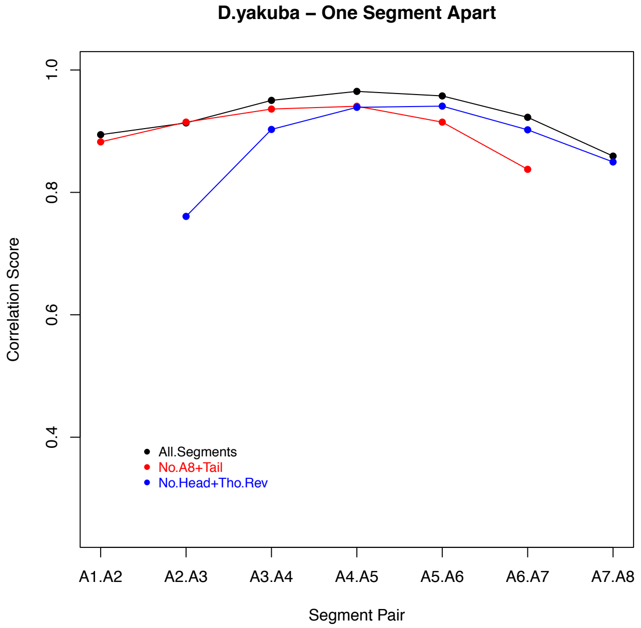


Supplementary Figure 15

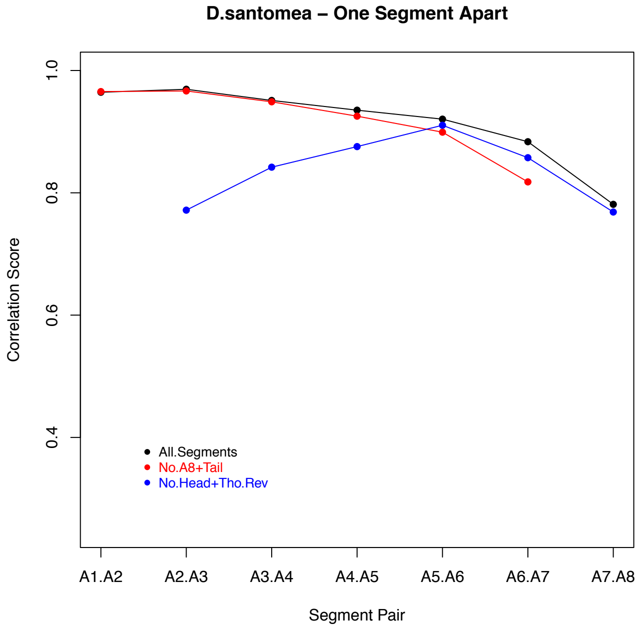


Supplementary Figure 15

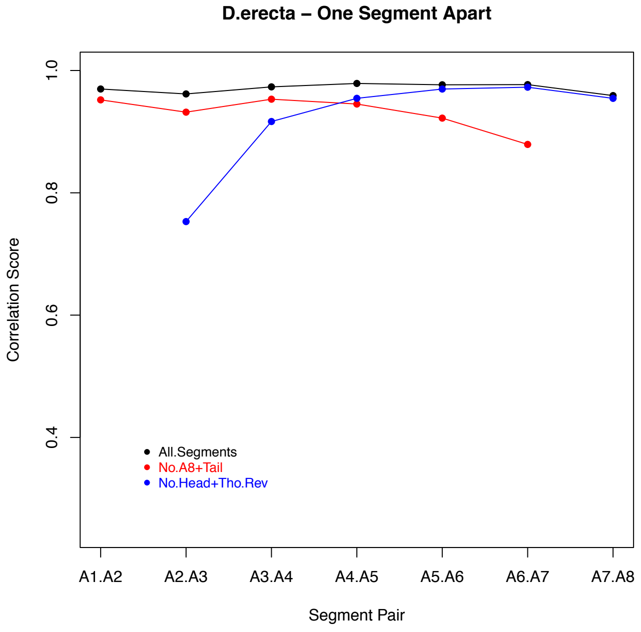

Supplementary Figure 15

**D.ananassae – One Segment Apart**

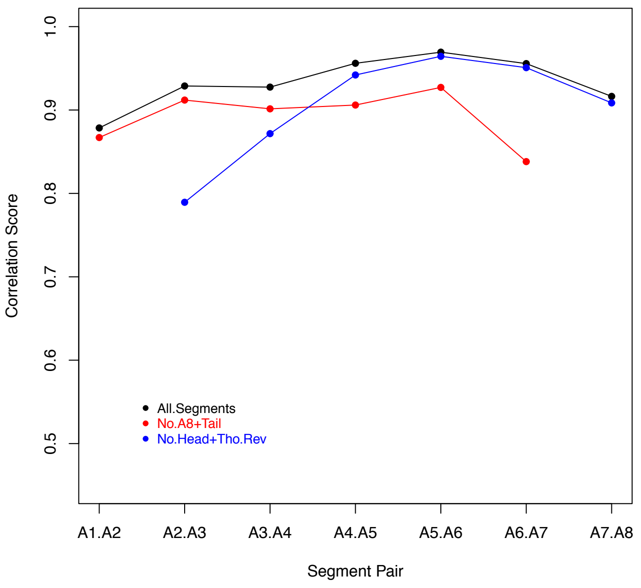


Supplementary Figure 15

**D.pseudoobscura – One Segment Apart**

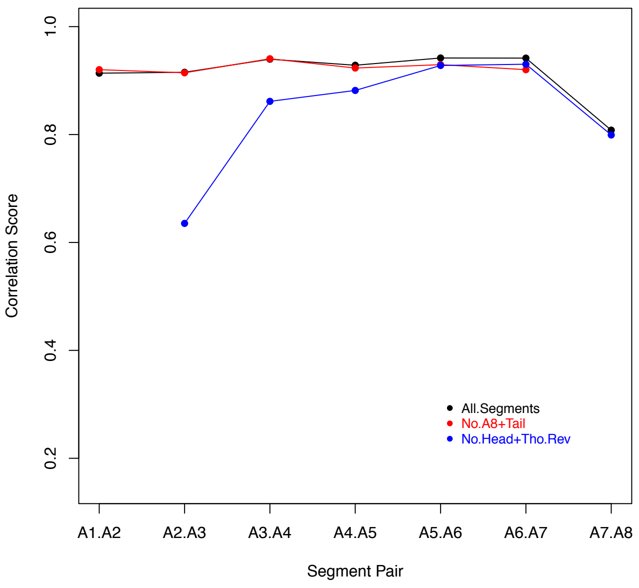


Supplementary Figure 15

**D.persimilis – One Segment Apart**

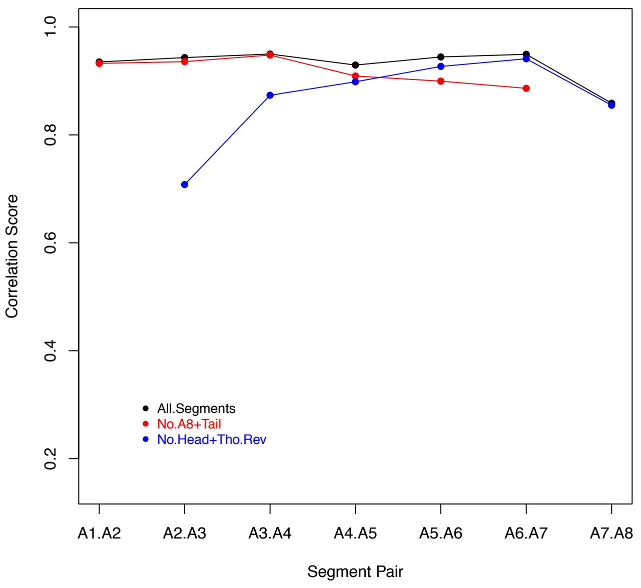


Supplementary Figure 15

**D.willistoni – One Segment Apart**

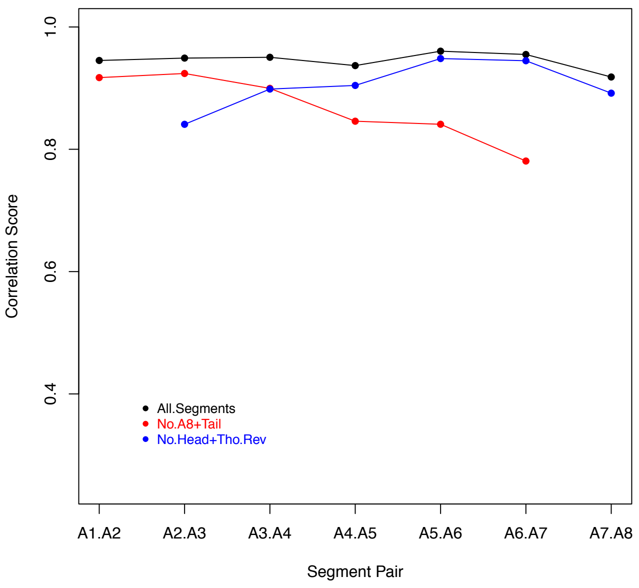


Supplementary Figure 15

**D.mojavensis – One Segment Apart**

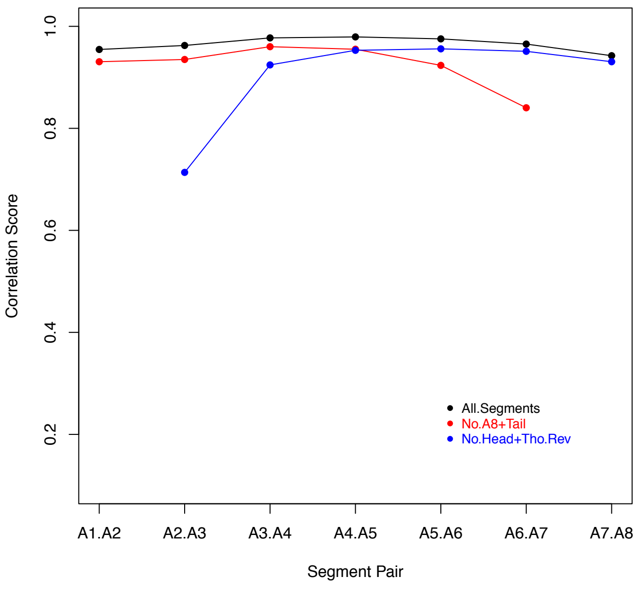


Supplementary Figure 15

**D.virilis – One Segment Apart**

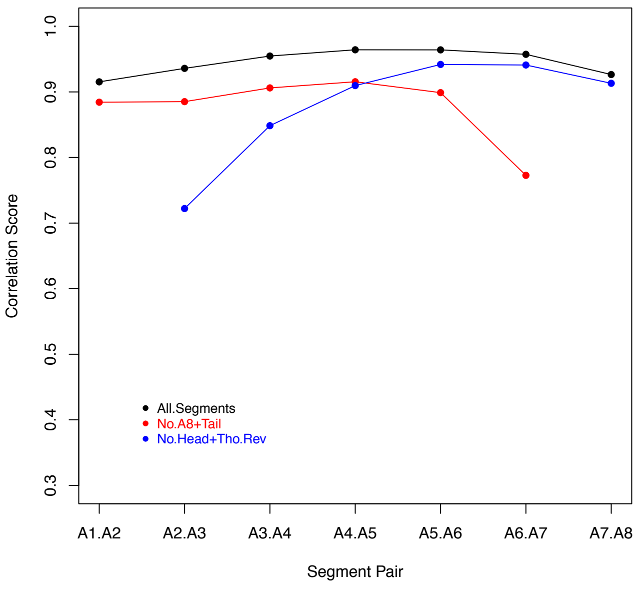
