## Supplementary material for "Evolution of Larval Segment Position across 12 *Drosophila* Species": FigS: File_S1.docx

**Supplementary File 1**

**Changes in the relative position of posterior border of denticle belts and changes in denticle width**

We also compared relative position of posterior border of denticle belts between species. Change in the position of the posterior border of a denticle belt is the sum of the shift in the anterior border plus the change in denticle belt size. Overall, the shift in the relative posterior position of denticle belts paralleled the shifts in the anterior positions (File S1 - Figure 1A). Thus, change in denticle belt size was not the most significant determinant of relative segment position. In 368 out of 410 significant shifts in segment position observed in species-pair comparisons, posterior border of the corresponding denticle belt also had an altered position. In all of these 368 position shifts, anterior and posterior border of the denticle belts shifted in the same direction (195 anterior shifts and 173 posterior shifts). The average magnitude of shift for each segment was a little higher for the posterior border of denticle belts as compared to the anterior border (File S1 - Figure 1B). This is likely due to the fact that posterior border of each denticle belt changes each time the anterior border shifts as well as when there is a change in the width of the corresponding denticle belt.

The anterior-posterior width of denticle belts also showed some variation between species (File S1 - Figure 1C). The width of denticle belt in segment A1 was the narrowest among all denticle belts in each species and also its width the most variable among species. The width Denticle belt for A8 was the second narrowest in all the species. The rest of the denticle belts were similar in size within a species along the anterior posterior axis, with some differences between species. *D. persimilis* stood out with having the widest denticle belts, except for the first one, among all the species.

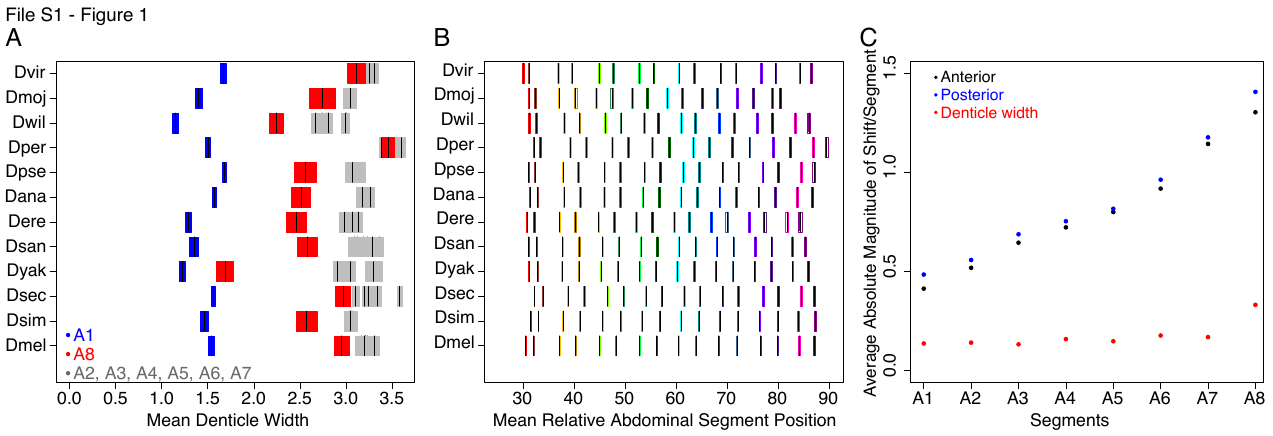
**File S1 – Figure 1.** (A) This graph shows differences in denticle width between 12 species. Blue represents segment A1, red represents segment A8 and gray represents the rest of the segments. Black bars represent the mean and the colored area around black bars are 95% confidence intervals. The y axis in this graph depicts 12 Drosophila species, the x axis shows mean denticle width. (B) This graph has the same format as Figure 2B, however in addition to differences in relative segment position (anterior border of denticle belts) it also shows changes in the posterior border of abdominal denticle belts between 12 species. The y axis depicts 12 species and x-axis shows mean relative abdominal segment position. Bars showing changes in posterior border of denticle belts are shaded darker than bars representing changes in the anterior borders. (C) This graph shows deviation from mean, averaged over all species, for the anterior (black), posterior (blue) and width (red) of denticle belts for each abdominal segment. The y axis show deviation from mean and the x axis depicts each abdominal segment.
