## Supplementary material for "Evolution of Larval Segment Position across 12 *Drosophila* Species": FigS: File_S2.pdf

### Supplemental Phylogenetic Methods

#### Contents

**S1 Phylogeny Estimation** **S2**
S1.1 Data . . . . . . . . . . S2
S1.2 Models . . . . . . . . . . S2
S1.3 Analyses . . . . . . . . . . S3
S1.4 Results . . . . . . . . . . S3

**S2 Morphological Evolution** **S5**
S2.1 Data . . . . . . . . . . S5
S2.2 Models . . . . . . . . . . S5
S2.3 Analyses . . . . . . . . . . S6
S2.4 Results . . . . . . . . . . S7

**S3 Joint Analysis** **S9**
S3.1 Data . . . . . . . . . . S9
S3.2 Models . . . . . . . . . . S9
S3.3 Analyses . . . . . . . . . . S9
S3.4 Results . . . . . . . . . . S10

### S1 Phylogeny Estimation

We adopted a Bayesian statistical approach to infer the phylogeny of our 12 *Drosophila* study species; we performed all phylogenetic analyses using the software package, RevBayes v.1.0.7 (Höhna et al. 2016). All of the RevBayes scripts used in this study are available in our Supplementary Data. We refer readers to those scripts for details regarding the hyperparameters and MCMC settings.

#### S1.1 Data

We used the 20 nuclear loci from Turelli et al. (2018) to estimate the *Drosophila* phylogeny. Specifically, these loci consist of: *aconitase*, *aldolase*, *bicoid*, *ebony*, *enolase*, *esc*, *g6pdh*, *glyp*, *glys*, *ninaE*, *pepck*, *pgi*, *pgm*, *pic*, *ptc*, *tpi*, *transaldolase*, *white*, *wingless* and *yellow*. We refer readers to Turelli et al. (2018) for specific details regarding the generation and alignment of these loci.

#### S1.2 Models

Our phylogenetic model consists of three components: (1) a substitution model that describes the process of evolution at each site in the alignment; (2) a node-age prior model that describes the distribution of branch-lengths, and; (3) a molecular branch-rate prior model that describes how rates of molecular evolution vary across the branches of the tree.

*Substitution model.*—We partitioned each of the 20 nuclear loci into first-, second-, and third-position sites. We assumed that each position for each locus evolved under an independent GTR+$\Gamma_4$ substitution model, for a total of 60 independent substitution models. We assumed conventional priors for each of the substitution-model parameters (see the Supplemental Data archive for details).

*Node-age prior model.*—We assumed that the phylogeny was generated by a sampled-birth-death node-age prior model. This model includes three parameters: (1) the speciation rate, $\lambda$, that describes the rate at which new species arise; (2) the extinction rate, $\mu$, that describe the rate at which species go extinct, and; (3) the sampling fraction $\rho$, that describes the fraction of extant species that are included in the sample. We fixed the sampling fraction, $\rho$, to be equal to the number of species in our sample divided by the number of described species in the clade, based on taxonomic information. Because we lack good fossil information regarding the age of the clade, we arbitrarily fixed the age of the root of the tree to one; consequently, all node ages and rates are measured relative to this fixed root age. We specified priors on the speciation- and extinction-rate parameters as described in our Supplemental Data archive.

*Molecular branch-rate prior models.*—The molecular branch-rate prior model describes how rates of molecular evolution vary among branches of the tree. Because divergence-time estimates are known to be sensitive to the assumed branch-rate prior model, we specified three different branch-rate prior models (Heath and Moore 2014): (1) an uncorrelated lognormal (UCLN) relaxed-molecular-clock model, which assumes that each branch draws its branch-rate independently from a lognormal distribution; (2) an uncorrelated exponential (UCED) relaxed-molecular-clock model, which assumes that each branch draws its branch-rate independently from an exponential distribution, and; (3) an autocorrelated lognormal (ACLN) relaxed molecular-clock model, which assumes that each branch draws its branch-rate from a lognormal distribution centered on the rate of the ancestral branch. Each of these branch-rate prior models contain hyperparameters that describe the average rate of evolution among branches and the degree of rate variation between branches; we estimate these hyperparameters—and the branch-specific rates of evolution—in our Bayesian model. The priors and hyperpriors specified in our molecular branch-rate prior models are detailed in the Supplementary Data archive.

#### S1.3 Analyses

*MCMC.*—We estimated divergence times for each combination of substitution model, node-age prior model, and branch-rate prior model described above. For each model, we performed four replicate MCMC analyses. We ensured that each independent MCMC simulation provided an adequate sample of the marginal posterior distribution of each parameter by computing the effective sample size (ESS) diagnostic computed using Tracer (Rambaut et al. 2018). We assessed convergence of the MCMC simulations to the target distribution (the joint posterior probability distribution) by comparing the marginal posterior distributions of each model parameter across the four replicates. We reran any analysis that failed to converge, or that failed to achieve ESS $> 500$ for each parameter.

#### S1.4 Results

After confirming that the four MCMC replicates for each branch-rate prior model converged, we then combined the samples from the four joint posterior distributions, and computed the the maximum *a posteriori* (MAP) summary tree from the resulting composite MCMC sample. This provided a single MAP summary tree for each of the three branch-rate prior models (Figures S1 to S3). While the tree topologies were consistent under the three branch-rate prior models, node-age estimates differed slightly. To ensure that our downstream morphological analyses (described below) were robust to the choice of the choice of branch-rate prior model (and the resulting differences in node-age estimates), we repeated each set of analyses on each of these three MAP trees.

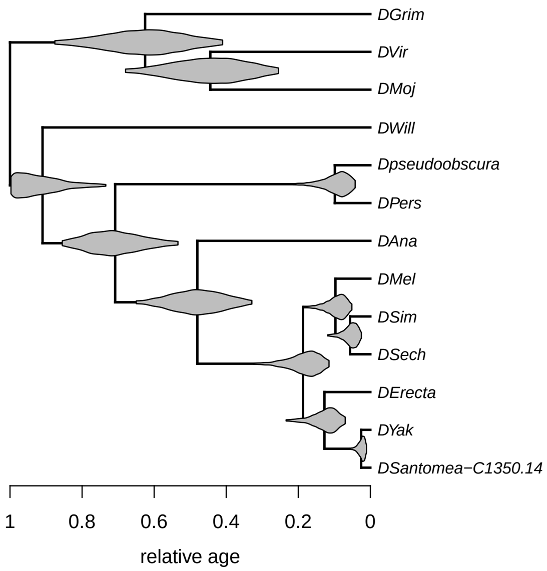


**Figure S1: The MAP phylogeny with relative divergence times inferred under the UCLN molecular branch-rate prior model.** Node ages represent the posterior-mean estimate of the divergence time for each node. Densities repsent the marginal posterior distribution (truncated in to the 95% credible interval) for each node.

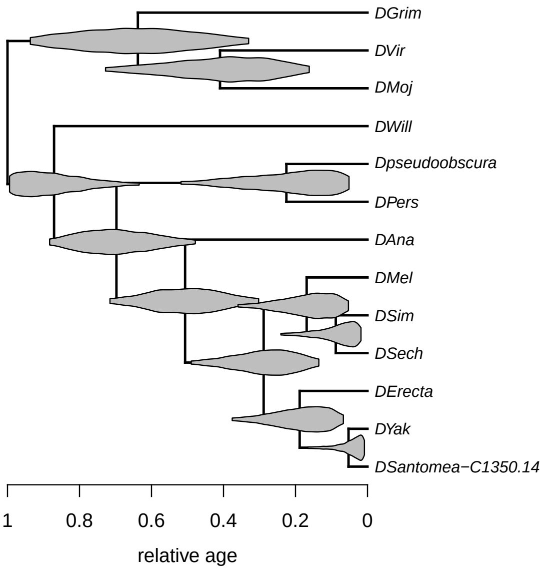


**Figure S2: The MAP phylogeny with relative divergence times inferred under the UCED molecular branch-rate prior model.** Node ages represent the posterior-mean estimate of the divergence time for each node. Densities repsent the marginal posterior distribution (truncated in to the 95% credible interval) for each node.

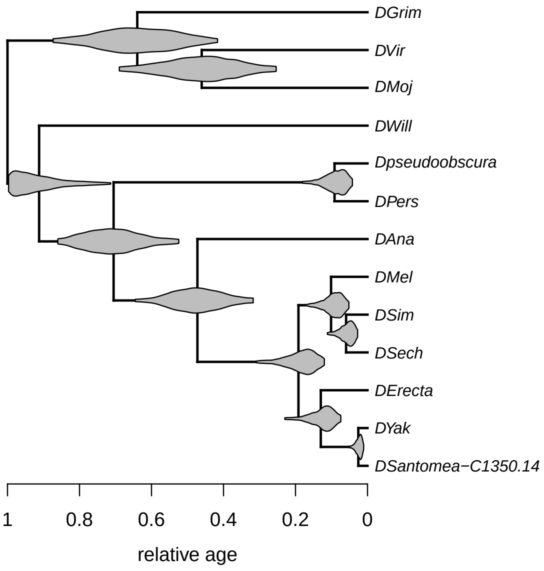


**Figure S3: The MAP phylogeny with relative divergence times inferred under the ACLN molecular branch-rate prior model.** Node ages represent the posterior-mean estimate of the divergence time for each node. Densities repsent the marginal posterior distribution (truncated in to the 95% credible interval) for each node.

### S2 Morphological Evolution

We assessed the relative fit of several candidate models of morphological evolution to the segment data using the phylogenies inferred from the previous step. We performed all morphological analyses using RevBayes v. 1.0.7 (Höhna et al. 2016). All of the RevBayes scripts used in these analyses are available in our Supplementary Data. We refer readers to those scripts for details regarding the hyperparameters and MCMC settings.

#### S2.1 Data

We constructed a data matrix of the eight segment positions, as well as total larval size, for each of our 12 *Drosophila* species. The Brownian-motion models of evolution we describe below assume that successive changes in each character are additive. This assumption has two practical consequences: (1) characters can become *negative* as a result of multiple decreases in size, and; (2) large characters may appear to evolve faster than small characters, as an additive increase of a given size will imply less *relative* change in larger characters. Therefore, we follow the convention of log-transforming our morphological data; this guarantees that the segment position can never become negative (because negative log-segment positions will still imply positive segment positions), and controls for the effect of absolute character size on evolutionary rates (because additive changes on the log scale are multiplicative on the absolute scale). We refer to the log-size of the $i^{\text{th}}$ segment for species $j$ as $x_j^i$ for each of the eight segments, and the log-total-body length (treated as the last character) for species $j$ as $x_j^9$. The collection of log-sizes across species comprises a "morphological alignment", which we refer to as $X$.

#### S2.2 Models

We model the evolution of the log-segment sizes using a phylogenetic multivariate Brownian-motion model. This model consists of two components: (1) an evolutionary model that describes how the relative rates of evolution vary among the characters, as well as the correlation structure among the characters, and; (2) a branch-rate prior model that describes how rates of morphological evolution vary across the branches of the tree.

*Evolutionary model.*—We assume that the segments evolve under a multivariate Brownian-motion (mvBM) model (Huelsenbeck and Rannala 2003; Lartillot and Poujol 2010). This model is comprised of two sets of parameters: $\boldsymbol{\sigma}^2$ and $R$. The relative-rate parameter $\boldsymbol{\sigma}^2$ is a vector that describes the rates of evolution among the continuous characters; $\sigma_i^2$ represents the rate for the $i^{\text{th}}$ character. (These rate parameters are *relative* because they are constrained to have a mean of 1; the absolute rate of evolution on a given branch in the phylogeny is determined by the morphological branch-rate prior model, described below). The correlation parameter $R$ is a matrix that describes the correlation between each pair of characters; $\rho_{ij}$ represents the degree to which evolutionary changes in $i$ and $j$ are correlated (*i.e.*, $\rho_{ij} = 1$ means that changes are perfectly correlated and $\rho_{ij} = 0$ means that changes are totally uncorrelated). Details of our prior on $\boldsymbol{\sigma}^2$ and $R$ are available in the Supplementary Data archive.

Together, the parameters $\boldsymbol{\sigma}^2$ and $R$ comprise the evolutionary variance-covariance matrix, $\Sigma$:

$$\Sigma = \begin{pmatrix} \sigma_1^2 & \sigma_1\sigma_2\rho_{12} & \cdots & \sigma_1\sigma_c\rho_{1c} \\ \sigma_2\sigma_1\rho_{12} & \sigma_2^2 & \cdots & \sigma_2\sigma_c\rho_{2c} \\ \vdots & \vdots & \ddots & \vdots \\ \sigma_c\sigma_1\rho_{1c} & \sigma_c\sigma_2\rho_{2c} & \cdots & \sigma_c^2 \end{pmatrix}.$$

*Morphological branch-rate prior models.*—We specify a morphological branch-rate prior model that is analogous to the molecular branch-rate prior model described above. In this case, the model describes how rates of *morphological* evolution vary among branches of the tree. We specified four different branch-rate prior models: (1) an uncorrelated lognormal (UCLN) relaxed-morphological-clock model, which assumes that each branch draws its branch-rate independently from a lognormal distribution; (2) an uncorrelated exponential (UCED) relaxed-morphological-clock model, which assumes that each branch draws its branch-rate independently from an exponential distribution, and; (2) an uncorrelated gamma (UCG) relaxed-morphological-clock model, which assumes that each branch draws its branch-rate independently from a gamma distribution, and; (3) an autocorrelated lognormal (ACLN) relaxed clock model, which assumes that each branch draws its branch-rate from a lognormal distribution centered on the rate of the ancestral branch. As with the relaxed molecular clock models, each of these morphological branch-rate prior models contains hyperparameters (*e.g.* the mean rate of evolution and the variance in the rate of evolution) that we estimate from the data. Details of our prior and hyperprior specifications for the morphological branch-rate prior models are available in the Supplementary Data archive.

#### S2.3 Analyses

*MCMC.*—We estimated the parameters of the mvBM model under each of the morphological branch-rate prior models described above. We repeated these analysis on each of the three MAP trees computed under the three molecular branch-rate prior models described in the previous section. For each model and MAP-tree combination, we performed four replicate MCMC analyses. We ensured that each independent MCMC simulation provided an adequate sample of the marginal posterior distribution of each parameter by computing the effective sample size (ESS) diagnostic computed using Tracer (Rambaut et al. 2018). We assessed convergence of the MCMC simulations to the target distribution (the joint posterior probability distribution) by comparing the marginal posterior distributions of each model parameter across the four replicates. We reran any analysis that failed to converge, or that failed to achieve ESS $> 500$ for each parameter.

*Marginal likelihood estimation.*—We compared the fit of the three candidate morphological branch-rate prior models to the segment data using Bayes factors (Jeffreys 1935). We first computed the marginal likelihood of the data under each morphological branch-rate prior model using stepping-stone sampling (Xie et al. 2010), implemented in `RevBayes`. For each model and MAP-tree combination, we performed four replicate stepping-stone analyses to confirm that our marginal-likelihood estimates were stable. We then compared the fit of each model by computing the Bayes factors between each pair of morphological branch-rate prior models; specifically, we computed $2 \ln \mathrm{BF}_{ij}$ between model $M_i$ and model $M_j$ as:

$$2 \ln \mathrm{BF}_{ij} = 2 \left[ \ln P(X \mid M_i) - \ln P(X \mid M_j) \right],$$

where $X$ is the morphological data, and $P(X \mid M_i)$ is the marginal likelihood of the morphological data under model $M_i$. A positive Bayes factor ($2 \ln \mathrm{BF}_{ij} > 0$) indicates support for model $M_i$, while a negative Bayes factor ($2 \ln \mathrm{BF}_{ij} < 0$) indicates support for model $M_j$ (Kass and Raftery 1995).

#### S2.4 Results

##### S2.4.1 Comparing the fit of alternative morphological branch-rate prior models

We assessed the relative fit of each morphological branch-rate model to the segment data. We repeated this model-comparison procedure for each of the three MAP trees estimated in the *Phylogeny Estimation* section. In all cases, the UCED and ACLN morphological branch-rate prior models were strongly disfavored compared to the best model (Tables S1 to S3). The marginal likelihoods for the UCLN and UCG models were very similar, with Bayes factors indicating equivocal support for either as the preferred model ($|2 \ln \mathrm{BF}| < 2$).

**Table S1:** Marginal likelihoods for each morphological branch-rate prior model with the UCLN MAP tree. Models are sorted from the highest (best) to lowest (worst) marginal likelihood. The $2 \ln \mathrm{BF}$ is measured against the best model (*i.e.*, negative values indicate the degree of evidence *against* the given model).

| | Marginal Likelihood | | | | | |
|---|---|---|---|---|---|---|
| Branch-rate model | Run 1 | Run 2 | Run 3 | Run 4 | Avg. | $2 \ln \mathrm{BF}$ |
| UCG | 126.913 | 126.036 | 125.733 | 127.137 | 126.455 | 0 |
| UCLN | 125.667 | 125.806 | 126.243 | 126.341 | 126.014 | −0.882 |
| UCED | 119.384 | 119.841 | 119.290 | 119.317 | 119.458 | −13.994 |
| ACLN | 115.597 | 109.474 | 116.178 | 113.731 | 113.745 | −25.420 |

**Table S2:** Marginal likelihoods for each morphological branch-rate prior model with the UCED MAP tree. Models are sorted from the highest (best) to lowest (worst) marginal likelihood. The $2 \ln \mathrm{BF}$ is measured against the best model (*i.e.*, negative values indicate the degree of evidence *against* the given model).

| | Marginal Likelihood | | | | | |
|---|---|---|---|---|---|---|
| Branch-rate model | Run 1 | Run 2 | Run 3 | Run 4 | Avg. | $2 \ln \mathrm{BF}$ |
| UCG | 127.873 | 127.903 | 125.971 | 127.740 | 127.372 | 0 |
| UCLN | 126.938 | 127.377 | 126.943 | 127.509 | 127.192 | −0.360 |
| UCED | 123.661 | 123.515 | 123.550 | 123.568 | 123.574 | −7.596 |
| ACLN | 116.826 | 117.390 | 118.431 | 115.103 | 116.938 | −20.868 |

**Table S3:** Marginal likelihoods for each morphological branch-rate prior model with the ACLN MAP tree. Models are sorted from the highest (best) to lowest (worst) marginal likelihood. The $2 \ln \mathrm{BF}$ is measured against the best model (*i.e.*, negative values indicate the degree of evidence *against* the given model).

| | Marginal Likelihood | | | | | |
|---|---|---|---|---|---|---|
| Branch-rate model | Run 1 | Run 2 | Run 3 | Run 4 | Avg. | $2 \ln \mathrm{BF}$ |
| UCG | 126.819 | 127.575 | 126.483 | 127.634 | 127.128 | 0 |
| UCLN | 126.313 | 126.090 | 126.000 | 125.969 | 126.093 | −2.070 |
| UCED | 118.691 | 119.222 | 118.842 | 119.022 | 118.919 | −16.418 |
| ACLN | 114.319 | 114.765 | 109.800 | 114.287 | 113.294 | −27.668 |

##### S2.4.2 Comparing branch-rate estimates under the preferred branch-rate prior models

For all MAP trees, the UCLN and UCG provided very similar fits to the data, as demonstrated by similar marginal-likelihood estimates. To understand whether these models—which provide an effectively indistinguishable fit to the morphological data—provide qualitatively different inferences regarding rates of segment evolution, we compared the posterior-mean estimate of each branch-specific rate parameter between the two models. These comparisons reveal that the branch-rate estimates under these competing (and equivocal) models are qualitatively identical for al three of the MAP trees (Figure S4).

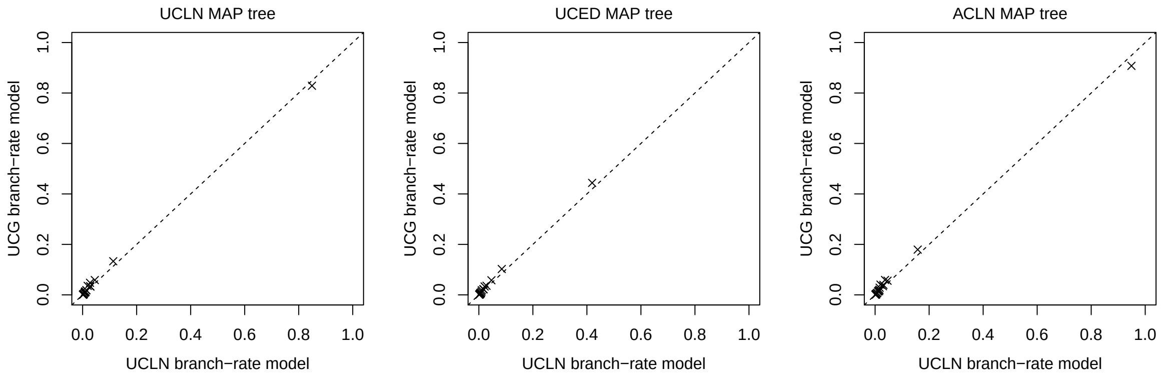


**Figure S4: Branch-rate estimates under the two (equivocally) preferred morphological branch-rate prior models.** We fit morphological branch-rate models using three MAP trees estimated under different molecular branch-rate prior models (UCLN, UCED, and ACLN; left, middle, and right panels, respectively). For each MAP tree, we compare the posterior-mean rate of evolution for branch $i$ under the UCLN model (x-axis) against the corresponding branch-rate estimate under the UCG model (y-axis). Branch-specific rate estimates effectively identical under the two morphological branch-rate models. By contrast, the branch-specific rates appear to vary between the alternative MAP trees.

### S3 Joint Analysis

We performed joint analyses—*i.e.*, where we simultaneously inferred the phylogeny, branch lengths, the parameters of the model of molecular evolution, and the parameters of the model of morphological evolution—from the combined molecular and morphological data. Analyzing the molecular and morphological data under a joint model allows us to average inferences about segment evolution over uncertainty in the tree topology, branch lengths, and substitution-model parameters.

For these analyses, we also estimated the posterior distribution of ancestral states of each segment size for each node in the tree. These ancestral-state estimates were used to compute the focal parameters: the rate of change in the *relative-segment size* along each branch in the tree, and the rate of change in the *relative-segment position* along each branch in the tree.

All of our joint analyses were performed using RevBayes v. 1.0.7 (Höhna et al. 2016). All of the RevBayes scripts we used to perform these available in our Supplementary Data. We refer readers to those scripts for details regarding the hyperparameters and MCMC settings.

#### S3.1 Data

For these joint analyses, we used the molecular alignments described in the *Phylogeny Estimation* section, and the morphological data described in the *Morphological Evolution* section, respectively.

#### S3.2 Models

For the molecular data, we used the substitution models, node-age prior models, and all three molecular branch-rate prior models, as described in the *Phylogeny Estimation* section. For the morphological data, we used the multivariate Brownian motion model and the uncorrelated lognormal morphological branch-rate prior models, as described in the *Morphological Evolution* section. This resulted in a total of three models, which differed by the assumed molecular branch-rate prior model.

*Ancestral-state estimation.*—We used data augmentation to generate samples from the posterior distribution of ancestral states for each larval segment for each node in the phylogeny (Lartillot and Poujol 2010). Briefly, this technique involves including the (log) segment sizes as parameter of our Bayesian model, and then using Markov chain Monte Carlo to sample the posterior distribution of these parameters, just as we would with any other parameter in the model. We refer to the size of the $i^{\text{th}}$ segment for internal node $n$ as $x_n^i$.

#### S3.3 Analyses

*MCMC.*—We estimated the parameters under each joint model, as described above. For each joint model, we performed four replicate MCMC analyses. We ensured that each independent MCMC simulation provided an adequate sample of the marginal posterior distribution of each parameter by computing the effective sample size (ESS) diagnostic computed using Tracer (Rambaut et al. 2018). We assessed convergence of the MCMC simulations to the target distribution (the joint posterior probability distribution) by comparing the marginal posterior distributions of each model parameter across the four replicates. We reran any analysis that failed to converge, or that failed to achieve ESS $>$ 500 for each parameter.

#### S3.4 Results

##### S3.4.1 Phylogeny estimates under the joint model

After confirming that the four MCMC replicates for each joint model converged, we combined the samples from the four joint posterior distributions and compute the the maximum *a posteriori* (MAP) summary tree. This resulted in a single MAP summary tree for each joint model (Figures S5 to S7).

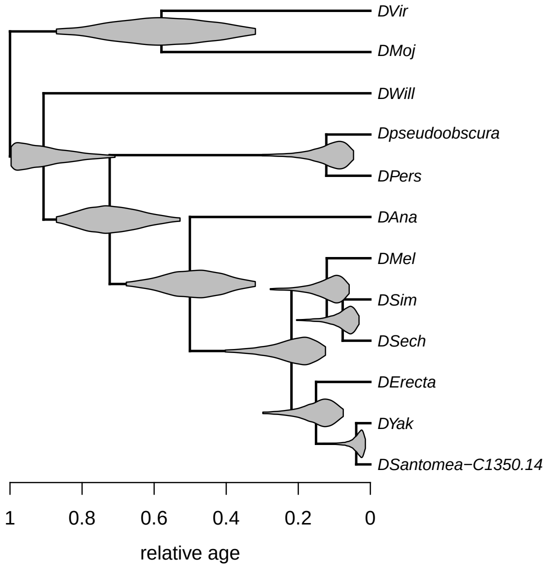


**Figure S5: The MAP phylogeny with relative divergence times inferred under the UCLN molecular branch-rate prior model.** Node ages represent the posterior-mean estimate of the divergence time for each node. Densities repsent the marginal posterior distribution (truncated in to the 95% credible interval) for each node.

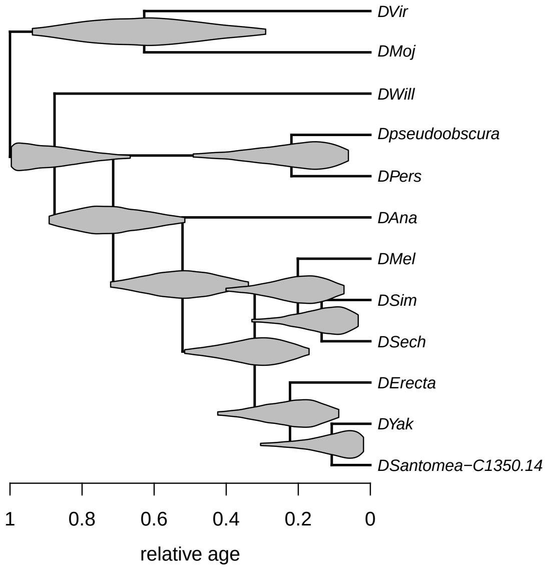


**Figure S6: The MAP phylogeny with relative divergence times inferred under the UCED molecular branch-rate prior model.** Node ages represent the posterior-mean estimate of the divergence time for each node. Densities repsent the marginal posterior distribution (truncated in to the 95% credible interval) for each node.

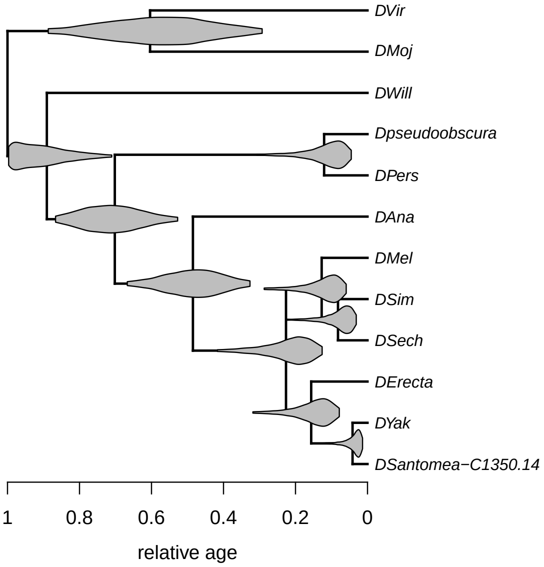


**Figure S7: The MAP phylogeny with relative divergence times inferred under the ACLN molecular branch-rate prior model.** Node ages represent the posterior-mean estimate of the divergence time for each node. Densities repsent the marginal posterior distribution (truncated in to the 95% credible interval) for each node.

##### S3.4.2 Variation in the rate of evolution among segments

The parameter $\sigma^2$ describes the relative rate of evolution for each segment: $\sigma_i^2$ is the relative rate of change of segment $i$ (compared to the rate of change for the "average" segment). The posterior estimates of $\sigma^2$ indicate that rates of evolution varies significantly among segments (Figures S8 to S10); notably, the eighth segment evolves much more quickly than the remaining segments.

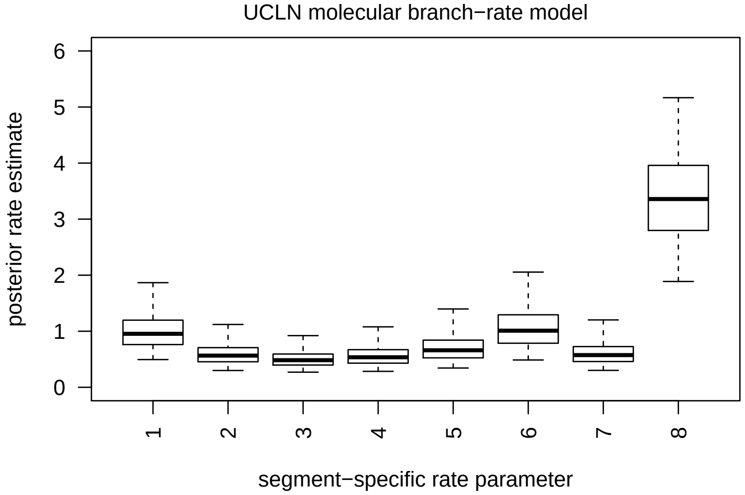


**Figure S8: Posterior distributions of segment-specific relative-rate parameters, $\sigma^2$, under the UCLN molecular-clock model.** Boxes represent the 50% credible interval for each posterior distribution, and whiskers represent the 95% credible interval.

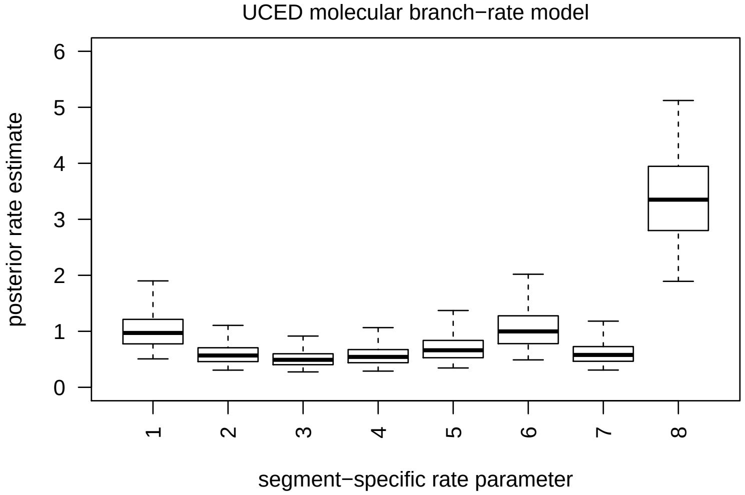


**Figure S9: Posterior distributions of segment-specific relative-rate parameters, $\sigma^2$, under the UCED molecular-clock model.** Boxes represent the 50% credible interval for each posterior distribution, and whiskers represent the 95% credible interval.

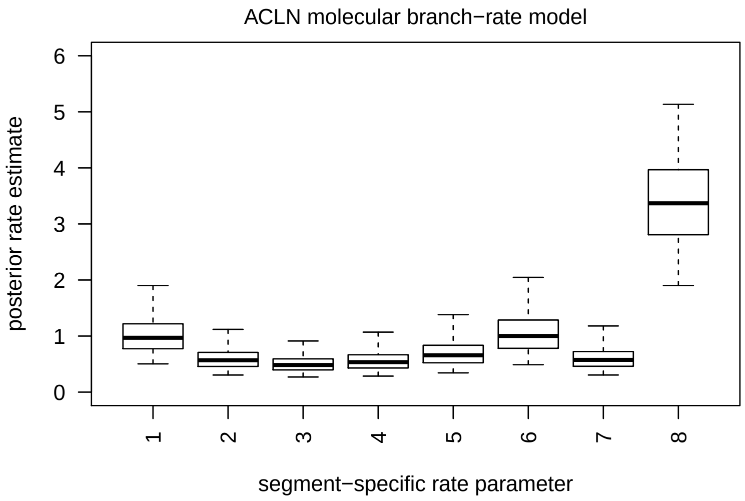


**Figure S10: Posterior distributions of segment-specific relative-rate parameters, $\sigma^2$, under the ACLN molecular-clock model.** Boxes represent the 50% credible interval for each posterior distribution, and whiskers represent the 95% credible interval.

##### S3.4.3 (Auto)correlation among segments

The correlation matrix $R$ describes evolutionary correlations between each pair of segments; the correlation between segment $i$ and segment $j$ is $\rho_{ij}$. To understand whether changes in segment size are correlated, we computed the posterior mean correlation matrix, $\widehat{R}$, for each analysis. The correlation coefficient between each pair of segments are positive (Figures S11 to S13, left panel); this indicates that segments tend to evolve in a consistent direction (among segments) during evolution.

To understand whether changes in segments are autocorrelated along the larva—*i.e.*, whether anatomically close segments demonstrate higher degrees of correlation relative to anatomically distant segments—we used the posterior-mean correlation matrix, $\widehat{R}$, to compute the posterior-mean *partial* correlation matrix, $\widehat{P}$. The partial correlation matrix measures the degree of correlation between each segment pair $i$ and $j$, controlling for their correlation induced through the other segments (Figures S11 to S13, right panel).

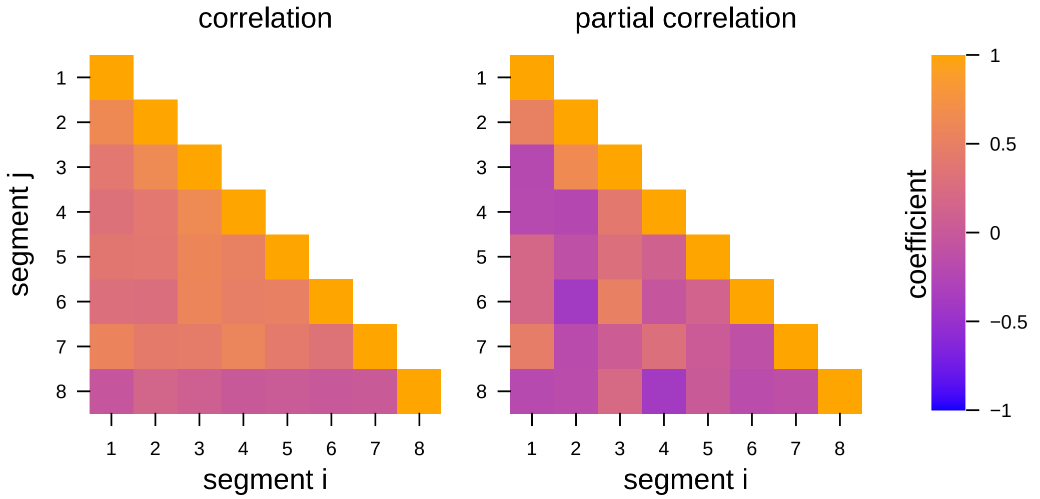


**Figure S11: Correlation coefficients between each pair of segments inferred under the UCLN molecular branch-rate model.** Correlation coefficients between each pair of segments (left panel). Partial correlation coefficients between each pair of segments (right panel).

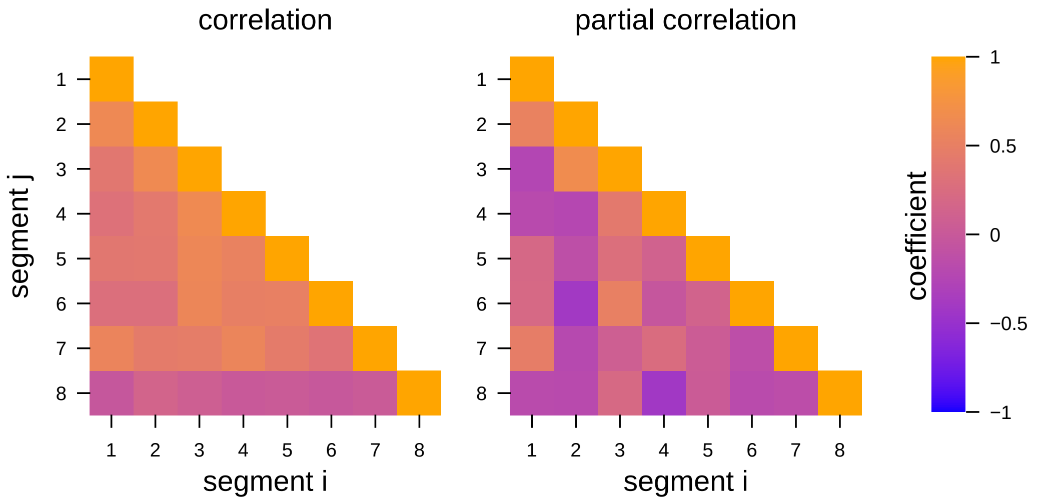


**Figure S12: Correlation coefficients between each pair of segments inferred under the UCED molecular branch-rate model.** Correlation coefficients between each pair of segments (left panel). Partial correlation coefficients between each pair of segments (right panel).

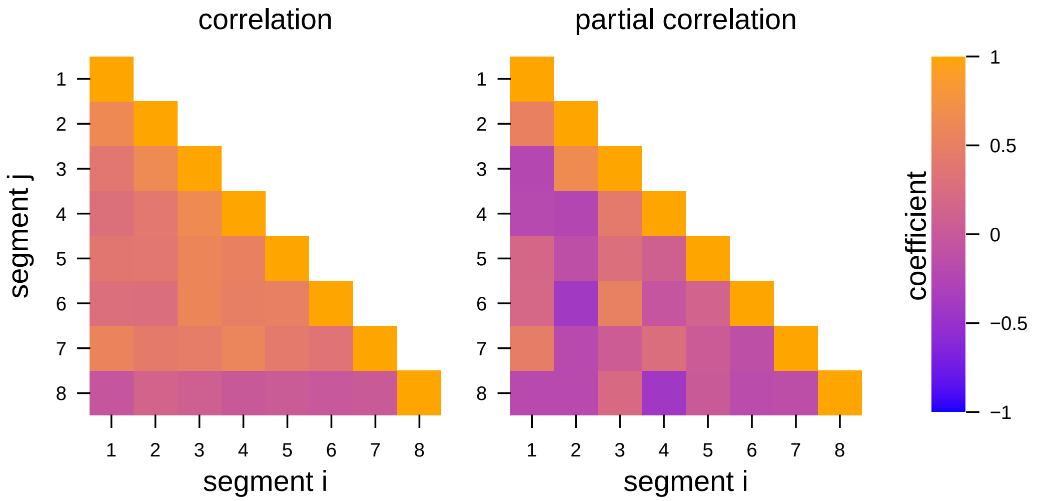


**Figure S13: Correlation coefficients between each pair of segments inferred under the ACLN molecular branch-rate model.** Correlation coefficients between each pair of segments (left panel). Partial correlation coefficients between each pair of segments (right panel).

##### S3.4.4 Variation in the rate of segment evolution across lineages

The morphological branch-rate model describes how rates of evolution vary among branches of the tree. Under this model, each branch in the tree is assigned a branch-specific rate parameter, $\beta^2$, that determines the rate of evolution for the continuous characters on that branch: the rate of evolution for character $i$ on branch $j$ is simply $\sigma_i^2 \times \beta_j^2$. The inferred posterior distributions of $\beta^2$ demonstrate that rates of segment evolution vary among branches of the phylogeny (Figures S14 to S16). The patterns of morphological rate variation among branches are qualitatively similar under each of the three molecular branch-rate prior models.

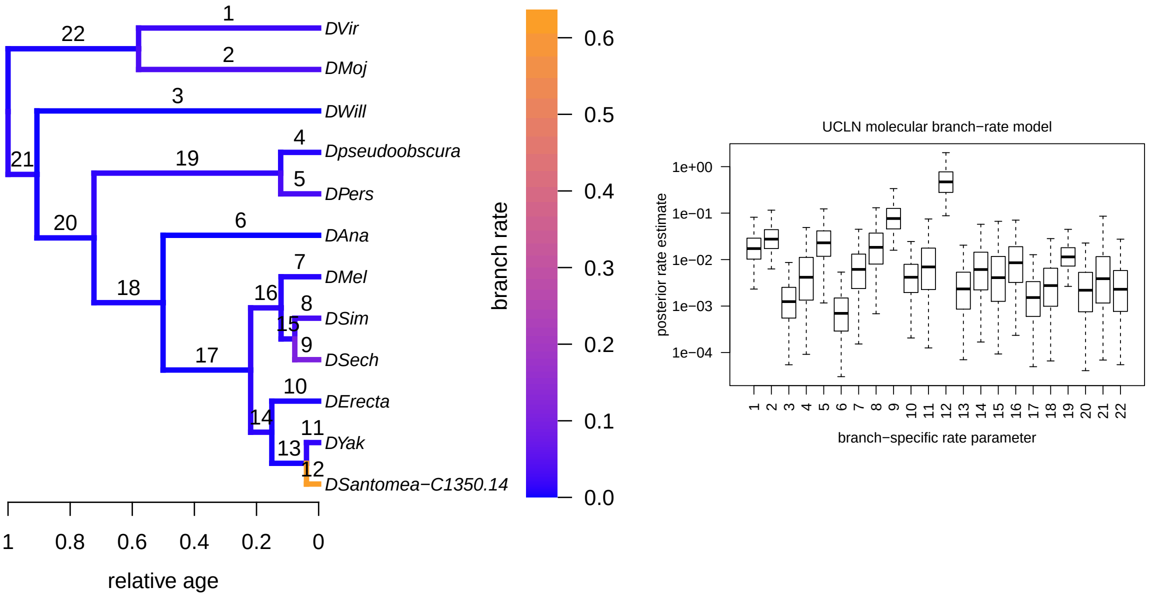


**Figure S14: Branch-specific rates of segment evolution under the UCLN molecular branch-rate prior model.** Left panel: branches are colored according to the posterior-mean rate of segment evolution on that branch. Labels above each branch indicate the corresponding branch index. Right panel: posterior distributions of branch-specific rate estimates (on a log scale). Boxes represent the 50% credible interval for each posterior distribution, and whiskers represent the 95% credible interval. Box indicies correspond to the branch labes indicated in the left panel.

File S2

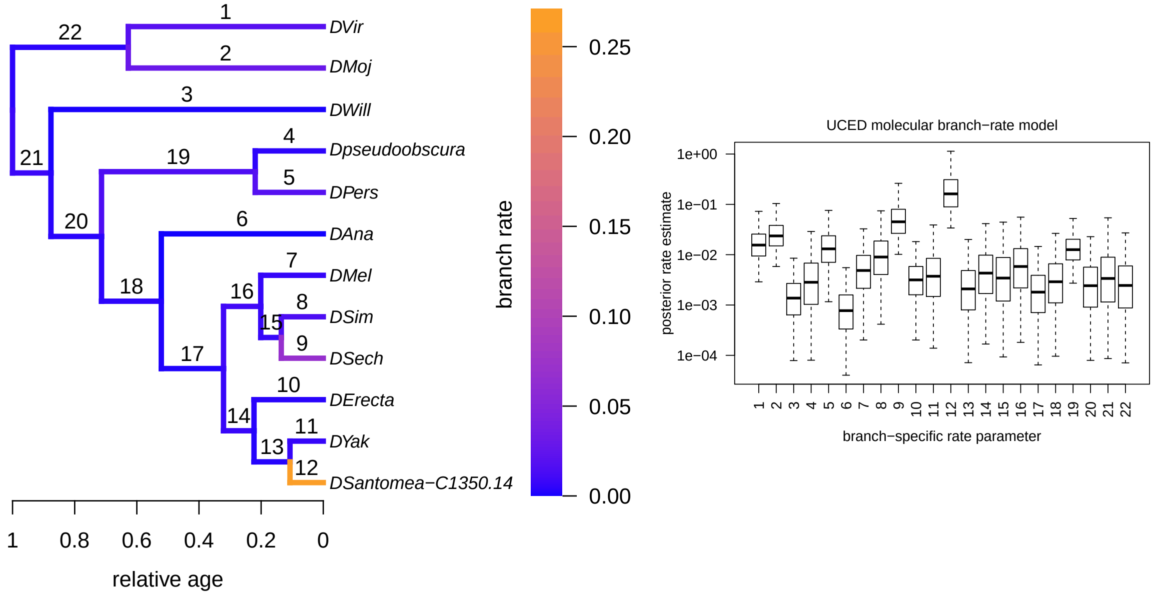


**Figure S15: Branch-specific rates of segment evolution under the UCED molecular branch-rate prior model.** Left panel: branches are colored according to the posterior-mean rate of segment evolution on that branch. Labels above each branch indicate the corresponding branch index. Right panel: posterior distributions of branch-specific rate estimates (on a log scale). Boxes represent the 50% credible interval for each posterior distribution, and whiskers represent the 95% credible interval. Box indicies correspond to the branch labes indicated in the left panel.

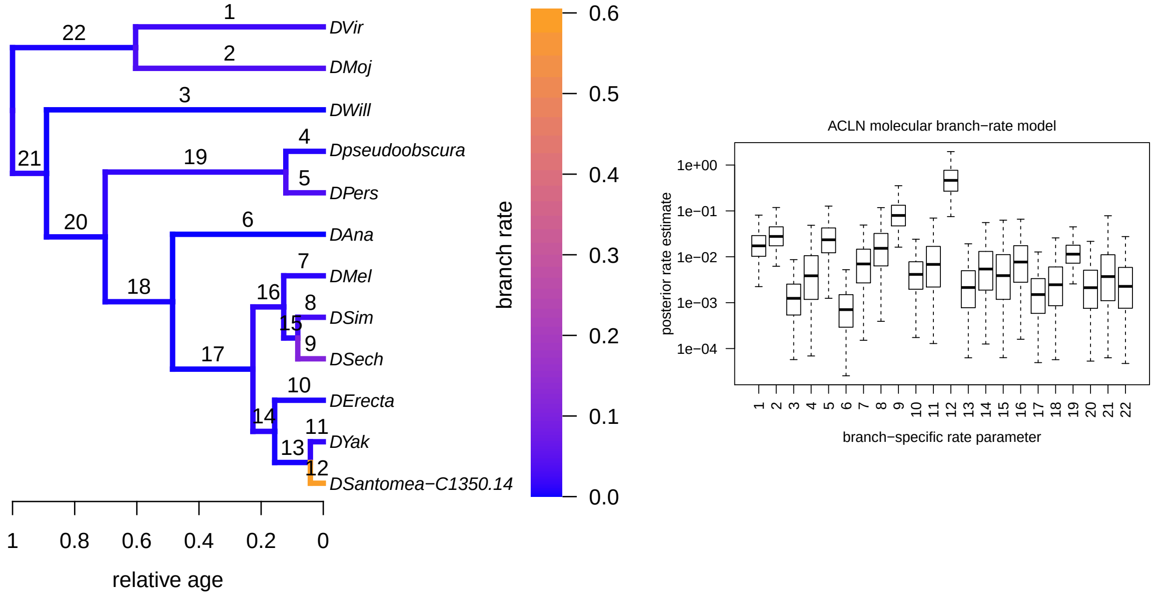


**Figure S16: Branch-specific rates of segment evolution under the ACLN molecular branch-rate prior model.** Left) Branches are colored according to the posterior mean rate of segment evolution on that branch. Labels above branches are the index of the branch. Right) Posterior distributions of branch-specific rate estimates (on a log scale). Boxes represent the 50% credible interval for each posterior distribution, and whiskers represent the 95% credible interval. Boxes are indexed according to the branch to which they correspond (as indicated in the left panel).

##### S3.4.5 Rates of change in relative segment size across branches

Under a constant-rate Brownian motion model (*i.e.*, where the rates of evolution on each branch are the same), changes in the continuous characters along branch $b$, $\delta_b$, are normally distributed with mean 0 and variance $t_b \Sigma$: because each branch may be a different length, the $\delta_b$ have different variances and thus are not drawn from the same normal distribution. Dividing the changes by the square-root of the branch length creates random variables that are independent and identically distributed:

$$\Delta_b = \delta_b / \sqrt{t_b}$$
$$\Delta_b \sim \text{MVN}(0, \Sigma).$$

In this section, we use posterior samples of the ancestral segment sizes to compute the *normalized* changes in the *relative segment sizes*. We then use these samples to summarize features of relative-segment-size evolution.

We used the posterior sample of ancestral log-segment sizes to summarize the rate of evolution of the *relative segment sizes* along each branch of the tree. For a given sample of ancestral states from the posterior distribution, we begin by computing the relative size of each segment, $i$, for each node, $n$, as follows:

$$r_n^i = \frac{\exp(x_n^i)}{\sum_j \exp(x_n^j)},$$

where we exponentiate each $x$ because we have samples of the log of the segment size, and the denominator is simply the sum of all segment sizes for node $n$. Next, we compute the amount of change in the $i^{\text{th}}$ segment on branch $b$ (we denote the node ancestral to branch $b$ as $b_a$, and the node descending from branch $b$ as $b_d$):

$$\delta_b^i = |r_d^i - r_a^i|.$$

We then compute the normalized rate of change in the relative size of segment $i$ by dividing the amount of change by the square-root of the length of branch $b$:

$$\Delta_b^i = \frac{\delta_b^i}{\sqrt{t_b}}.$$

We repeat this procedure for each segment and for each branch, and for each sample of ancestral states from the posterior distribution. This results in the posterior distribution of the normalized rate of change in each relative segment size along each branch in the phylogeny (Figures S17 to S19). We also summarized the posterior distribution of normalized changes in relative segment size across all segments and branches.

We used the sampled values of $\Delta_b^i$ to compute the correlation in rates of relative segment size evolution across the branches of the tree. In this case, we imagine that rate on each branch, $\Delta_b^i$, represents a sample from a multivariate distribution, and we compute the correlation matrix and partial correlation matrix between segments $i$ and $j$ using these samples (Figures S24, S26 to S28, S43 and S45). To understand how rates varied among segments, we computed the variance in the rate for each segment, $\text{Var}(\Delta_b^i)$ (Figures S29 to S31).

We also computed the normalized rate of change across all segments by computing the total amount of change on branch $b$:

$$\delta_b = \sqrt{\sum_{i=1}^{8} \left( r_d^i - r_a^i \right)^2}.$$

The normalized rate of change across all segments on branch $b$ is then:

$$\Delta_b = \frac{\delta_b}{\sqrt{t_b}}.$$

This results in the posterior distribution of the rate of change in relative segment size (across segments) for each branch in the phylogeny (Figures S32 to S34).

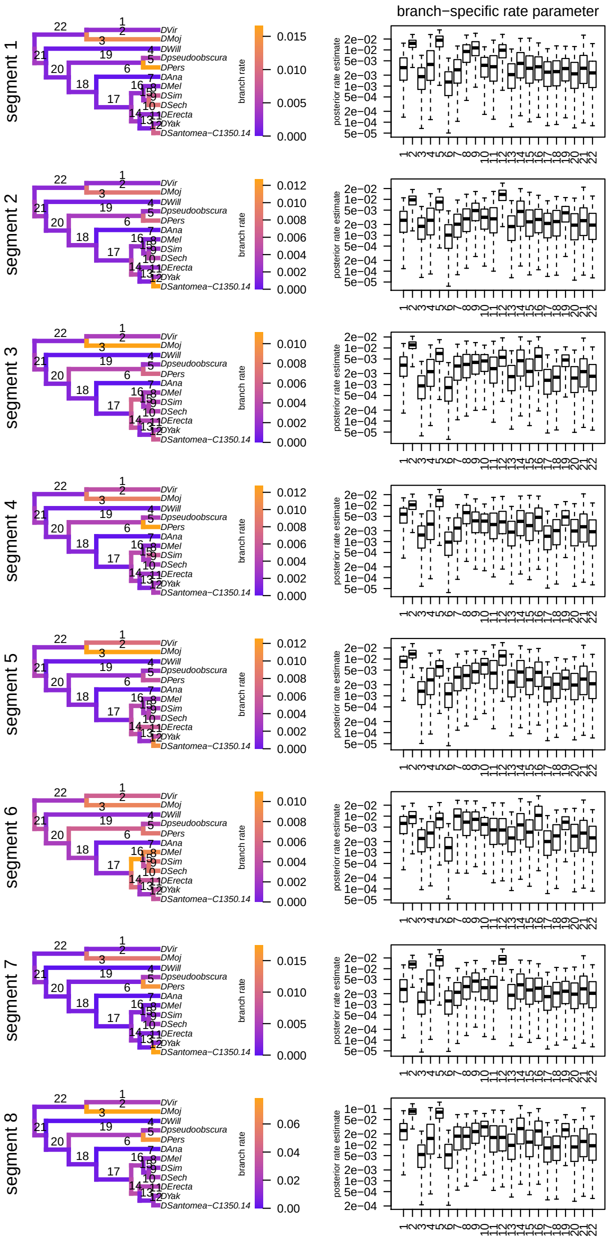


**Figure S17: Normalized rates of change in relative segment size for each segment under the UCLN molecular branch-rate model.** Left panels) Branches are colored according to the posterior mean rate of relative segment size evolution on that branch. Labels above branches are the index of the branch. Right panels) Posterior distributions of branch-specific rate estimates (on a log scale). Boxes represent the 50% credible interval for each posterior distribution, and whiskers represent the 95% credible interval. Boxes are indexed according to the branch to which they correspond (as indicated in the left panel).

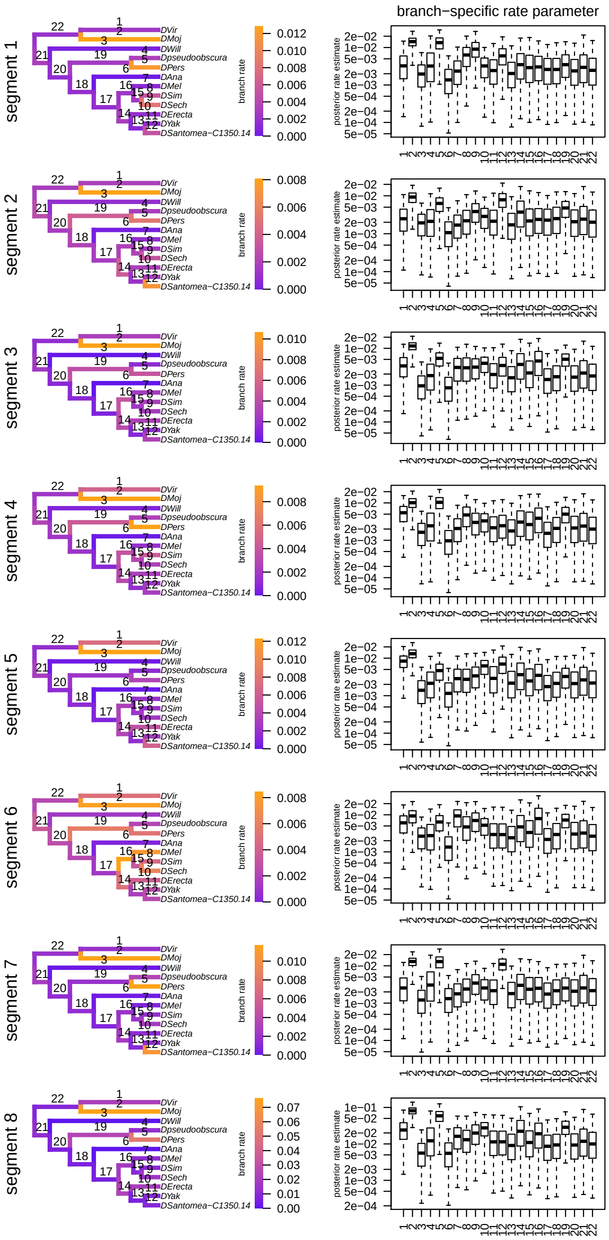


**Figure S18: Estimated rates of change in relative segment size for each segment under the UCED molecular branch-rate model.** Left panels) Branches are colored according to the posterior mean rate of relative segment size evolution on that branch. Labels above branches are the index of the branch. Right panels) Posterior distributions of branch-specific rate estimates (on a log scale). Boxes represent the 50% credible interval for each posterior distribution, and whiskers represent the 95% credible interval. Boxes are indexed according to the branch to which they correspond (as indicated in the left panel).

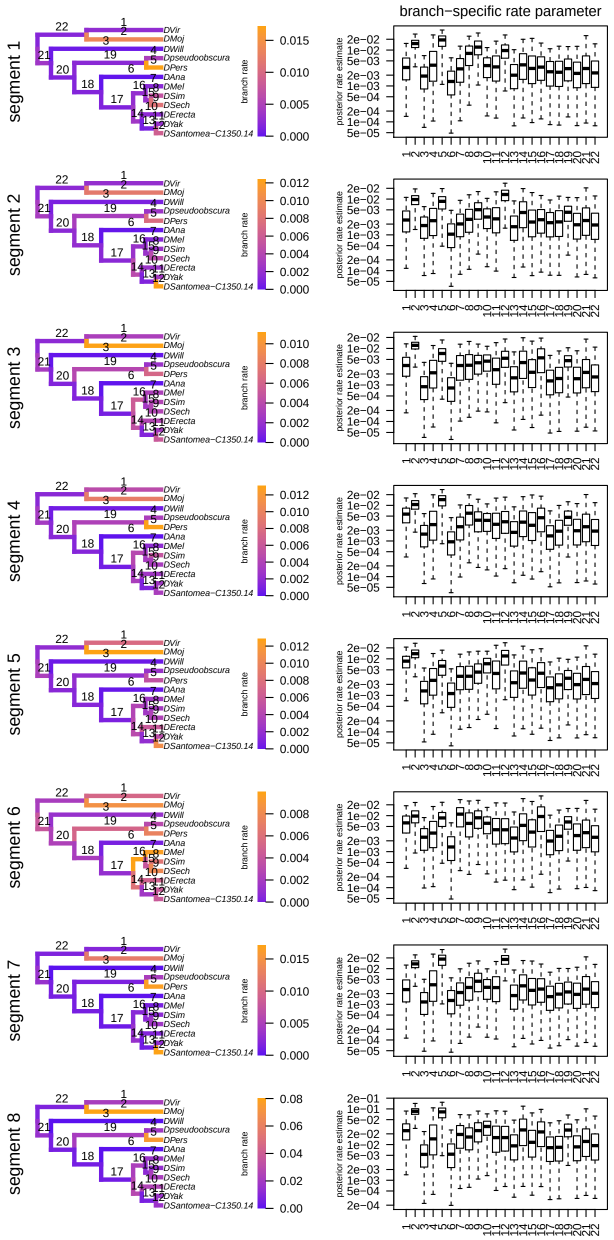


**Figure S19: Estimated rates of change in relative-segment size for each segment under the ACLN molecular branch-rate model.** Left panels: branches are colored according to the posterior-mean rate of relative-segment size evolution on that branch. Labels above each branch indicate the corresponding branch index. Right panels. posterior distributions of branch-specific rate estimates (on a log scale). Boxes represent the 50% credible interval for each posterior distribution, and whiskers represent the 95% credible interval. Box indicies correspond to the branch labes indicated in the left panel.

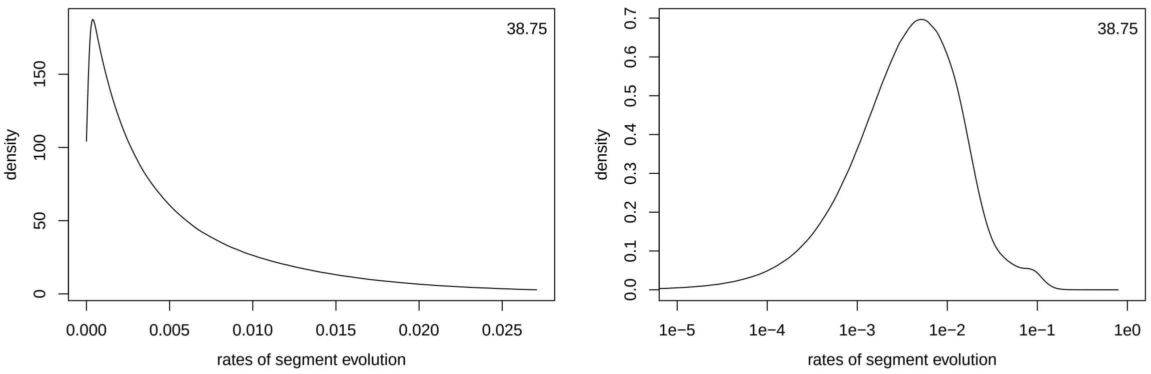


**Figure S20: Normalized changes in relative segment size across all segments and branches under the UCLN molecular branch-rate model.** Left) The posterior distribution of normalized changes in relative segment sizes across branches and across all samples of the posterior distribution. Right) The posterior distribution of normalized changes in relative segment sizes across branches and across all samples of the posterior distribution (on the log scale). Numbers in the upper right are the excess kurtosis.

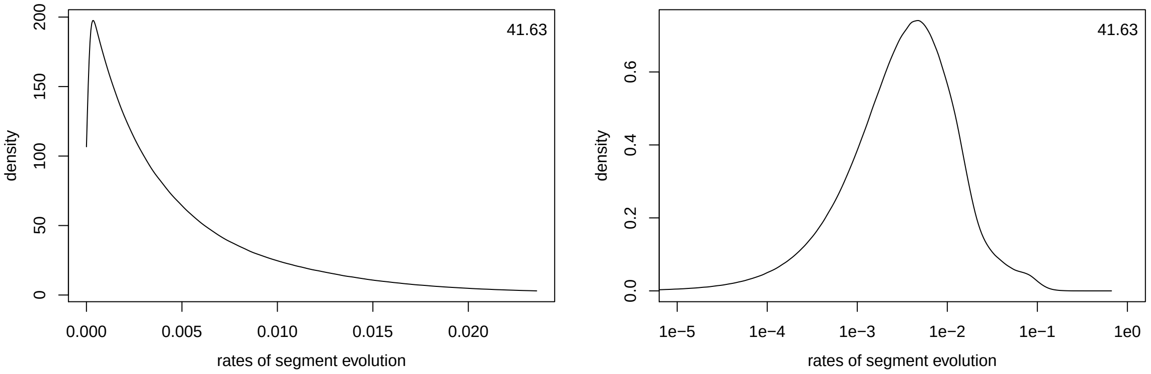


**Figure S21: Normalized changes in relative segment size across all segments and branches under the UCED molecular branch-rate model.** Left) The posterior distribution of normalized changes in relative segment sizes across branches and across all samples of the posterior distribution. Right) The posterior distribution of normalized changes in relative segment sizes across branches and across all samples of the posterior distribution (on the log scale). Numbers in the upper right are the excess kurtosis.

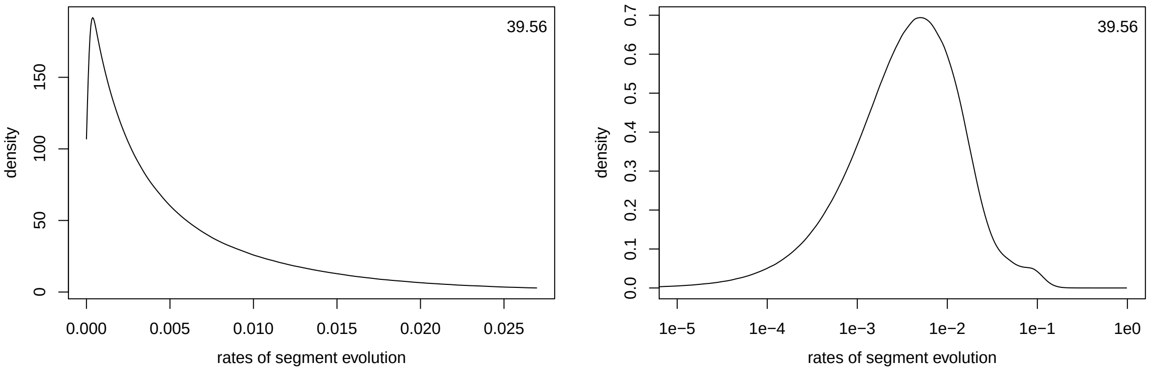


**Figure S22: Normalized changes in relative segment size across all segments and branches under the ACLN molecular branch-rate model.** Left) The posterior distribution of normalized changes in relative segment sizes across branches and across all samples of the posterior distribution. Right) The posterior distribution of normalized changes in relative segment sizes across branches and across all samples of the posterior distribution (on the log scale). Numbers in the upper right are the excess kurtosis.

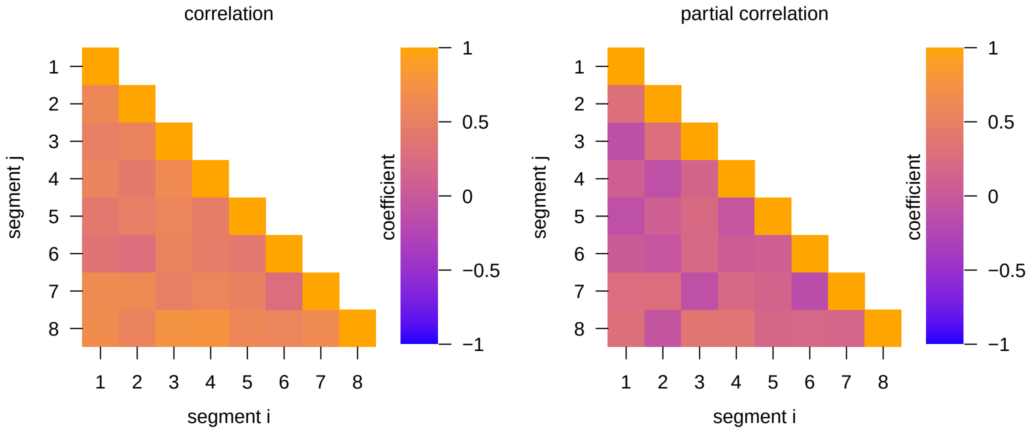


**Figure S23: Correlation coefficients between rates of relative-segment-size evolution for each pair of segments under the UCLN molecular branch-rate model.** Left) Correlation coefficients between rates of evolution for each pair of relative segment sizes. Right) Partial correlation coefficients between rates of evolution for each pair of relative segment sizes.

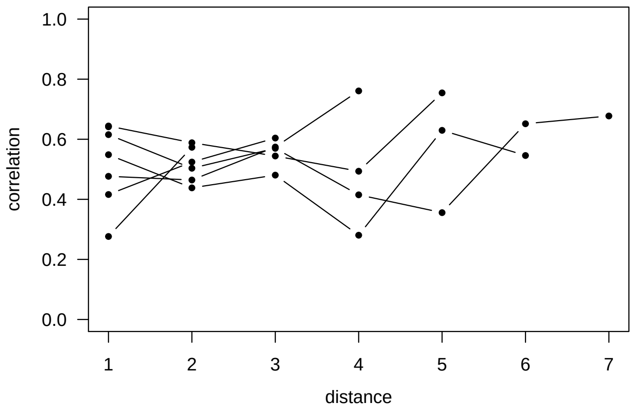


**Figure S24: Correlation coefficients between rates of evolution for adjacent pairs of relative segment sizes under the UCLN molecular branch-rate model.** Posterior mean correlation coefficients (blue) and partial correlation coefficients (orange) between rates of evolution for adjacent pairs of relative segment sizes. Whiskers represent the 95% posterior credible interval.

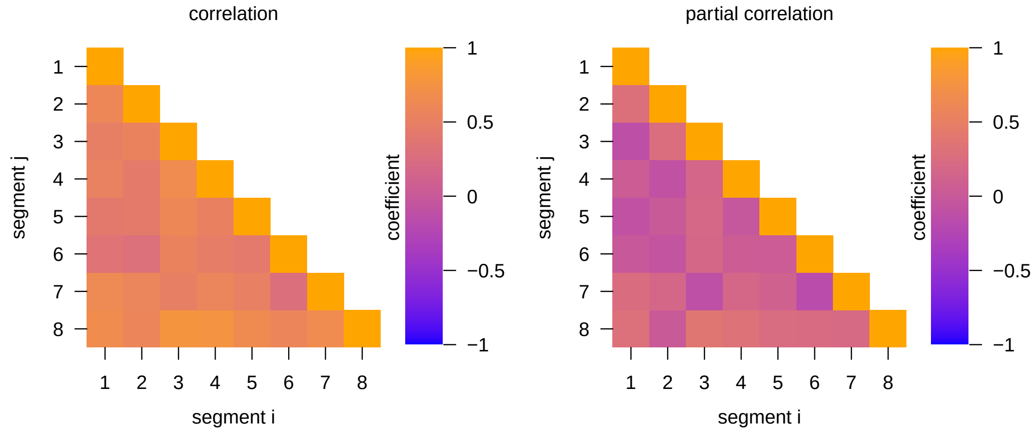


**Figure S25: Correlation coefficients between rates of relative-segment-size evolution for each pair of segments under the UCED molecular branch-rate model.** Left) Correlation coefficients between rates of evolution for each pair of relative segment sizes. Right) Partial correlation coefficients between rates of evolution for each pair of relative segment sizes.

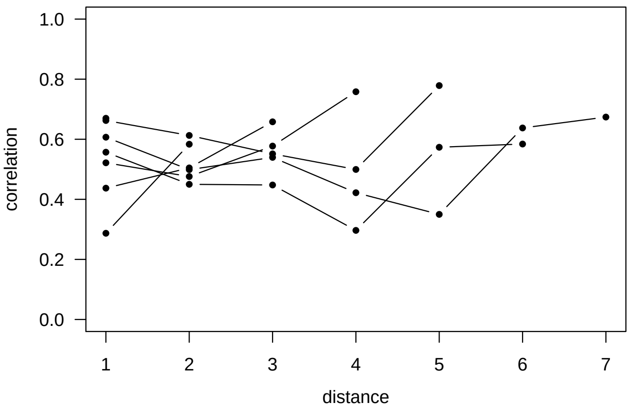


**Figure S26: Correlation coefficients between rates of evolution for adjacent pairs of relative segment sizes under the UCED molecular branch-rate model.** Posterior mean correlation coefficients (blue) and partial correlation coefficients (orange) between rates of evolution for adjacent pairs of relative segment sizes. Whiskers represent the 95% posterior credible interval.

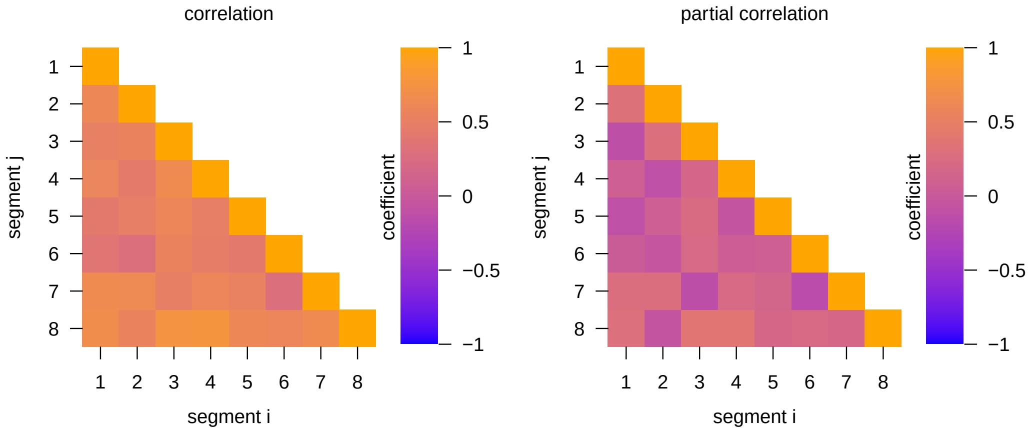


**Figure S27: Correlation coefficients between rates of relative-segment-size evolution for each pair of segments under the ACLN molecular branch-rate model.** Left) Correlation coefficients between rates of evolution for each pair of relative segment sizes. Right) Partial correlation coefficients between rates of evolution for each pair of relative segment sizes.

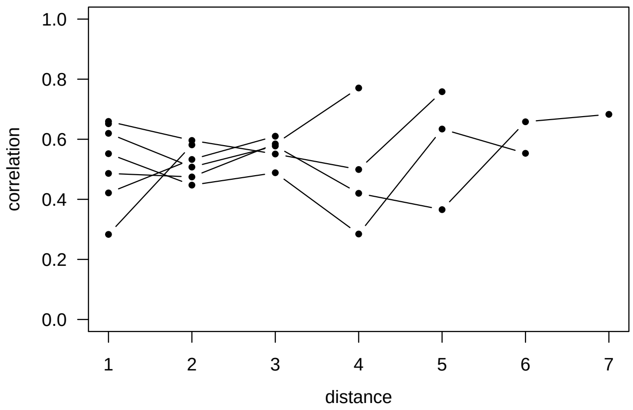


**Figure S28: Correlation coefficients between rates of evolution for adjacent pairs of relative segment sizes under the ACLN molecular branch-rate model.** Posterior mean correlation coefficients (blue) and partial correlation coefficients (orange) between rates of evolution for adjacent pairs of relative segment sizes. Whiskers represent the 95% posterior credible interval.

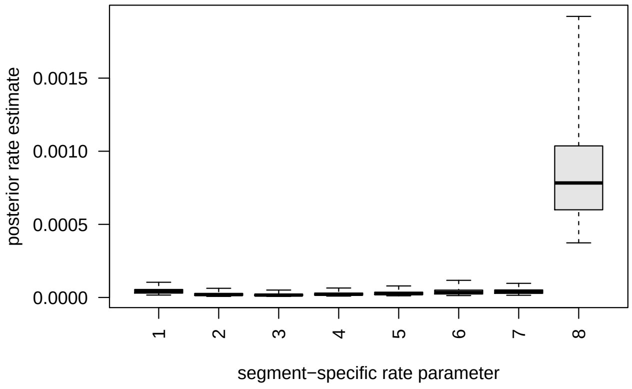


**Figure S29: Rates of size evolution among segments under the UCLN molecular branch-rate model.** Boxes represent the 50% credible interval for each posterior distribution, and whiskers represent the 95% credible interval.

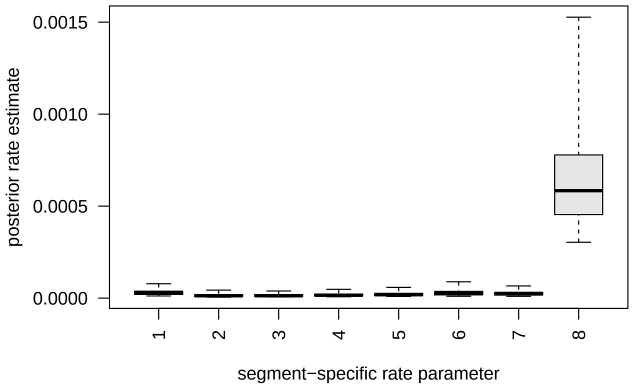


**Figure S30: Rates of size evolution among segments under the UCED molecular branch-rate model.** Boxes represent the 50% credible interval for each posterior distribution, and whiskers represent the 95% credible interval.

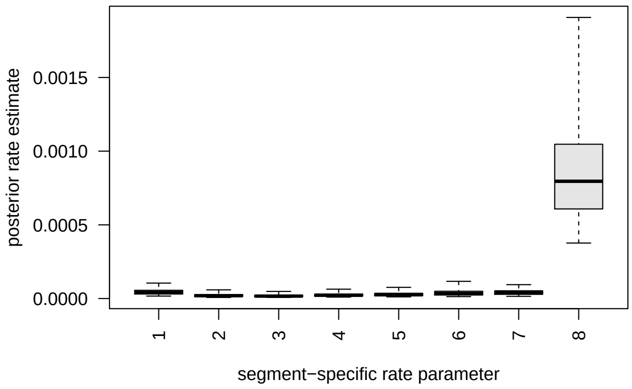


**Figure S31: Rates of size evolution among segments under the ACLN molecular branch-rate model.** Boxes represent the 50% credible interval for each posterior distribution, and whiskers represent the 95% credible interval.

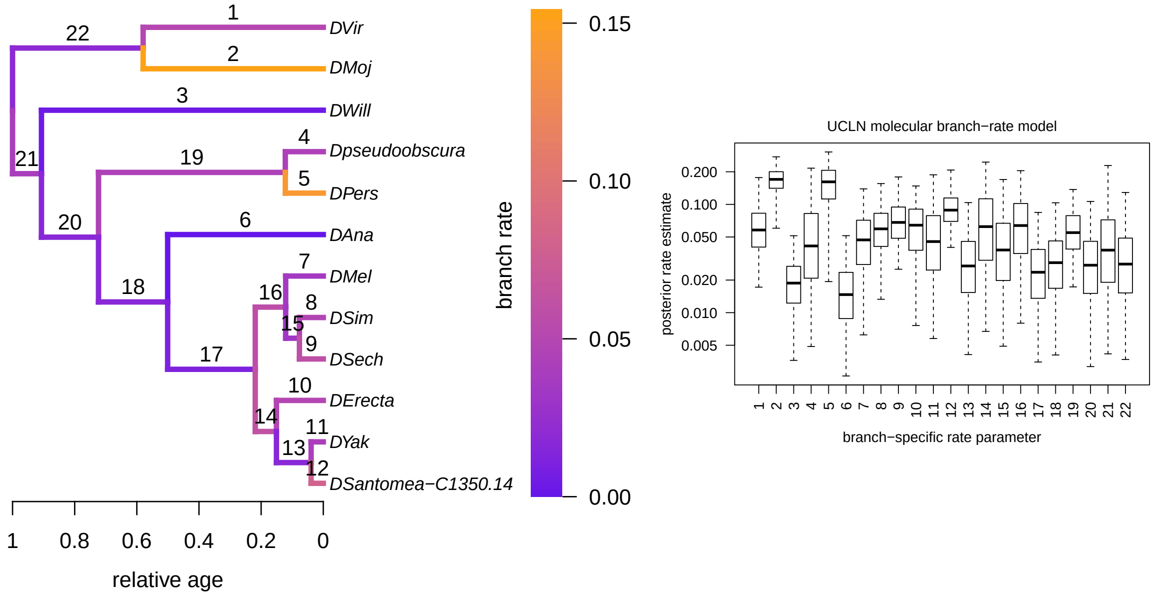


**Figure S32: Branch-specific rates of relative segment-size evolution under the UCLN molecular branch-rate prior model.** Left) Branches are colored according to the posterior mean rate of relative segment-size evolution, $\rho_b$, on that branch. Labels above branches are the index of the branch. Right) Posterior distributions of branch-specific rates of relative semgnet-size evolution, $\rho_b$ (on a log scale). Boxes represent the 50% credible interval for each posterior distribution, and whiskers represent the 95% credible interval. Boxes are indexed according to the branch to which they correspond (as indicated in the left panel).

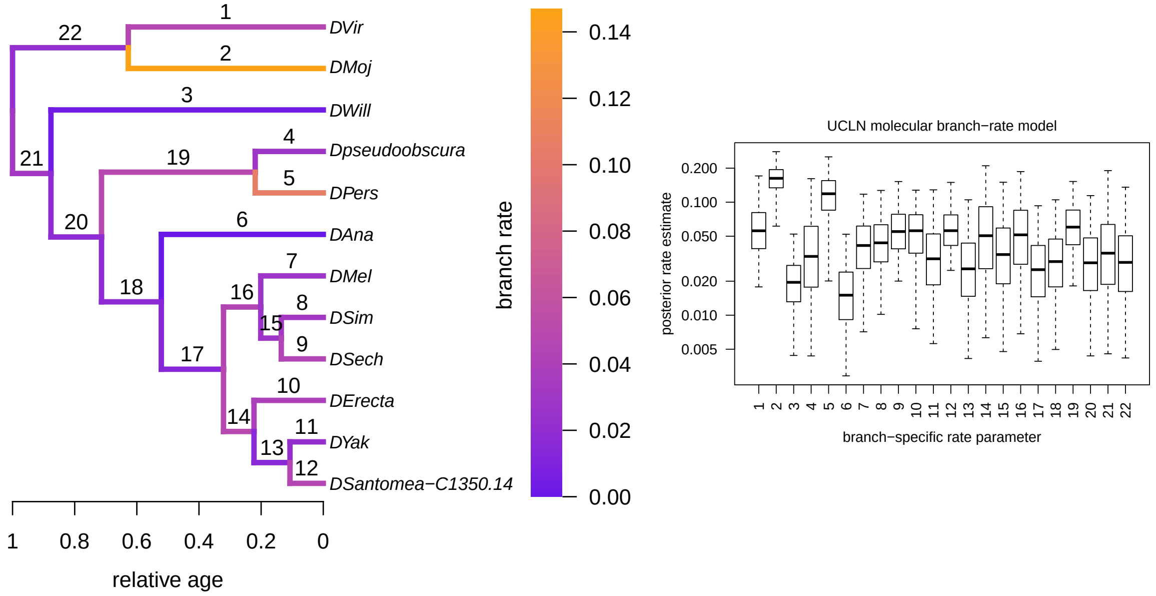


**Figure S33: Branch-specific rates of relative segment-size evolution under the UCED molecular branch-rate prior model.** Left) Branches are colored according to the posterior mean rate of relative segment-size evolution, $\rho_b$, on that branch. Labels above branches are the index of the branch. Right) Posterior distributions of branch-specific rates of relative semgnet-size evolution, $\rho_b$ (on a log scale). Boxes represent the 50% credible interval for each posterior distribution, and whiskers represent the 95% credible interval. Boxes are indexed according to the branch to which they correspond (as indicated in the left panel).

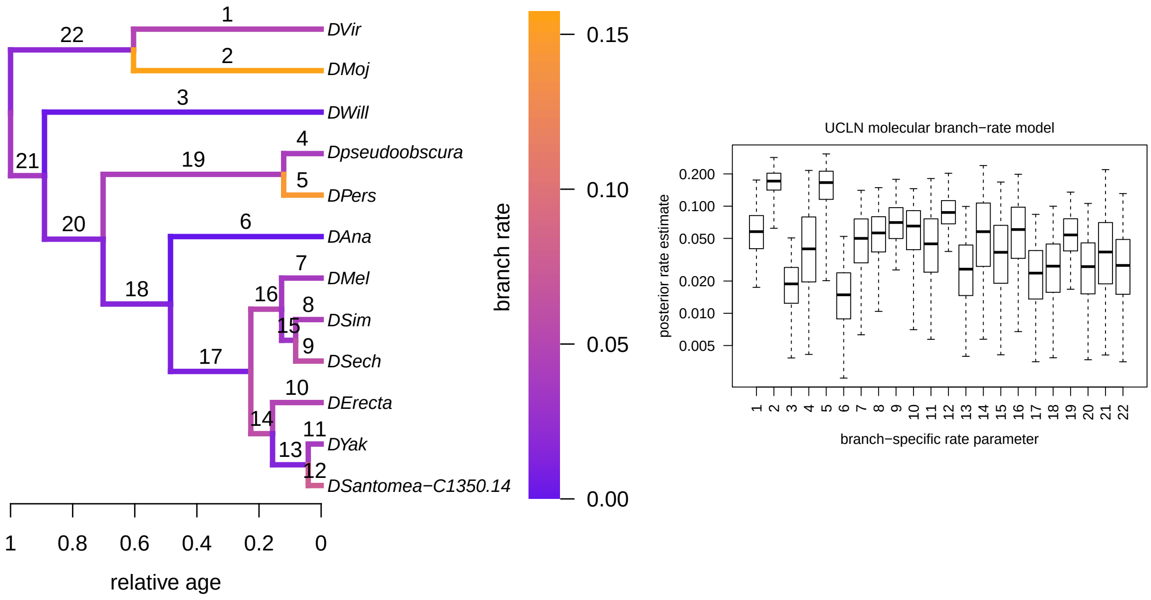


**Figure S34: Branch-specific rates of relative segment-size evolution under the ACLN molecular branch-rate prior model.** Left) Branches are colored according to the posterior mean rate of relative segment-size evolution, $\rho_b$, on that branch. Labels above branches are the index of the branch. Right) Posterior distributions of branch-specific rates of relative semgnet-size evolution, $\rho_b$ (on a log scale). Boxes represent the 50% credible interval for each posterior distribution, and whiskers represent the 95% credible interval. Boxes are indexed according to the branch to which they correspond (as indicated in the left panel).

##### S3.4.6 Rates of change in relative segment position across branches

We also summarized the rate to summarize the rate at which *relative-segment position* evolves along each branch of the tree. For a given sample of ancestral states from the posterior distribution, we begin by computing the size of the head and thorax for each node, $n$, as follows:

$$h_n = \exp(x_n^9) - \sum_{i=1}^{8} \exp(x_n^i),$$

where $x_n^9$ is the ancestral-state estimate for the *total-body length*, and the sum is over the eight remaining segment sizes. We then compute the position of the anterior boundary of each segment of the eight segments, $i = \{1, 2, \ldots, 8\}$, for each node, $n$, as follows:

$$p_n^i = \begin{cases} h_n / \exp(x_n^9) & \text{if } i = 1 \\ p_n^{i-1} + \exp(x_n^i) / \exp(x_n^9) & \text{otherwise,} \end{cases}$$

*i.e.*, the anterior boundary of the first segment is simply the length of the head and thorax (divided by the total-body length), while the anterior position for each other segment is the anterior position for the previous segment plus the size of the focal segment (divided by the total body length).

We repeated each summary described in the previous section on the relative segment positions.

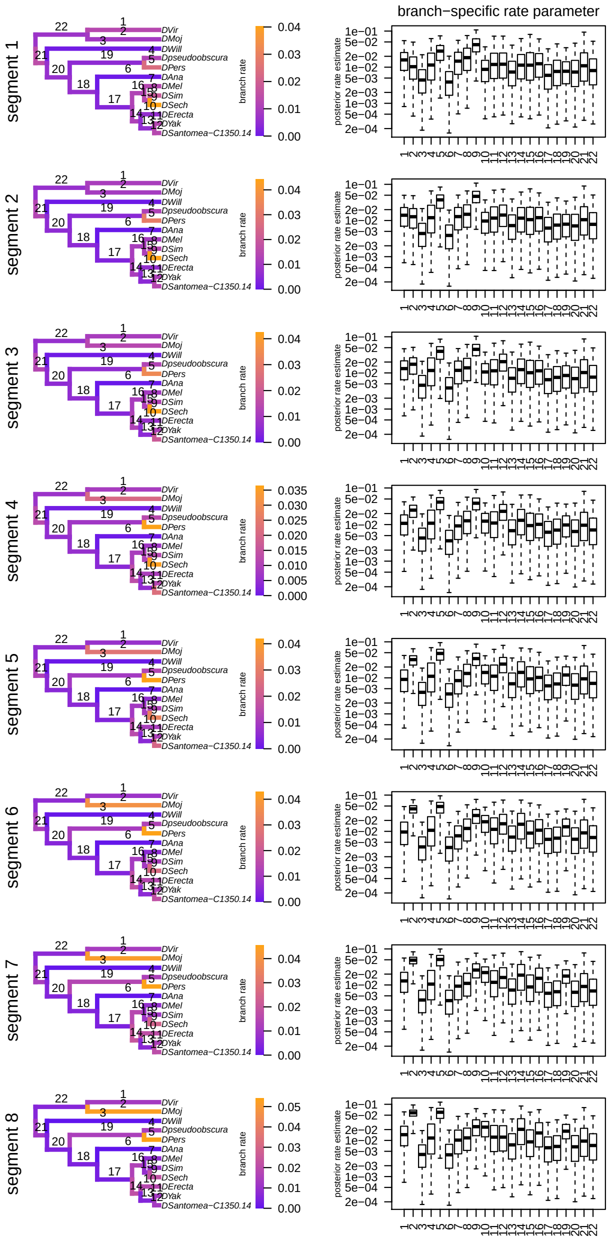


**Figure S35: Normalized rates of change in relative segment position for each segment under the UCLN molecular branch-rate model.** Left panels) Branches are colored according to the posterior mean rate of relative segment position evolution on that branch. Labels above branches are the index of the branch. Right panels) Posterior distributions of branch-specific rate estimates (on a log scale). Boxes represent the 50% credible interval for each posterior distribution, and whiskers represent the 95% credible interval. Boxes are indexed according to the branch to which they correspond (as indicated in the left panel).

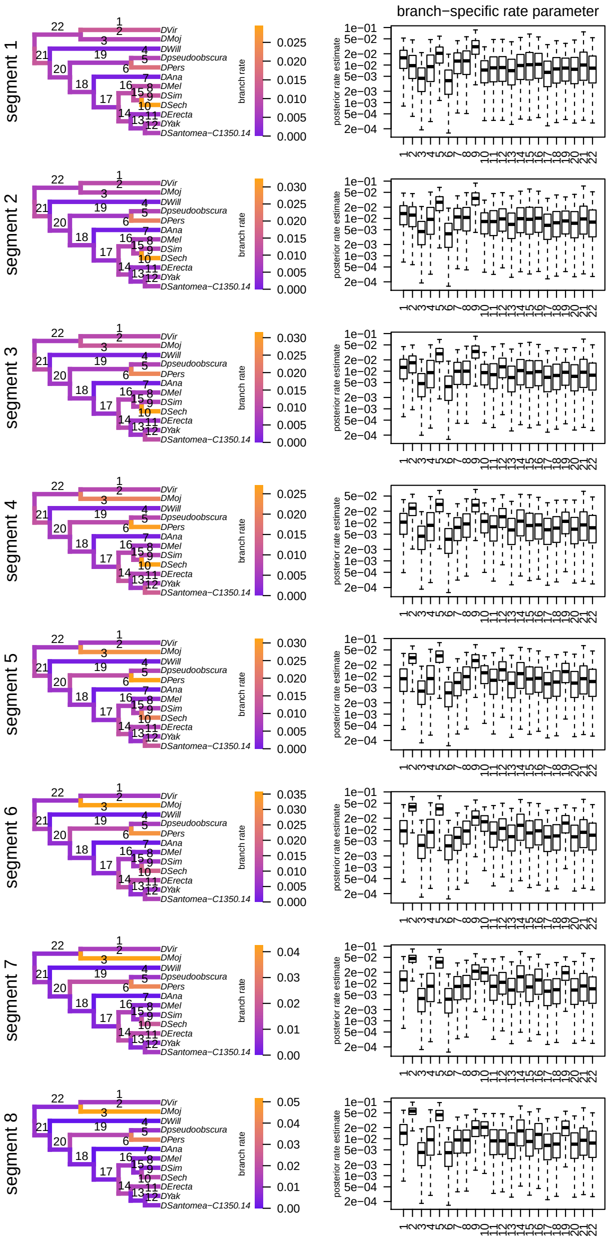


**Figure S36: Normalized rates of change in relative segment position for each segment under the UCED molecular branch-rate model.** Left panels) Branches are colored according to the posterior mean rate of relative segment position evolution on that branch. Labels above branches are the index of the branch. Right panels) Posterior distributions of branch-specific rate estimates (on a log scale). Boxes represent the 50% credible interval for each posterior distribution, and whiskers represent the 95% credible interval. Boxes are indexed according to the branch to which they correspond (as indicated in the left panel).

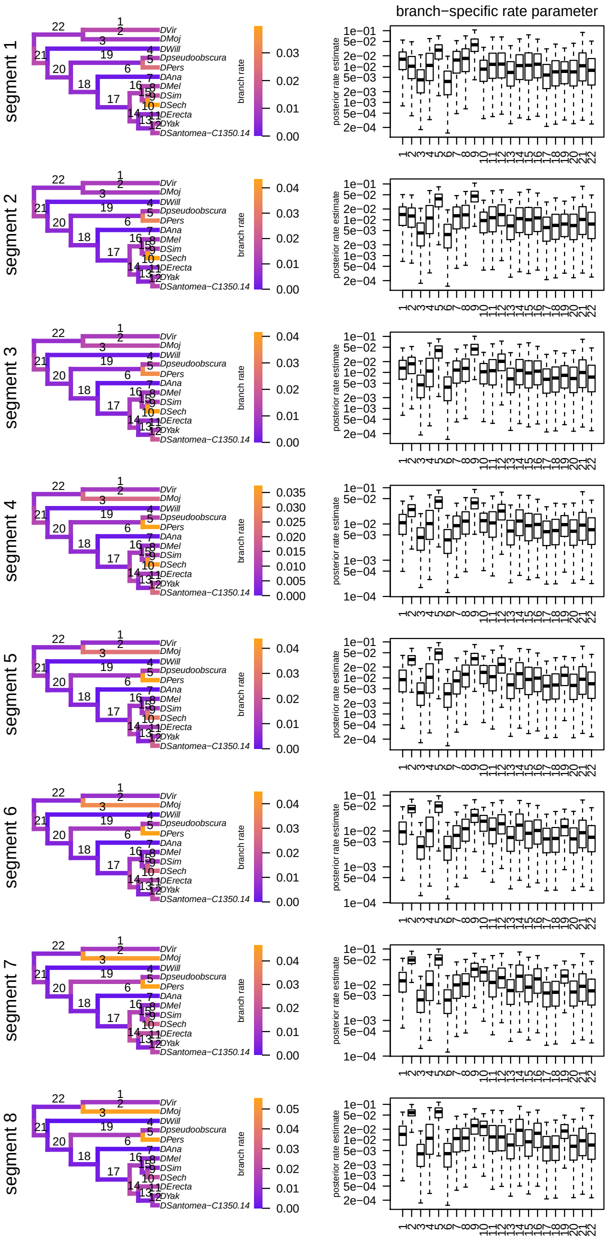


**Figure S37: Normalized rates of change in relative segment position for each segment under the ACLN molecular branch-rate model.** Left panels) Branches are colored according to the posterior mean rate of relative segment position evolution on that branch. Labels above branches are the index of the branch. Right panels) Posterior distributions of branch-specific rate estimates (on a log scale). Boxes represent the 50% credible interval for each posterior distribution, and whiskers represent the 95% credible interval. Boxes are indexed according to the branch to which they correspond (as indicated in the left panel).

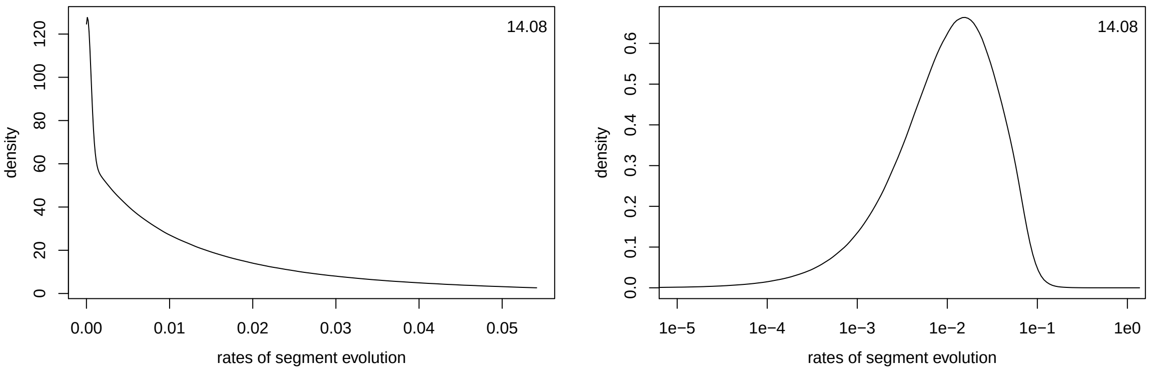


**Figure S38: Normalized changes in relative segment position across all segments and branches under the UCLN molecular branch-rate model.** Left) The posterior distribution of normalized changes in relative segment positions across branches and across all samples of the posterior distribution. Right) The posterior distribution of normalized changes in relative segment positions across branches and across all samples of the posterior distribution (on the log scale). Numbers in the upper right are the excess kurtosis.

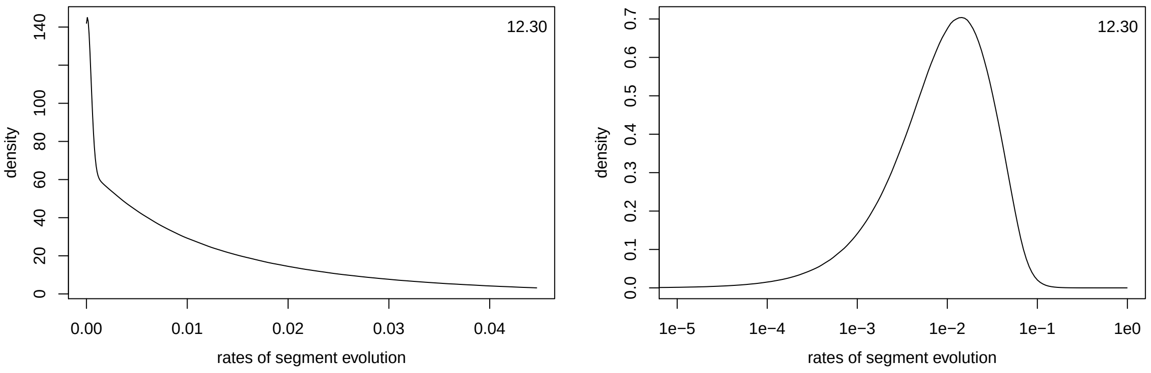


**Figure S39: Normalized changes in relative segment position across all segments and branches under the UCED molecular branch-rate model.** Left) The posterior distribution of normalized changes in relative segment positions across branches and across all samples of the posterior distribution. Right) The posterior distribution of normalized changes in relative segment positions across branches and across all samples of the posterior distribution (on the log scale). Numbers in the upper right are the excess kurtosis.

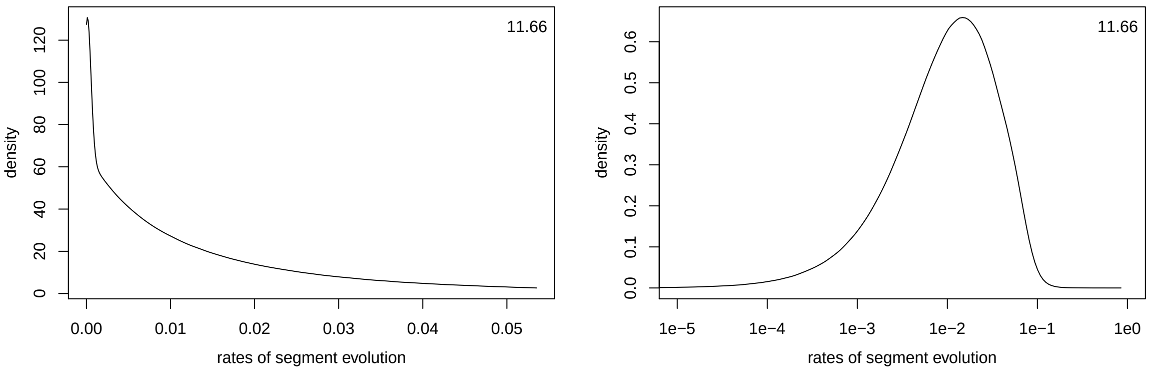


**Figure S40: Normalized changes in relative segment position across all segments and branches under the ACLN molecular branch-rate model.** Left) The posterior distribution of normalized changes in relative segment positions across branches and across all samples of the posterior distribution. Right) The posterior distribution of normalized changes in relative segment positions across branches and across all samples of the posterior distribution (on the log scale). Numbers in the upper right are the excess kurtosis.

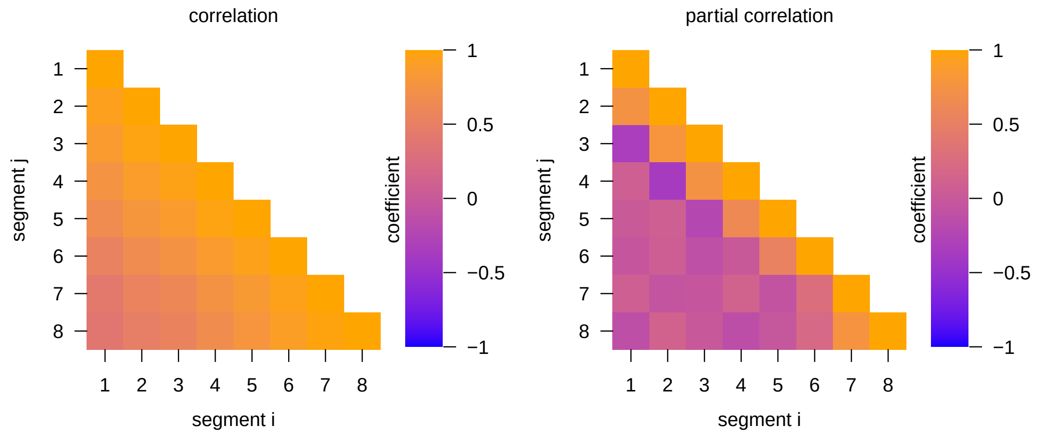


**Figure S41: Correlation coefficients between rates of relative-segment-position evolution for each pair of segments under the UCLN molecular branch-rate model.** Left) Correlation coefficients between rates of evolution for each pair of relative segment positions. Right) Partial correlation coefficients between rates of evolution for each pair of relative segment positions.

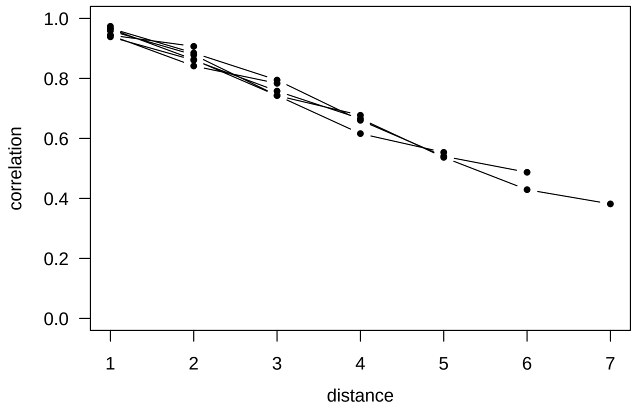


**Figure S42: Correlation coefficients between rates of evolution for adjacent pairs of relative segment positions under the UCLN molecular branch-rate model.** Posterior mean correlation coefficients (blue) and partial correlation coefficients (orange) between rates of evolution for adjacent pairs of relative segment positions. Whiskers represent the 95% posterior credible interval.

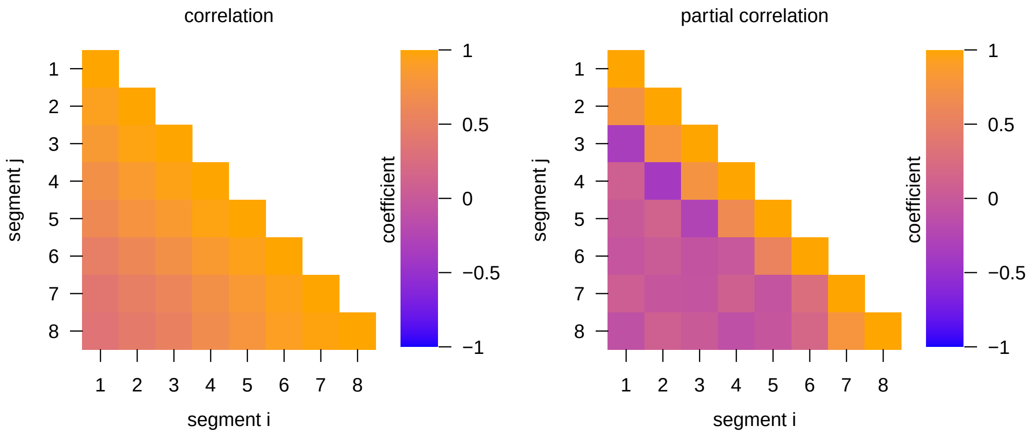


**Figure S43: Correlation coefficients between rates of relative-segment-position evolution for each pair of segments under the UCED molecular branch-rate model.** Left) Correlation coefficients between rates of evolution for each pair of relative segment positions. Right) Partial correlation coefficients between rates of evolution for each pair of relative segment positions.

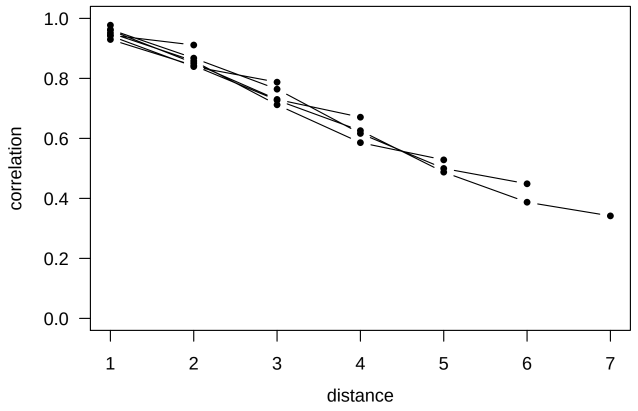


**Figure S44: Correlation coefficients between rates of evolution for adjacent pairs of relative segment positions under the UCED molecular branch-rate model.** Posterior mean correlation coefficients (blue) and partial correlation coefficients (orange) between rates of evolution for adjacent pairs of relative segment positions. Whiskers represent the 95% posterior credible interval.

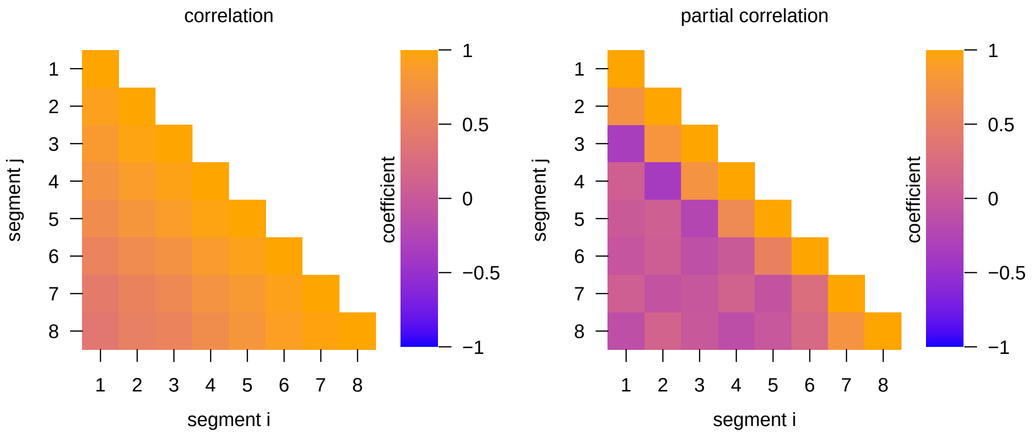


**Figure S45: Correlation coefficients between rates of relative-segment-position evolution for each pair of segments under the ACLN molecular branch-rate model.** Left) Correlation coefficients between rates of evolution for each pair of relative segment positions. Right) Partial correlation coefficients between rates of evolution for each pair of relative segment positions.

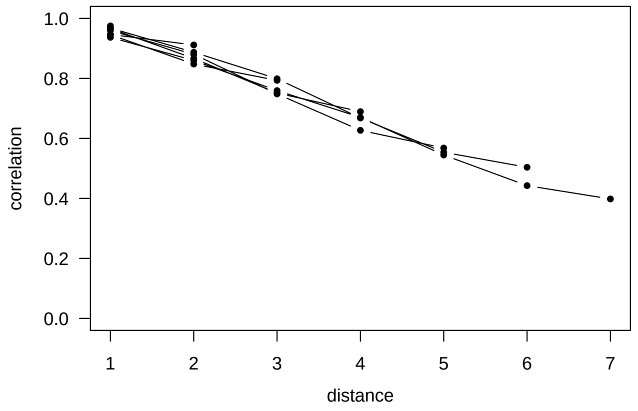


**Figure S46: Correlation coefficients between rates of evolution for adjacent pairs of relative segment positions under the ACLN molecular branch-rate model.** Posterior mean correlation coefficients (blue) and partial correlation coefficients (orange) between rates of evolution for adjacent pairs of relative segment positions. Whiskers represent the 95% posterior credible interval.

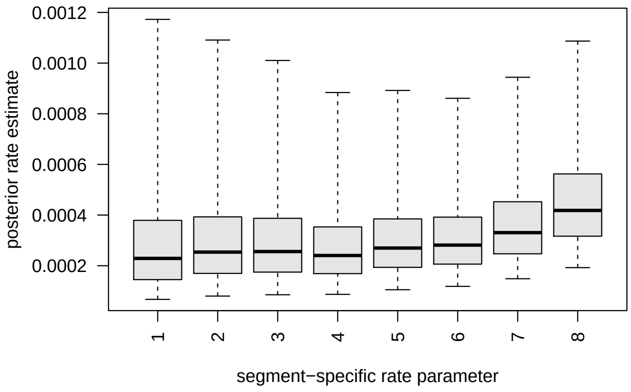


**Figure S47: Rates of position evolution among segments under the UCLN molecular branch-rate model.** Boxes represent the 50% credible interval for each posterior distribution, and whiskers represent the 95% credible interval.

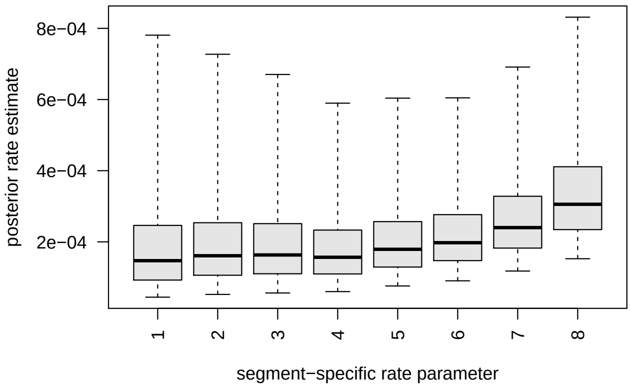


**Figure S48: Rates of position evolution among segments under the UCED molecular branch-rate model.** Boxes represent the 50% credible interval for each posterior distribution, and whiskers represent the 95% credible interval.

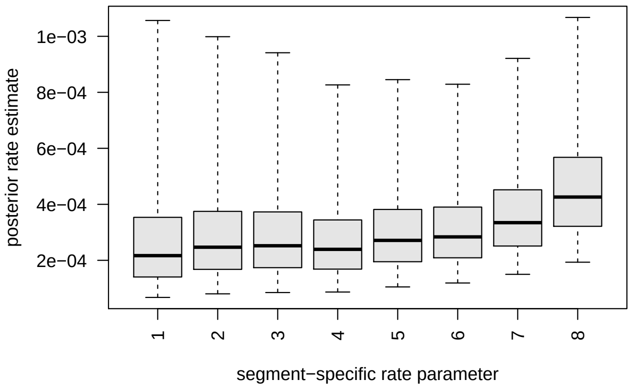


**Figure S49: Rates of position evolution among segments under the ACLN molecular branch-rate model.** Boxes represent the 50% credible interval for each posterior distribution, and whiskers represent the 95% credible interval.

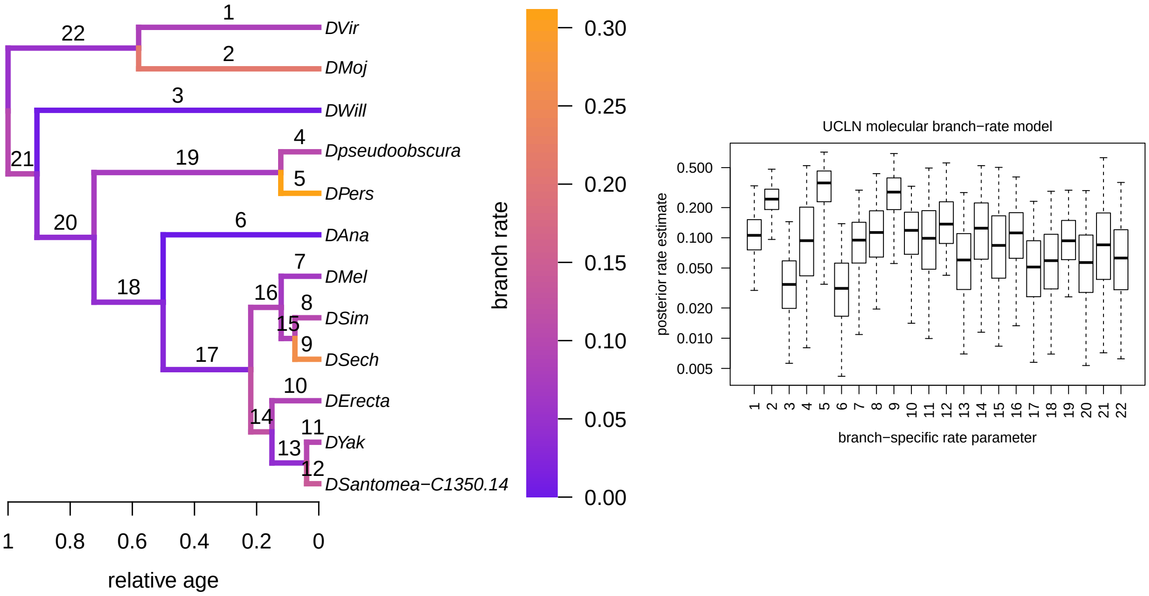


**Figure S50: Branch-specific rates of relative segment-size evolution under the UCLN molecular branch-rate prior model.** Left) Branches are colored according to the posterior mean rate of relative segment-position evolution, $\rho_b$, on that branch. Labels above branches are the index of the branch. Right) Posterior distributions of branch-specific rates of relative semgnet-position evolution, $\rho_b$ (on a log scale). Boxes represent the 50% credible interval for each posterior distribution, and whiskers represent the 95% credible interval. Boxes are indexed according to the branch to which they correspond (as indicated in the left panel).

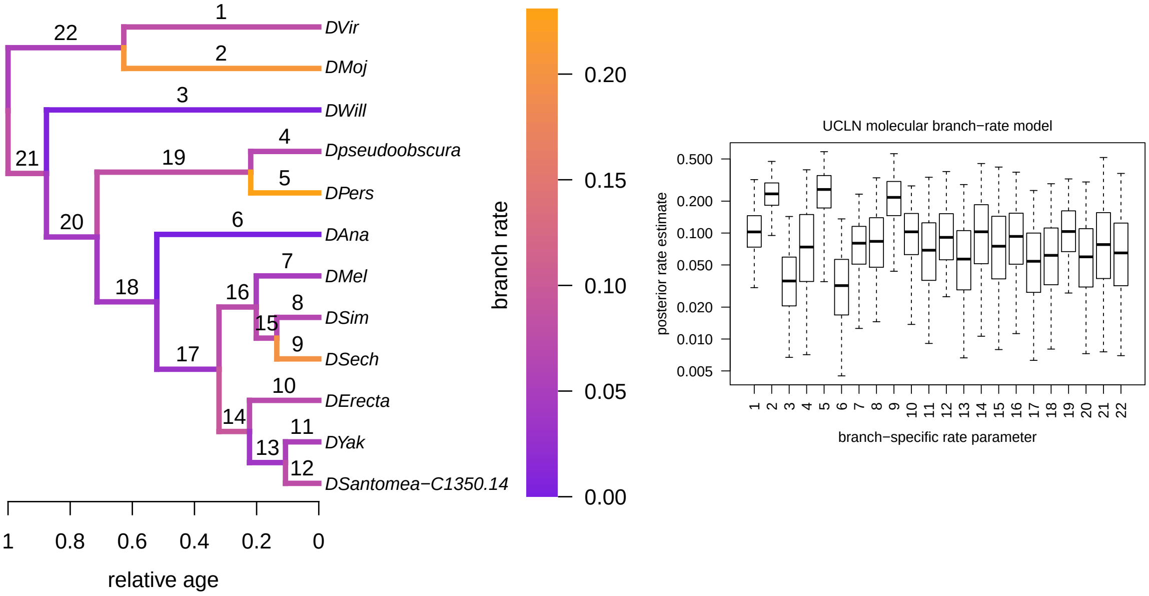


**Figure S51: Branch-specific rates of relative segment-size evolution under the UCED molecular branch-rate prior model.** Left) Branches are colored according to the posterior mean rate of relative segment-position evolution, $\rho_b$, on that branch. Labels above branches are the index of the branch. Right) Posterior distributions of branch-specific rates of relative semgnet-position evolution, $\rho_b$ (on a log scale). Boxes represent the 50% credible interval for each posterior distribution, and whiskers represent the 95% credible interval. Boxes are indexed according to the branch to which they correspond (as indicated in the left panel).

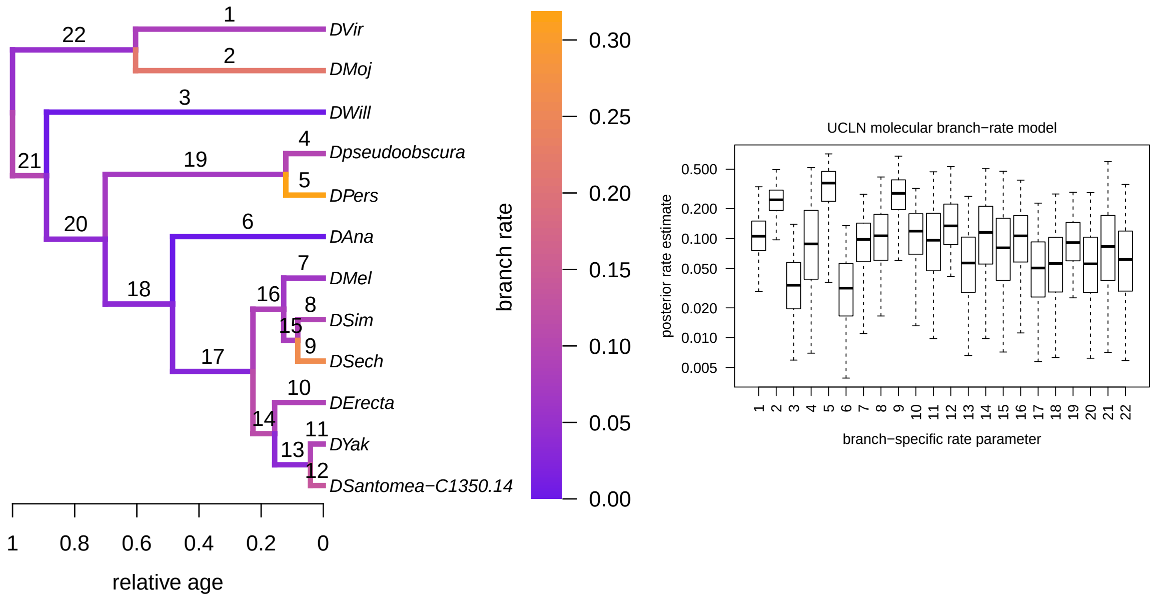


**Figure S52: Branch-specific rates of relative segment-size evolution under the ACLN molecular branch-rate prior model.** Left) Branches are colored according to the posterior mean rate of relative segment-position evolution, $\rho_b$, on that branch. Labels above branches are the index of the branch. Right) Posterior distributions of branch-specific rates of relative semgnet-position evolution, $\rho_b$ (on a log scale). Boxes represent the 50% credible interval for each posterior distribution, and whiskers represent the 95% credible interval. Boxes are indexed according to the branch to which they correspond (as indicated in the left panel).
