## Supplementary material for "Evolution of Larval Segment Position across 12 *Drosophila* Species": FigS: File_S3.docx

**Supplementary File 3**

**Method for removing anterior or posterior end from segment position calculations**

*Calculating relative segment position in the absence of posterior-most segments*

In order to calculate relative segment position in the absence of segment A8 and body parts posterior to it (A8 to the tail of the larva), we used the distance from the anterior-most point of the larvae to the anterior border of the eighth abdominal denticle belt as the full body length. Then, the distance between the anterior-most point of the larvae and the anterior border of the seven other denticle belts was divided by the newly calculated full body length.

*Calculating relative segment position in the absence of anterior-most segments*

In order to calculate relative segment position in the absence of head and thoracic segments (h+t), we removed these segments from our position calculations. Further, to make segment position measurements comparable to when we removed the posterior-most segments (above), we measured segment position and length starting from the posterior-most point of the larvae instead of the anterior-most point. We used the distance from the posterior-most point of the larvae to the anterior border of the first abdominal denticle belt as the full body length. Then the distance from the posterior-most point of the larvae to the anterior border of each of the seven denticle belts was measured and divided by the newly calculated full body length. Notably, we found that taking segment position measurements in the reverse orientation does not change the number and magnitude of position changes between species when compared to measurements taken in the forward orientation.

**Additional effects of “end removal” on observed segment position differences between species**

We examined how the number of significant shifts in relative position is affected from the absence of A8+tail or h+t segments for each individual abdominal segment. We saw that when A8+tail was removed from the relative segment position calculations, both number of significant shifts and average magnitude of shift in segment A7 seemed to be affected the most (~4x less many significant shifts observed as compared to the calculation that included all segments) (Figure S4B,C). When h+t were removed from relative segment position calculations, the number of significant shifts for each segment paralleled the way they were distributed when segment position calculations were made with all segments included, but were slightly higher (Figure S4B). The magnitude of shift appeared to increase towards the posterior end, even more so than they did when calculations were made with all segments included (Figure S4C). These results suggest that changes in the size of A8+tail have strong local effects on segment position whereas changes in the size of h+t have more of a distributed effect along the anterior posterior axis of the larva.

In order to determine if segment positioning in the anterior of the embryo is less noisy as compared to the posterior, we measured coefficient of variation for each segment across species. When all segments are present in position calculations, then the coefficient of variation averaged across all species does not vary considerably across the length of the larva and stays low (~0.03) (Figure S6A). When segment A8+tail or h+t is removed from position calculations then coefficient of variation for the position of nearby segments is reduced. In the case of removal of h+t, coefficient of variation for the position of more posterior segments increased, suggesting that normally h+t may have a stabilizing effect that decreases variation in the position of those segments.

**Differences in the size of A8+tail and h+t between species when all segments are included is segment position calculations**

We also compared the relative size of the head and thorax region (we will refer to this region as h+t), i.e., head and three thoracic segments, normalized to the whole body, among 12 *Drosophila* species (Figure S5B). Relative size of A8+tail is a linear transformation of relative position of segment A8, whereas relative size of h+t is essentially the same as relative position of segment A1.

*D. virilis*, which is the third largest species at the first instar larval stage, had the smallest h+t segments. The biggest h+t segments belonged to *D. persimilis*, the smallest species and to *D. sechellia*, the second largest species at the first instar larval stage. For the rest of the species, relative size of h+t seemed to fall in between, with only few significant differences. Contrarily, we found many significant differences in the relative size of A8+tail segment (Figure S5C). Besides the largest and smallest A8+tail segments observed in *D. mojavensis* and *D. persimilis*, respectively, there are many differences of smaller magnitude between the rest of the species.
