## Supplementary material for "Evolution of Larval Segment Position across 12 *Drosophila* Species": FigS: File_S4.docx

**Supplementary File 4**

**Correlation between changes in the position of adjacent segments along the anterior posterior axis for each species**

We asked if positions were more tightly coordinated between some segments along the anterior posterior axis of the larva as compared to other segments. When each species was considered separately, we found that correlation between the shifts in the positions of adjacent segments was highest and least variable from anterior to posterior in *D. erecta* and *D. mojavensis* as compared to the rest of the species (average correlation coefficients 0.97 and 0.96, respectively) (File S4 - Figures 1, 2). This held true even for segments that were not adjacent. This implies that the local control of segment position is extremely precise along the length of the larvae in these two species. On the other hand, correlation between the shifts in the positions of adjacent segments (except for A1-A2, A2-A3), was lowest for *D. sechellia*. Correlations scores got progressively worse as the segments became further apart.

Some species (*D. simulans*, *D. sechellia*, *D. santomea*) had higher correlations between adjacent segments in the anterior of the larvae (File S4 - Figure 2; segment A7-A8 was left out of these analyses as it was almost always significantly lower and thus biased the analysis), potentially implying stronger coordination of segments in the anterior. In contrast, other species (*D. melanogaster*, *D. yakuba,* *D. virilis*) showed higher correlations in the posterior (File S4 – Figure 2). In a number of species (*D. erecta*, *D. pseudoobscura*, *D. persimilis*, *D. willistoni* and *D. mojavensis*), correlations between adjacent segment pairs were largely unchanged across the larva.

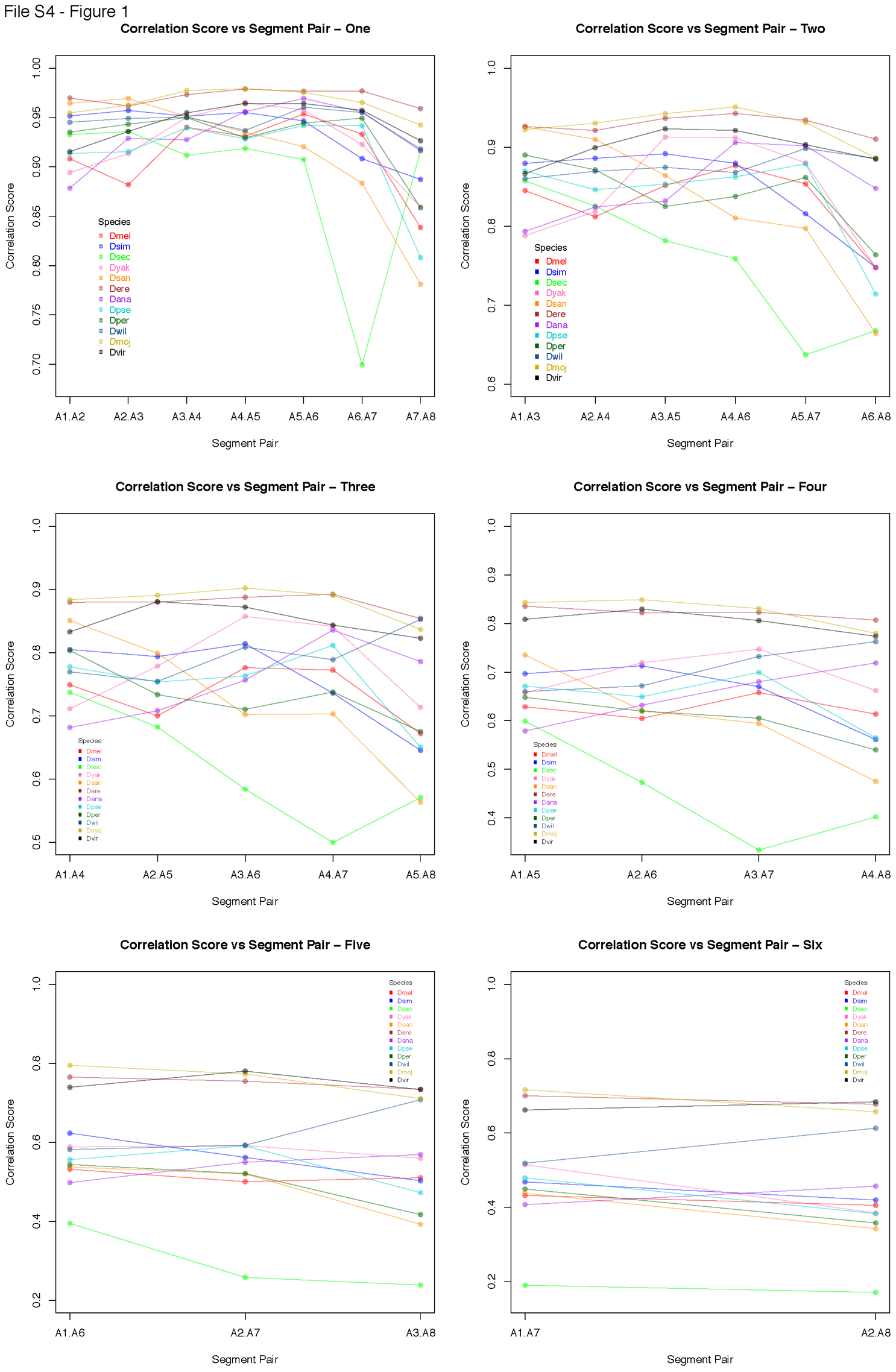

**File S1 – Figure 1.** This series of graphs show how correlation scores change from anterior to posterior in all species. In each graph in the series, segment pairs become further apart from each other (from one to seven segments apart). y axis represents correlation scores and the x axis represents each of the eight abdominal segments. Each species is represented by a different color. Notice, especially as the pairs of segments become further apart, *D. erecta*, *D. mojavensis* and also *D. virilis* stand out as the species with the highest correlation scores.

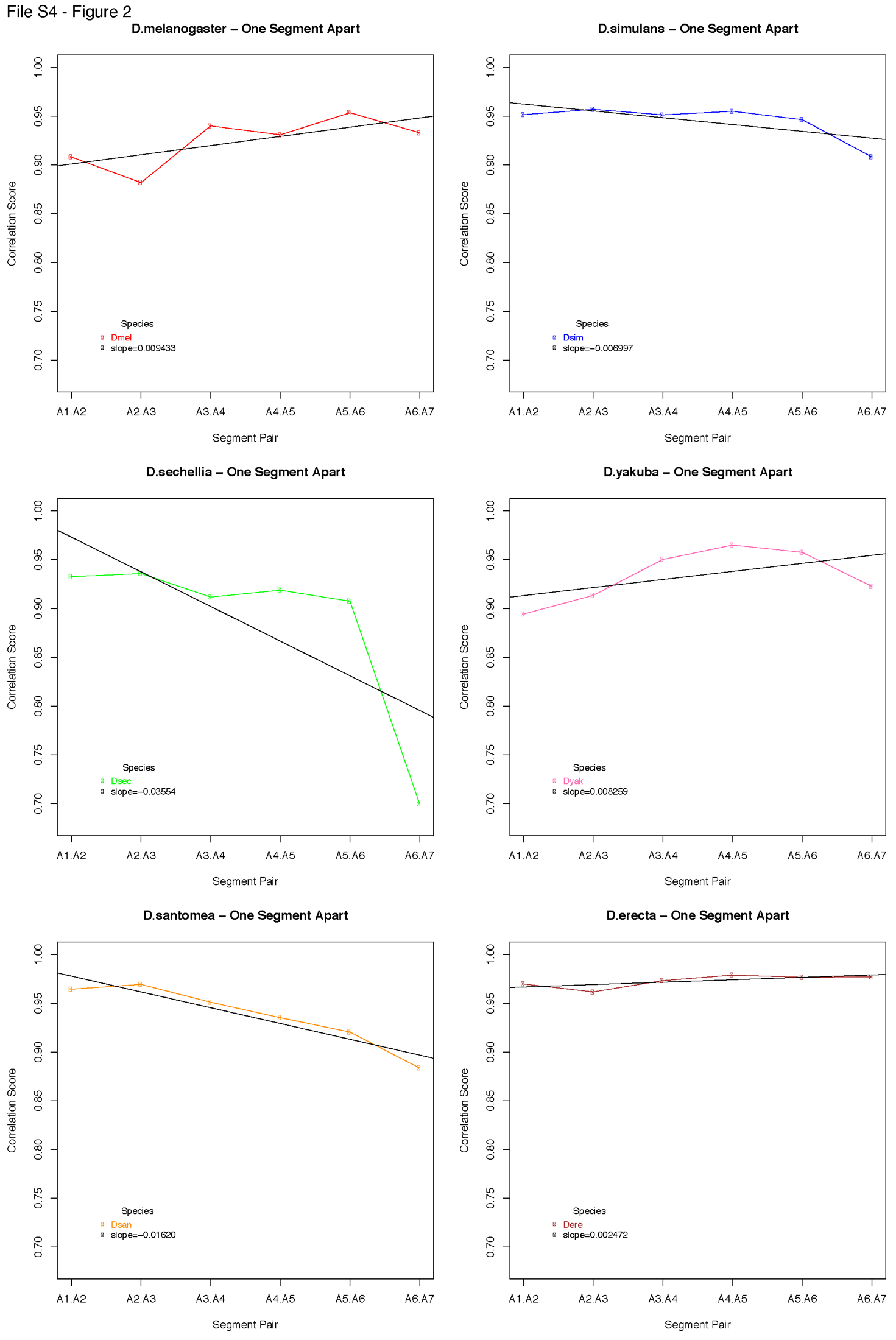

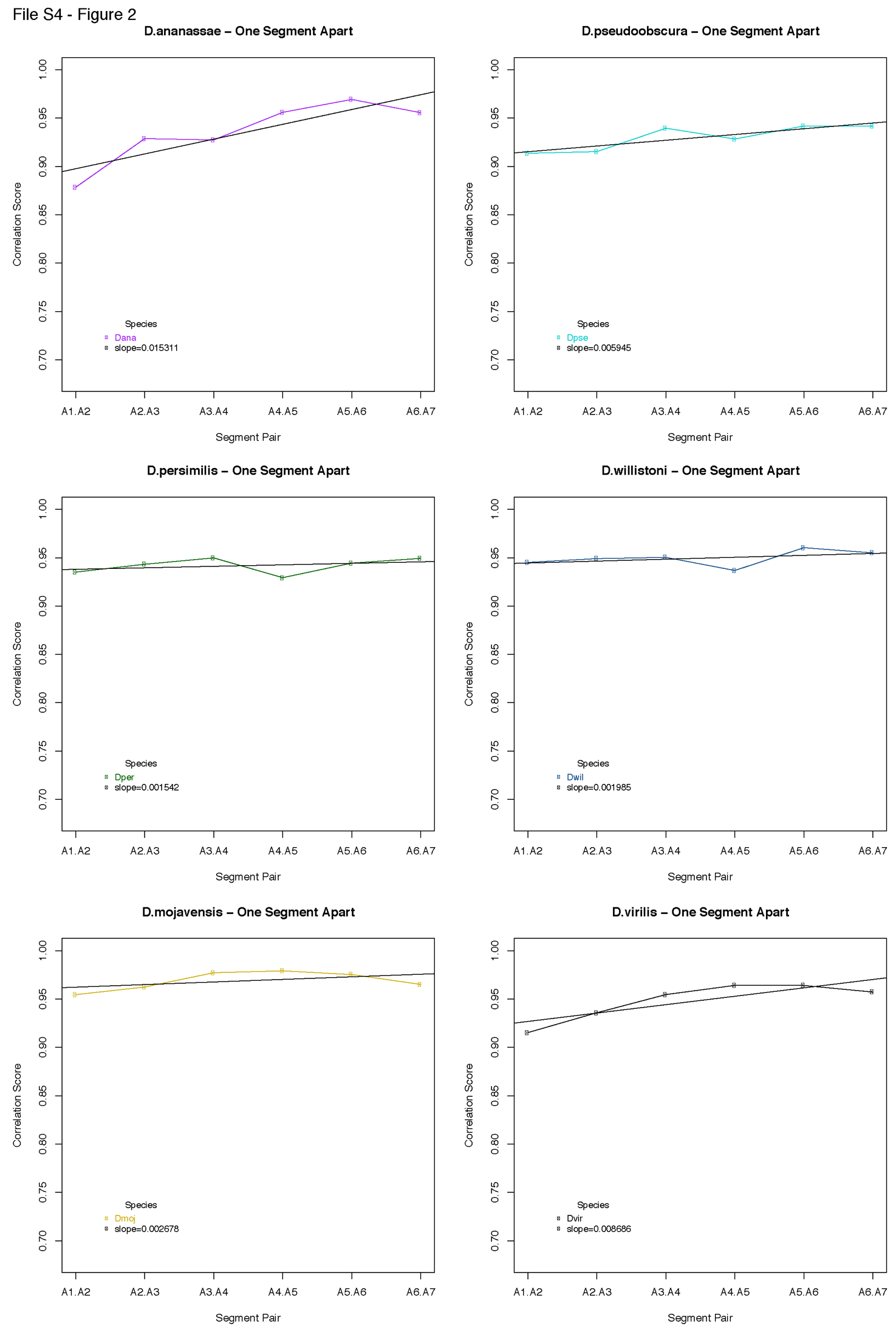

**File S1 – Figure 2.** This series of graphs show how correlation scores change for each species from anterior to posterior. It is essentially a broken down version of the first graph in File S1 – Figure 1, without “A7.A8”. Slope for each trend line is reported in the legend to give an idea of direction of change in correlation scores.
