## Supplementary material for "Evolution of Larval Segment Position across 12 *Drosophila* Species": FigS: Table_S1.docx

| **Table S1. Experimental methods used for number of flies and days needed in a bottle to control population density of each species.** | | | | | |
| --- | --- | --- | --- | --- | --- |
| **Species Name** | **Number of Male and Female flies per bottle** | **Number of days in bottle before removing adults** | **Number of days from egg to larva at 20°C (hours)** | **Number of minutes at 60° oven** | **Number of larval samples from which measurements were taken for data analysis** |
| D. melanogaster (Ore-R) | 15 each | 4-5 days | ~36 | 50 | 145 |
| D. simulans | 15 each | 4-5 days | ~24 | 50 | 129 |
| D. sechellia | 20 each | 7 days | ~36 | 90 | 120 |
| D. yakuba | 10 each | 4-5 days | ~24 | 50 | 143 |
| D. santomea | 25 each | 7 days | ~36 | 50 | 134 |
| D. erecta | 10 each | 4-5 days | ~24 | 50 | 105 |
| D. ananassae | 20 each | 7 days | ~36 | 50 | 107 |
| D. pseudoobscura | 10 each | 7days | ~36 | 50 | 118 |
| D. persimilis | 15 each | 7 days | ~36 | 50 | 113 |
| D. willistoni | 25 each | 7 days | ~48 | 50 | 112 |
| D. mojavensis | 25 each | 7 days | ~72 | 45 | 110 |
| D. virilis | 15 each | 7 days | ~60 | 90 | 113 |
