## Supplementary material for "Evolution of Larval Segment Position across 12 *Drosophila* Species": FigS: Table_S2.docx

| **Supplementary Table S2. Lists the number of segments that are differentially positioned between pairs of species** | | | | |
| --- | --- | --- | --- | --- |
|  | species one | species two | number of segments differentially positioned | divergence time (millions of years) |
| 1 | Dana | Dere | 8 | 15 |
| 2 | Dana | Dmel | 5 | 15 |
| 3 | Dana | Dmoj | 7 | 32 |
| 4 | Dana | Dper | 8 | 24 |
| 5 | Dana | Dpse | 4 | 24 |
| 6 | Dana | Dsan | 3 | 15 |
| 7 | Dana | Dsec | 8 | 15 |
| 8 | Dana | Dsim | 0 | 15 |
| 9 | Dana | Dvir | 6 | 32 |
| 10 | Dana | Dwil | 0 | 32 |
| 11 | Dana | Dyak | 7 | 15 |
| 12 | Dere | Dmel | 5 | 3.4 |
| 13 | Dere | Dmoj | 5 | 32 |
| 14 | Dere | Dper | 8 | 24 |
| 15 | Dere | Dpse | 7 | 24 |
| 16 | Dere | Dsan | 7 | 2.7 |
| 17 | Dere | Dsec | 8 | 3.4 |
| 18 | Dere | Dsim | 8 | 3.4 |
| 19 | Dere | Dvir | 6 | 32 |
| 20 | Dere | Dwil | 8 | 32 |
| 21 | Dere | Dyak | 8 | 32 |
| 22 | Dmel | Dmoj | 7 | 32 |
| 23 | Dmel | Dper | 8 | 24 |
| 24 | Dmel | Dpse | 7 | 24 |
| 25 | Dmel | Dsan | 6 | 3.4 |
| 26 | Dmel | Dsec | 7 | 1.4 |
| 27 | Dmel | Dsim | 5 | 1.4 |
| 28 | Dmel | Dvir | 2 | 32 |
| 29 | Dmel | Dwil | 7 | 32 |
| 30 | Dmel | Dyak | 6 | 3.4 |
| 31 | Dmoj | Dper | 8 | 32 |
| 32 | Dmoj | Dpse | 7 | 32 |
| 33 | Dmoj | Dsan | 7 | 32 |
| 34 | Dmoj | Dsec | 8 | 32 |
| 35 | Dmoj | Dsim | 8 | 32 |
| 36 | Dmoj | Dvir | 7 | 10 |
| 37 | Dmoj | Dwil | 7 | 27 |
| 38 | Dmoj | Dyak | 7 | 32 |
| 39 | Dper | Dpse | 8 | 0.6 |
| 40 | Dper | Dsan | 8 | 24 |
| 41 | Dper | Dsec | 7 | 24 |
| 42 | Dper | Dsim | 8 | 24 |
| 43 | Dper | Dvir | 8 | 32 |
| 44 | Dper | Dwil | 8 | 32 |
| 45 | Dper | Dyak | 8 | 24 |
| 46 | Dpse | Dsan | 6 | 24 |
| 47 | Dpse | Dsec | 4 | 24 |
| 48 | Dpse | Dsim | 7 | 24 |
| 49 | Dpse | Dvir | 6 | 32 |
| 50 | Dpse | Dwil | 3 | 32 |
| 51 | Dpse | Dyak | 6 | 24 |
| 52 | Dsan | Dsec | 8 | 3.4 |
| 53 | Dsan | Dsim | 5 | 3.4 |
| 54 | Dsan | Dvir | 6 | 32 |
| 55 | Dsan | Dwil | 3 | 32 |
| 56 | Dsan | Dyak | 1 | 0.4 |
| 57 | Dsec | Dsim | 8 | 0.25 |
| 58 | Dsec | Dvir | 5 | 32 |
| 59 | Dsec | Dwil | 7 | 32 |
| 60 | Dsec | Dyak | 8 | 3.4 |
| 61 | Dsim | Dvir | 5 | 32 |
| 62 | Dsim | Dwil | 4 | 32 |
| 63 | Dsim | Dyak | 6 | 3.4 |
| 64 | Dvir | Dwil | 6 | 27 |
| 65 | Dvir | Dyak | 6 | 32 |
| 66 | Dwil | Dyak | 5 | 32 |
