## Supplementary material for "Evolution of Larval Segment Position across 12 *Drosophila* Species": FigS: Table_S3.docx

| **Supplementary Table 3. Lists the deviation of the position of each segment in each species from the “across-species” mean.** | | | | | |
| --- | --- | --- | --- | --- | --- |
| species | segment | deviation from the species mean | absolute deviation from the species mean | total (absolute) deviation of mean over all segments | In of all species pair comparisons, number of significant segment position changes for each species |
| Dana | A1 | 0.1568381 | 0.1568381 | 1.74198203 | 56 (13.66%) |
| Dana | A2 | 0.23681895 | 0.23681895 |  |  |
| Dana | A3 | 0.25958437 | 0.25958437 |  |  |
| Dana | A4 | 0.22392065 | 0.22392065 |  |  |
| Dana | A5 | 0.18130454 | 0.18130454 |  |  |
| Dana | A6 | 0.22764646 | 0.22764646 |  |  |
| Dana | A7 | 0.15954071 | 0.15954071 |  |  |
| Dana | A8 | 0.29632824 | 0.29632824 |  |  |
| Dere | A1 | -0.4150311 | 0.41503106 | 8.86873982 | 78 (19.02%) |
| Dere | A2 | -0.5967867 | 0.59678675 |  |  |
| Dere | A3 | -0.8037267 | 0.80372671 |  |  |
| Dere | A4 | -1.0135141 | 1.01351415 |  |  |
| Dere | A5 | -1.1776149 | 1.17761491 |  |  |
| Dere | A6 | -1.4161008 | 1.41610076 |  |  |
| Dere | A7 | -1.7061228 | 1.70612284 |  |  |
| Dere | A8 | -1.7398427 | 1.73984265 |  |  |
| Dmel | A1 | -0.6439966 | 0.64399657 | 3.66891602 | 65 (15.85%) |
| Dmel | A2 | -0.6237096 | 0.62370957 |  |  |
| Dmel | A3 | -0.6214804 | 0.6214804 |  |  |
| Dmel | A4 | -0.3758557 | 0.37585569 |  |  |
| Dmel | A5 | -0.2248037 | 0.22480374 |  |  |
| Dmel | A6 | 0.00472026 | 0.00472026 |  |  |
| Dmel | A7 | 0.51753233 | 0.51753233 |  |  |
| Dmel | A8 | 0.65681745 | 0.65681745 |  |  |
| Dmoj | A1 | -0.0417583 | 0.04175833 | 18.4652117 | 78 (19.02%) |
| Dmoj | A2 | -0.7664274 | 0.76642744 |  |  |
| Dmoj | A3 | -1.2101293 | 1.21012931 |  |  |
| Dmoj | A4 | -1.8972977 | 1.8972977 |  |  |
| Dmoj | A5 | -2.5590608 | 2.55906079 |  |  |
| Dmoj | A6 | -3.28717 | 3.28717002 |  |  |
| Dmoj | A7 | -3.9781228 | 3.97812284 |  |  |
| Dmoj | A8 | -4.7252452 | 4.72524525 |  |  |
| Dper | A1 | 0.94027868 | 0.94027868 | 18.2363837 | 87 (21.22%) |
| Dper | A2 | 1.50173748 | 1.50173748 |  |  |
| Dper | A3 | 1.87890288 | 1.87890288 |  |  |
| Dper | A4 | 2.15942475 | 2.15942475 |  |  |
| Dper | A5 | 2.54855787 | 2.54855787 |  |  |
| Dper | A6 | 2.72579136 | 2.72579136 |  |  |
| Dper | A7 | 3.02886831 | 3.02886831 |  |  |
| Dper | A8 | 3.45282233 | 3.45282233 |  |  |
| Dpse | A1 | -0.0783361 | 0.07833614 | 4.61316168 | 65 (15.85%) |
| Dpse | A2 | 0.06823666 | 0.06823666 |  |  |
| Dpse | A3 | 0.3435009 | 0.3435009 |  |  |
| Dpse | A4 | 0.5554342 | 0.5554342 |  |  |
| Dpse | A5 | 0.67677896 | 0.67677896 |  |  |
| Dpse | A6 | 0.8069178 | 0.8069178 |  |  |
| Dpse | A7 | 1.01891106 | 1.01891106 |  |  |
| Dpse | A8 | 1.06504597 | 1.06504597 |  |  |
| Dsan | A1 | -0.0797325 | 0.07973255 | 2.00922525 | 60 (14.63%) |
| Dsan | A2 | -0.0622402 | 0.0622402 |  |  |
| Dsan | A3 | -0.0621062 | 0.06210624 |  |  |
| Dsan | A4 | -0.1140358 | 0.11403583 |  |  |
| Dsan | A5 | -0.1653498 | 0.1653498 |  |  |
| Dsan | A6 | -0.3448783 | 0.3448783 |  |  |
| Dsan | A7 | -0.4844661 | 0.48446613 |  |  |
| Dsan | A8 | -0.6964162 | 0.69641621 |  |  |
| Dsec | A1 | 1.03255228 | 1.03255228 | 7.58229589 | 78 (19.02%) |
| Dsec | A2 | 1.03628468 | 1.03628468 |  |  |
| Dsec | A3 | 0.93766615 | 0.93766615 |  |  |
| Dsec | A4 | 0.88333109 | 0.88333109 |  |  |
| Dsec | A5 | 0.88232557 | 0.88232557 |  |  |
| Dsec | A6 | 0.79912543 | 0.79912543 |  |  |
| Dsec | A7 | 0.98804382 | 0.98804382 |  |  |
| Dsec | A8 | 1.02296687 | 1.02296687 |  |  |
| Dsim | A1 | 0.32465887 | 0.32465887 | 1.76293065 | 64 (15.61%) |
| Dsim | A2 | 0.02282344 | 0.02282344 |  |  |
| Dsim | A3 | -0.0737156 | 0.07371563 |  |  |
| Dsim | A4 | 0.0538214 | 0.0538214 |  |  |
| Dsim | A5 | 0.22511821 | 0.22511821 |  |  |
| Dsim | A6 | 0.29352272 | 0.29352272 |  |  |
| Dsim | A7 | 0.35652832 | 0.35652832 |  |  |
| Dsim | A8 | 0.41274207 | 0.41274207 |  |  |
| Dvir | A1 | -1.1391903 | 1.13919035 | 5.17973079 | 63 (15.36%) |
| Dvir | A2 | -0.8658731 | 0.86587314 |  |  |
| Dvir | A3 | -0.7026016 | 0.70260155 |  |  |
| Dvir | A4 | -0.4642036 | 0.46420357 |  |  |
| Dvir | A5 | -0.1616191 | 0.16161912 |  |  |
| Dvir | A6 | 0.41118959 | 0.41118959 |  |  |
| Dvir | A7 | 0.66408955 | 0.66408955 |  |  |
| Dvir | A8 | 0.77096392 | 0.77096392 |  |  |
| Dwil | A1 | 0.06595109 | 0.06595109 | 2.19713164 | 58 (14.15%) |
| Dwil | A2 | 0.2999573 | 0.2999573 |  |  |
| Dwil | A3 | 0.52305901 | 0.52305901 |  |  |
| Dwil | A4 | 0.48764061 | 0.48764061 |  |  |
| Dwil | A5 | 0.27636128 | 0.27636128 |  |  |
| Dwil | A6 | 0.19125043 | 0.19125043 |  |  |
| Dwil | A7 | -0.1766943 | 0.17669427 |  |  |
| Dwil | A8 | -0.1762177 | 0.17621765 |  |  |
| Dyak | A1 | -0.0419541 | 0.04195413 | 3.27637951 | 68 (16.58%) |
| Dyak | A2 | -0.1425883 | 0.14258828 |  |  |
| Dyak | A3 | -0.3278426 | 0.32784259 |  |  |
| Dyak | A4 | -0.4455774 | 0.44557742 |  |  |
| Dyak | A5 | -0.5241657 | 0.52416569 |  |  |
| Dyak | A6 | -0.5134567 | 0.51345674 |  |  |
| Dyak | A7 | -0.6486823 | 0.64868228 |  |  |
| Dyak | A8 | -0.6321124 | 0.63211238 |  |  |
