## Supplementary figures and images for "Evolution of Larval Segment Position across 12 *Drosophila* Species"

### Figure_S13_corrscoresantpost.pdf

Supplementary Figure 13

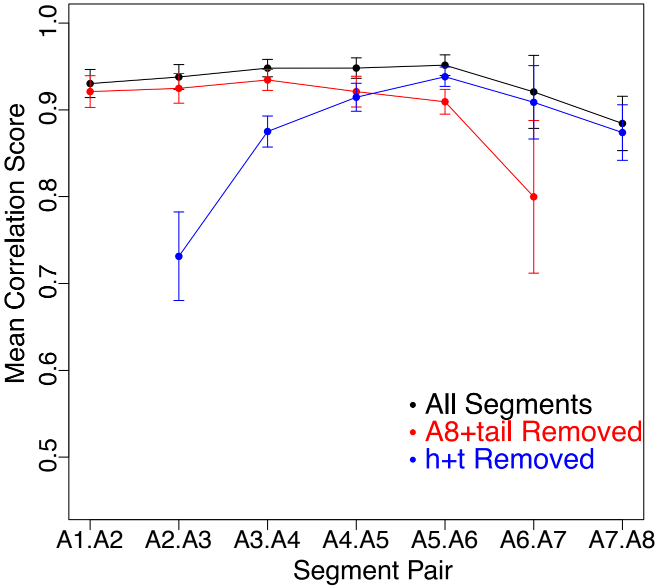
